# Pharmacogenomic screening identifies and repurposes leucovorin and dyclonine as pro-oligodendrogenic compounds in brain repair

**DOI:** 10.1101/2024.08.08.607135

**Authors:** Jean-Baptiste Huré, Louis Foucault, Litsa Maria Ghayad, Corentine Marie, Nicolas Vachoud, Lucas Baudouin, Rihab Azmani, Natalija Ivjanin, Alvaro Arevalo-Nuevo, Morgane Pigache, Lamia Bouslama-Oueghlani, Julie-Anne Chemelle, Marie-Aimée Dronne, Raphaël Terreux, Bassem Hassan, François Gueyffier, Olivier Raineteau, Carlos Parras

## Abstract

Oligodendrocytes are the myelin-forming cells of the central nervous system (CNS), with oligodendroglial pathologies leading to strong disabilities, from early preterm-birth brain injury (PBI) to adult multiple sclerosis (MS). No medication presenting convincing repair capacity in humans has been approved for these pathologies so far. Here, we present a pharmacogenomic approach leading to the identification of small bioactive molecules with a large pro-oligodendrogenic activity, selected through an expert curation scoring strategy (OligoScore) of their large impact on transcriptional programs controlling oligodendrogenesis and (re)myelination. We demonstrate the pro-oligodendrogenic activity of these compounds *in vitro,* using neural and oligodendrocyte progenitor cell (OPC) cultures, as well as *ex vivo,* using organotypic cerebellar explant cultures. Focusing on the two most promising molecules, i.e. leucovorin and dyclonine, we tested their therapeutic efficacy using a mouse model of neonatal chronic hypoxia, which faithfully mimics aspects of PBI. In this model, both compounds promoted proliferation and oligodendroglial fate acquisition from neural stem/progenitor cells, with leucovorin also promoting their differentiation. We extended these findings to an adult focal de/remyelination mouse MS model, in which both compounds improved lesion repair by promoting OPC differentiation while maintaining the pool of OPCs, and in parallel, by accelerating the transition from pro-inflammatory to pro-regenerative microglial profiles and myelin debris clearance. This study paves the way for clinical trials aimed at repurposing these FDA-approved compounds to treat myelin pathologies such as PBI and MS.

**Graphical abstract:** 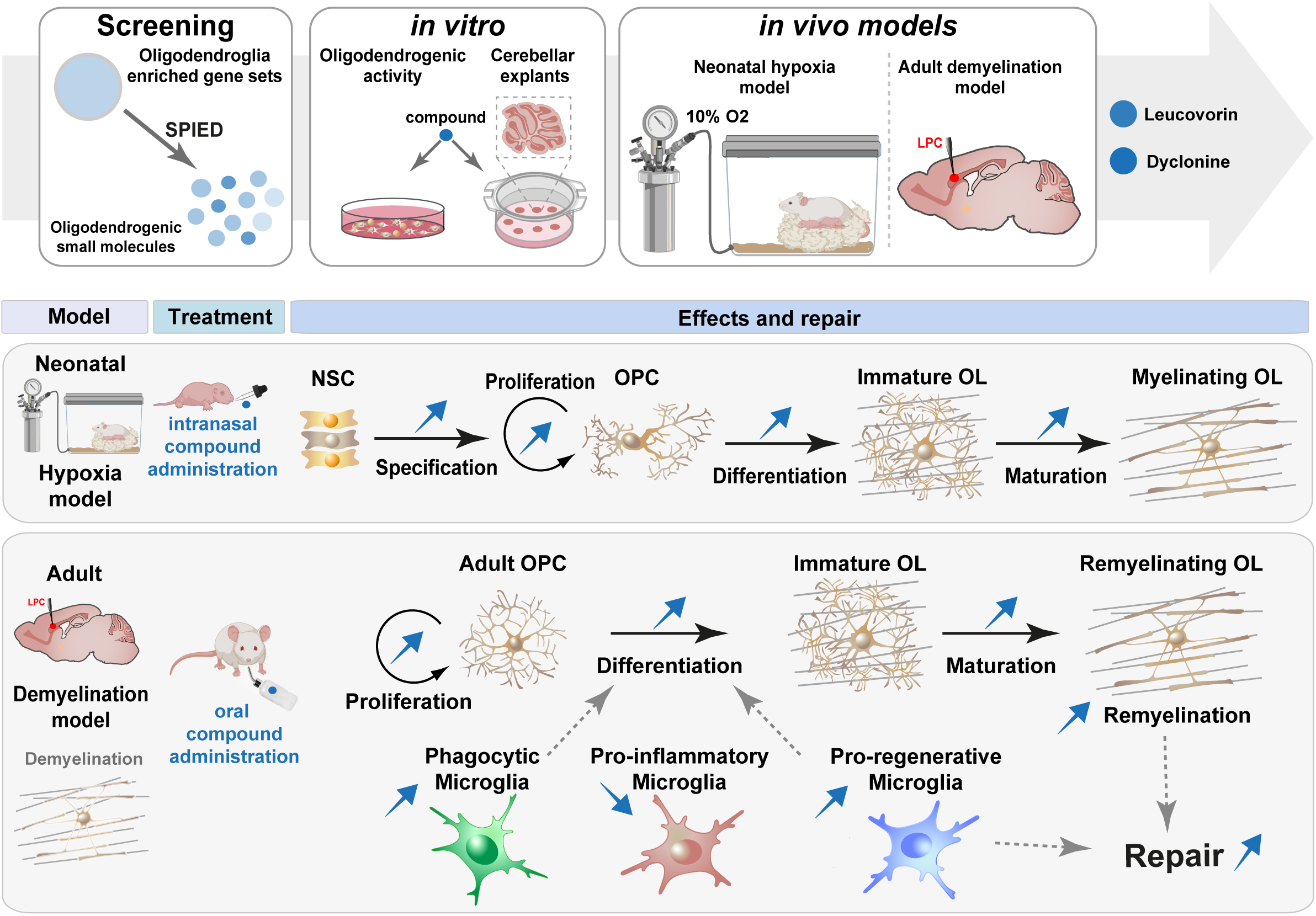

**Pharmacogenomic screening identifies and repurposes dyclonine and leucovorin as pro-oligodendrogenic and pro-myelinating compounds.** Schematics of the pharmacogenomic approach leading to the identification of small bioactive molecules (compounds) with potential pro-oligodendrogenic activity, followed by the *in vitro* validation of the top compounds using neural and oligodendrocyte progenitor cell (OPC) cultures as well as organotypic cerebellar explants. The therapeutic efficacy of the top two compounds, leucovorin and dyclonine, both approved by the Food and Drug Administration (FDA), was assessed *in vivo* using two clinically relevant mouse models of myelin pathologies. In the neonatal hypoxia mouse model, mimicking some aspects of preterm brain injury, both leucovorin and dyclonine promoted neural stem cell (NSC) differentiation into OPCs and OPC proliferation, with leucovorin additionally restoring the density of myelinating OLs found in normoxic conditions. In an adult focal de/remyelination mouse model of multiple sclerosis, both compounds significantly improved lesion repair in adult mice by promoting OPC differentiation while preserving the pool of OPCs, and by accelerating myelin debris clearance and shifting microglia from pro-inflammatory to pro-regenerative profiles.

## Introduction

Preterm birth brain injury (PBI) is the most common cause of death and disability in children under 5 years, affecting 15 million infants yearly born before 37 gestational weeks. Rates of PBI are increasing in developing countries (e.g., 7% in France and the UK, 13% in the US), and are associated with a high level of morbidity and neonatal encephalopathies, leading to persistent cognitive and neuropsychiatric deficits (such as autism spectrum, attention-deficit disorders, and epilepsy) (Harrison and Goldenberg, 2016; Luu et al., 2017; Walani, 2020). Alterations of oxygen concentrations and perinatal neuroinflammation of PBI dramatically affect the survival and maturation of oligodendrocyte precursor cells (OPCs) (Back et al., 2001; Volpe, 2009), resulting in hypomyelination, abnormal connectivity, and synaptopathies. Indeed, in both humans and rodents, the perinatal period is a time of active oligodendrogenesis, myelination, and axonal growth within the developing subcortical white matter (Fletcher et al., 2021). Given the severity of long-term consequences and the absence of current treatment, there is an urgent need to develop novel therapeutic approaches for promoting perinatal brain repair through oligodendrocyte (OL) regeneration.

Another myelin pathology affecting many adults (3 million people) is multiple sclerosis (MS), a chronic autoimmune-mediated disease characterized by focal demyelinated lesions of the central nervous system (CNS) (Lassmann, 2018; Reich et al., 2018). Although current immunomodulatory therapies can reduce the frequency and severity of relapses, they show limited impact on disease progression and are inefficient in lesion repair (Hauser and Cree, 2020). Therefore, complementary strategies to protect OLs and induce the generation of new remyelinating OLs are urgently needed to promote repair and eventually end and even reverse MS progression. Accordingly, animal models have demonstrated that remyelination can reestablish fast saltatory conduction velocity (Smith et al., 1979), prevent axonal loss, and exert neuroprotection (Irvine and Blakemore, 2008), resulting in functional clinical recovery (Duncan et al., 2009; Jeffery and Blakemore, 1997). Brain remyelination is thought to result mainly from the differentiation of parenchymal OPCs, in regions close to the lateral ventricles, also from adult SVZ-NSC-derived OPCs (Samanta et al., 2015; Xing et al., 2014), and to a lesser extent from mature OLs (Bacmeister et al., 2020; Duncan et al., 2018; Mezydlo et al., 2023; Yeung et al., 2019). The incomplete remyelination in MS patients (Bodini et al., 2016) is attributed to a variety of oligodendroglia-associated intrinsic as well as lesion-environment-related extrinsic factors (Galloway et al., 2020), and the extent of remyelination differs considerably between individual patients and lesion location, declining with disease progression (Patrikios et al., 2006). Despite some factors and small molecules (compounds) being shown to improve myelination in cell cultures, as well as remyelination in animal models, no medication presenting convincing remyelinating capacity in humans has been approved for MS patients so far, with clemastine and analogs of thyroid hormone still having to complete phase III clinical trials (Liu et al., 2016; Mei et al., 2014; Rodriguez-Pena, 1999; Schoonover et al., 2004). Thus, enhancing (re)myelination remains an unmet medical need both in the context of preterm birth brain injury and MS patients.

Owing to their diverse etiologies, onset ages, and potential repair mechanisms, the discovery of new treatments for treating myelin pathologies should aim at globally stimulating endogenous oligodendrogenesis, known to spontaneously increase following injury, at multiple levels, i.e. from the specification of new OPCs from neural progenitor cells (NPCs), to the maturation of pre-existing ones within the parenchyma. A promising drug discovery strategy is to use connectivity maps to identify compounds able to induce transcriptional signatures similar to those observed in a given cell lineage. This approach lies behind the connectivity-map initiative, which has been extensively used in various tissues [reviewed in (Lamb, 2007)]. This strategy has resulted in successful drug repurposing in several contexts [e.g., dietary restriction (Calvert et al., 2016); aging and oxidative stress (Ye et al., 2014; Ziehm et al., 2017)], also allowing to discover new modes of action for compounds, as well as predicting off-target or side effects.

While most previous studies have followed a gene/pathway candidate approach, we aimed here at using a more comprehensive strategy (Fig. 1 extended data 1). To this end, we developed an *in-silico* approach combining both unbiased identification of transcriptional signatures associated with oligodendroglial formation (i.e., transcriptomes) and knowledge-driven curation of transcriptional changes associated with oligodendrogenesis (i.e., OligoScore). We then conducted a pharmacogenomic analysis to identify small bioactive molecules (compounds) capable of globally mimicking this transcriptional signature. Using a scoring strategy based on genes involved in oligodendrogenesis as well as pharmacogenetics/pharmacokinetics criteria, we ranked and selected the most promising compounds. Finally, after validating their pro-oligodendrogenic activity *in vitro* and *ex vivo*, we selected the top two compounds, leucovorin and dyclonine, and demonstrated their capacity to promote oligodendrogenesis and myelin repair *in vivo*, using both neonatal and adult murine models of myelin pathologies.

**Figure 1.**
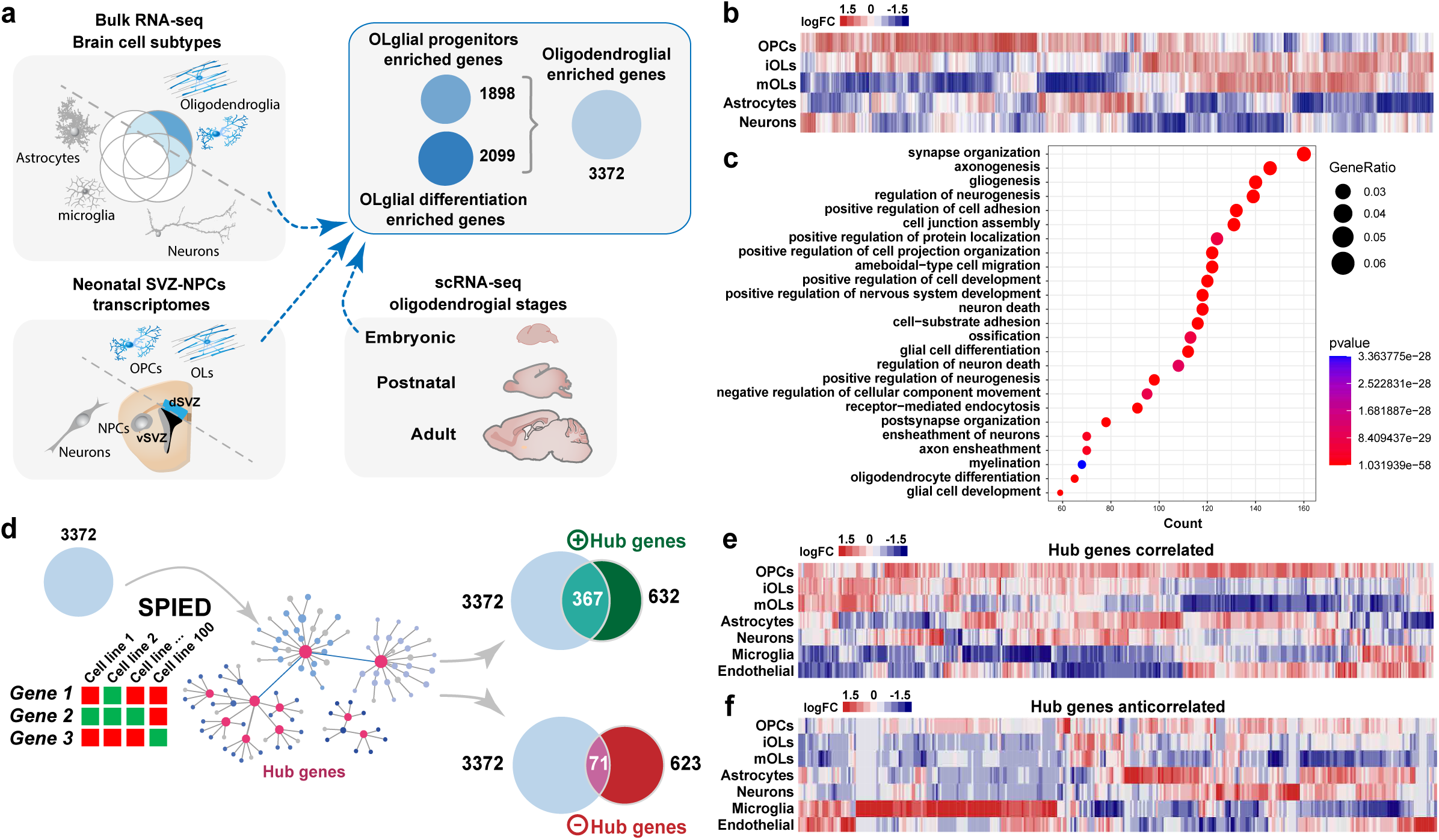
In-silico strategy to generate oligodendroglial-enriched gene sets and their use in pharmacogenomics to identify new hub genes. **(a)** Use of both bulk and single-cell RNA-seq datasets from different brain cell types and oligodendroglial stages, together with transcriptomic comparison of dorsal (gliogenic) vs. ventral (neurogenic) neonatal forebrain progenitors to identify gene sets enriched either in progenitors, more mature cells, or both, depicted as circles indicating the number of genes. **(b)** Heat map showing the differential expression of the 3372 gene set in different neural cell types. **(c)** Top gene ontology biological processes related to the 3372 genes. **(d)** Schematics representing the interrogation of the SPIED platform with the 3372 genes to identify hub genes correlated (positive hub genes) or anti-correlated (negative hub genes). Note that most of the positively correlated hub genes are included in the 3372 gene set, whereas only a few of the anti-correlated ones. (e, f) Heatmap visualization of the expression of correlated- **(e)** and anticorrelated-hub genes **(f)** in different brain cells indicates that most of the correlated genes are expressed in oligodendroglia while many anticorrelated hub genes are enriched in microglia, astrocytes, and endothelial cells.

## Results

### Generation of a transcriptional signature and associated “hub genes” defining oligodendrogenesis

We first set out to generate a broad transcriptional signature of oligodendrocyte lineage cells (Supplementary Table 1) by comparing the transcriptome of these cells at different developmental stages, using recently published single-cell- and bulk-transcriptomic datasets from mouse brain cells (Falcão et al., 2018; Jäkel et al., 2019; Marques et al., 2018; Marques et al., 2016; Saunders et al., 2018; Zhang et al., 2014). Given that oligodendroglial cells are robustly generated in mice from the dorsal aspect of the ventricular-subventricular zone (V-SVZ) at neonatal stages (Kessaris et al., 2006; Suzuki and Goldman, 2003), we integrated the transcriptional signature obtained by comparing dorsal *vs*. lateral neonatal subventricular progenitors (Supplementary Table 2) (Azim et al., 2017; Azim et al., 2015) to include the transcriptional changes contributing to OL specification from NPCs. Using this strategy, we obtained oligodendroglial transcriptional signatures (Supplementary Table 3) enriched either in progenitor cells (1898 genes) or in more mature stages (2099 genes), which we refined to 3372 genes (Supplementary Table 4) using further selection criteria (Methods). This oligodendroglial signature was indeed enriched in oligodendroglia compared to other neural cells, as illustrated by its expression profile in postnatal brain cells (Fig. 1b) and its enrichment in gene ontology (GO) terms related to OL differentiation, gliogenesis, and glial cell differentiation (Fig. 1c, Supplementary Table 4), together with processes linked to neuronal development, synaptic and dendritic organization, known to be modulated by oligodendroglial-dependent brain plasticity and adaptive myelination (Buchanan et al., 2021; Foster et al., 2019; Pease-Raissi and Chan, 2021; Xiao et al., 2022).

To identify “hub genes” involved in oligodendrogenesis, we used our oligodendroglial signature to query the SPIED platform (Fig. 1d), which provides a fast and simple quantitative interrogation of publicly available gene expression datasets (Williams, 2013). This analysis resulted in the identification of hub genes correlated (632 genes) and anticorrelated (623 genes) with our oligodendroglial transcriptional signature (Supplementary Tables 5 and 6). Correlated hub genes contained numerous key regulators of oligodendrogenesis (including *Olig1, Olig2, Sox8, Sox10, Nkx2-2)*, as well as commonly used oligodendroglial markers (such as *Cnp, Mag, Mbp, Mobp, Mog, Opalin, Plp1;* Supplementary Table 5). Their expression pattern was enriched in oligodendroglia compared to other brain cells (Fig. 1e) and their associated biological processes were enriched in gliogenesis, CNS myelination, and OL differentiation (Supplementary Table 5), thus validating the efficiency of our strategy. Notably, while many correlated hub genes (367/632 genes) were included in our oligodendroglial signature, only a few anti-correlated ones (71/623) were present (Fig. 1e). Interestingly, a large fraction of anti-correlated hub genes (150/632) was expressed in microglia (Fig. 1f) and enriched in GO terms associated with inflammation and immune cell activation (such as IL12, IFNγ or ROS production; Supplementary Table 6), known to inhibit OL generation. Altogether, these results indicate that this methodology constitutes an efficient approach to identify a large oligodendroglial transcriptional signature, and to infer hub genes positively or negatively associated with oligodendrogenesis.

### Identification and expert curation of compounds acting on oligodendroglial-associated genes

We then queried SPIED with our oligodendroglial signature to obtain small bioactive molecules (compounds) inducing similar transcriptional changes and found a large number (449) of positively associated compounds. To rank them by the impact of genes involved in oligodendrogenesis, we introduced a knowledge-driven scoring procedure that we named OligoScore (here provided for the community as a resource; Fig. 2 extended data 1; https://oligoscore.icm-institute.org). This curation strategy included more than 430 genes for which loss-of-function and gain-of-function studies have demonstrated their requirement in the main processes of oligodendrogenesis (that we categorized in specification, proliferation, migration, survival, differentiation, myelination, and remyelination). Genes were scored in each process from 1 to 3 (low, medium, strong) either positively or negatively (promoting or inhibiting, respectively), depending on the severity of gain- or loss-of-function phenotypes. For example, *Olig2* and *Sox10*, key regulators of oligodendrogenesis (Lu et al., 2002; Stolt et al., 2002; Stolt and Wegner, 2010; Yu et al., 2013), have the highest positive values in specification and differentiation, respectively, while *Tnf* (Minchenberg et al., 2015; Nakazawa et al., 2006) or *Tlr2* (Sloane et al., 2010) have negative values in processes such as survival and differentiation (Fig. 2a). We validated the efficacy of the OligoScore strategy by querying it with genes differentially expressed in OPCs upon either a genetic (*Chd7* specific deletion in OPCs, (Marie et al., 2018)) or environmental perturbation (systemic injection of the pro-inflammatory cytokine IL1β, (Schang et al., 2022)). Our analysis confirmed that OligoScore accurately predicted the deregulation of oligodendrogenesis-related processes identified in these studies (Fig. 2 extended data 2, and Methods). Then, using these curated gene sets involved in oligodendrogenesis (Fig. 2b), we interrogated SPIED, finding a large number (393) of positively associated compounds, with 156 in common with those obtained using our broad oligodendroglial signature (Fig. 2c). Focusing on these common 156 compounds, we next used OligoScore to rank them by their total scores (sum of each process’ score, defined as pharmacogenomic score) thus selecting those with broader pro-oligodendrogenic activities (Fig. 2d-f; Supplementary Table 7). Validating the pertinence of our strategy, among the top-ranked compounds, we found molecules reported to promote oligodendrogenesis, including liothyronine (thyroid hormone), clemastine (Green et al., 2017; Mei *et al*., 2014), clomipramine (Faissner et al., 2017), piperidolate (Abiraman et al., 2015), luteolin (Barbierato et al., 2015), ifenprodil (Hubler et al., 2018), and propafenone (Najm et al., 2015). For further analyses, we selected the top 40 compounds ranked by their pharmacogenomic scores and therefore by their broader putative pro-oligodendrogenic activities, excluding those already known for their implication in oligodendrogenesis. After studying their pharmacological properties (summarized in Methods Table 8), we then selected the top 11 compounds (Sm1-Sm11; Methods Table 1) for their pharmacogenomic scores (Fig. 2d), ability to cross the BBB, and low potential toxicity (Fig. 2g, h).

**Figure 2.**
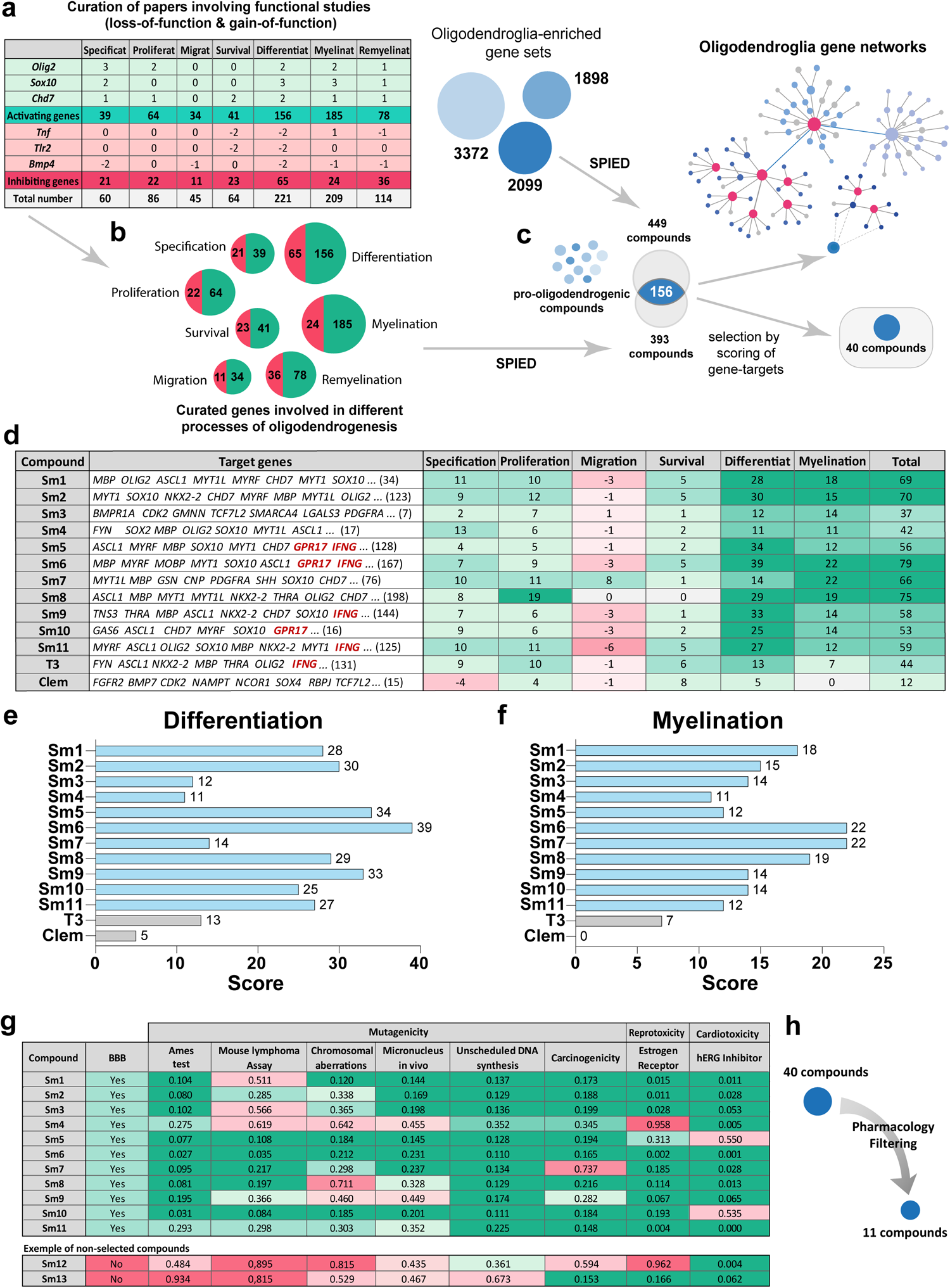
Expert curation ranking and selection of small molecules. (**a**) Table illustrating the scoring of selected genes for their regulation in different oligodendrogenesis processes (positive in green or negative in red), based on bibliographic curation of functional studies. (**b**) Curated genes subsets promoting (green) or inhibiting (red) each oligodendrogenesis process. (**c**) Using both curated and oligodendroglial-enriched genes sets to interrogate the SPIED platform and generate a list of small molecules /compounds regulating their expression. Shared compounds were then ranked by the scoring of their curated gene signatures, in order to select 40 compounds with novel pro-oligodendrogenic transcriptional activity. (**d**) Table listing the top 11 small molecules (Sm) alongside two positive controls (T3 and clemastine) exemplifying some of their curated gene targets and their corresponding scoring values (green: positive, red: negative) for each oligodendrogenesis process. (**e**, **f**) Barplots illustrating the score of selected compounds in the differentiation (**e**) and myelination (**f**) processes, respectively. (**g**) Table and (**h**) schematics illustrating the pharmacological filtering criteria (BBB permeability, mutagenicity, reprotoxicity, cardiotoxicity, etc.) used to select the top 11 compounds. Green to red gradient highlights the probability values from positive to negative activity on each pharmacological parameter. BBB: Blood-Brain-Barrier, hERG: ether-a-go-go related gene potassium channel.

### Pro-oligodendrogenic activity of selected compounds in neonatal neural progenitor cells

To evaluate the activity of selected pro-oligodendrogenic compounds in the context of different neural cell types, we first used mixed neural progenitor cell (NPC) cultures obtained from the neonatal V-SVZ (postnatal day 0 to 1, P0-P1) and amplified using the neurosphere protocol (Parras et al., 2004; Reynolds and Weiss, 1992) (Fig. 3a). We assessed the dose effect of selected compounds to promote NPC differentiation into So×10^+^ oligodendroglia using three different concentrations (250nM, 500nM, and 750nM) based on previous reports (Eleuteri et al., 2017; Mei *et al*., 2014; Najm *et al*., 2015). Automatic quantification of So×10^+^ cells indicated that all selected compounds presented the strongest effects to promote So×10^+^ oligodendroglia at 750nM (Fig. 3 extended data 1). We thus used this concentration for the following *in vitro* studies. To assess the capacity of compounds to promote NPC differentiation into oligodendroglia, compounds were supplemented in both proliferation and differentiation media, and clemastine and thyroid hormone were included as positive controls for comparison, given their demonstrated activities on OPC differentiation and myelin formation (Liu *et al*., 2016; Mei *et al*., 2014; Rodriguez-Pena, 1999; Schoonover *et al*., 2004). Remarkably, immunofluorescence labeling of neurons (β-III-tubulin), astrocytes (GFAP), and oligodendroglia (combining PDGFRα and CNP) showed that all 11 selected compounds increased by ∼1.5-fold the number of oligodendroglial cells compared to vehicle (DMSO or PBS, Methods), as illustrated for Sm11 (Fig. 3b, c). In contrast, this increase in oligodendroglia was not found with T3 or clemastine treatment (Fig. 3c), known to only promote OPC differentiation (Barres et al., 1994; Bernal, 2002; Mei *et al*., 2014). Remarkably, the increased number of oligodendroglial cells fostered by our selected compounds was not accompanied by reductions in the number of neurons and astrocytes, nor by changes in overall cell density (Fig. 3d-f). It was rather paralleled by a reduction of undifferentiated NPCs (i.e. unlabeled cells; marker negative cells, Fig. 3g), suggesting that selected compounds foster NPC differentiation towards the oligodendroglial fate, with Sm9 and Sm11 also showing a positive effect on neurogenesis (Fig. 3e). To determine their effect on OL differentiation, we quantified the number of immature OLs (iOLs, So×10^high^ cells being PDGFRα^−^ and MBP^−^), and found a ∼2-fold increase in iOLs (So×10^high^ cells) for T3 and all the selected compounds at 750nM, as illustrated for Sm5 (Fig. 3h, i and Fig. 3 extended data 1). Altogether, these results confirm that our selected compounds have a pro-oligodendrogenic effect in NPC differentiation cultures, without negative impacts on the number of neurons and astrocytes.

**Figure 3.**
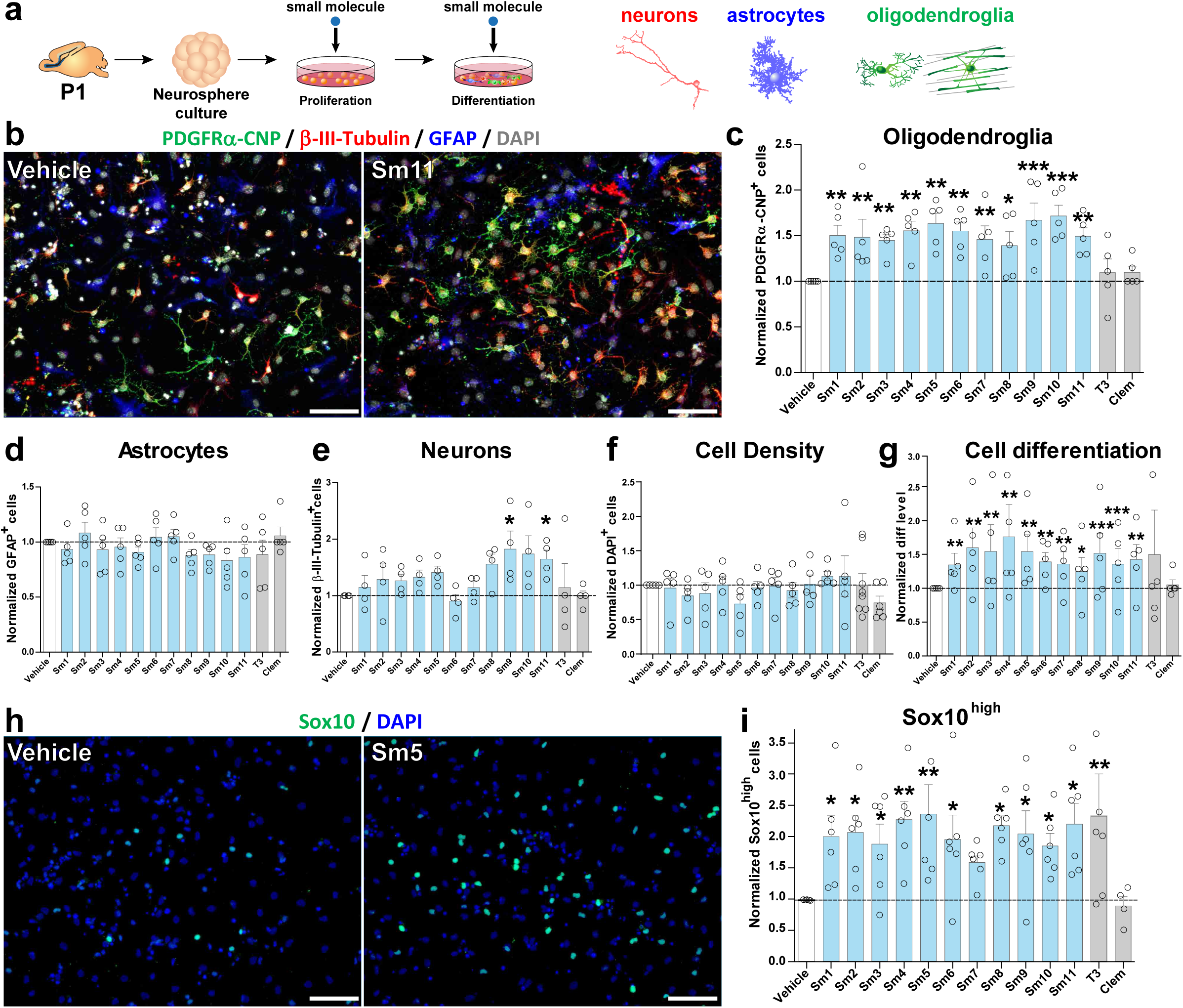
Pro-oligodendrogenic activity of small molecules in neonatal neural progenitor cultures. (**a**) Schematics representing the protocol of neurosphere-derived neural progenitor cell cultures and small molecule administration. (**b**) Representative images illustrating the immunodetection of neurons (ß-III-tubulin^+^ cells, red), astrocytes (GFAP^+^ cells, blue), and oligodendroglia (PDGFRα^+^ OPCs and CNP^+^ OLs, green) in vehicle vs. Sm11 treated cultures after 2 days of differentiation. Note the increase of oligodendroglial cells in the Sm11-treated condition. (**c**-**g**) Quantifications showing the increase of oligodendroglial cells (PDGFRα^+^ and CNP^+^ cells) in cultures treated with each of the selected compounds, but not upon T3 and clemastine treatment (**c**), no changes in the number of astrocytes (GFAP^+^ cells) (**d**), neuronal cells (β-III-tubulin^+^ cells, **e**), or in cell density (**f**), with an increase in cell differentiation (more cells labelled by any of the markers and less DAPI-only cells) for most selected compounds (**g**). (**h**) Representative images of the immunofluorescence for So×10^high^ (iOLs, green) and DAPI (blue) illustrating the increase in treated cultures (Sm5) compared to vehicle cultures after 2 days differentiation. (**i**) Quantification showing the increase of So×10^high^ cells in most compound-treated cultures, including T3. Data are presented as mean ±SEM of fold change normalized to vehicle. N> 5 replicates. *p < 0.05; **p < 0.01; ***p < 0.001. Statistical analysis used linear mixed-effects models followed by Type II Wald chi-square tests (ANOVA). Scale bars: b, 20 μm; h, 50 μm.

### Cell-autonomous effect of selected compounds in OPC differentiation

Given that NPC cultures include several neural cell types, and thus do not allow the evaluation of the direct, cell-autonomous effects of the compounds on OL differentiation/maturation, we turned to primary OPC cultures purified from neonatal (P4) mouse cortices through magnetic cell sorting (MACS). We treated OPCs for three days in the presence of six compounds, for which we did not find any evidence for ongoing research related to oligodendrogenesis (Fig. 4a). Interestingly, automatic quantifications of total cells in each culture (Fig. 4 extended data 1) showed that, while no significant changes were seen in the variable density of PDGFRα^+^ OPCs (Fig. 4b, c), except for Sm7, both T3, clemastine, and all tested compounds (Sm1, Sm2, Sm5, Sm6, and Sm11) increased the number of MBP^+^ OLs compared to vehicle-treated controls (Fig. 4b, d). These results demonstrate the capacity of most selected compounds to cell-autonomously foster OPC differentiation.

**Figure 4.**
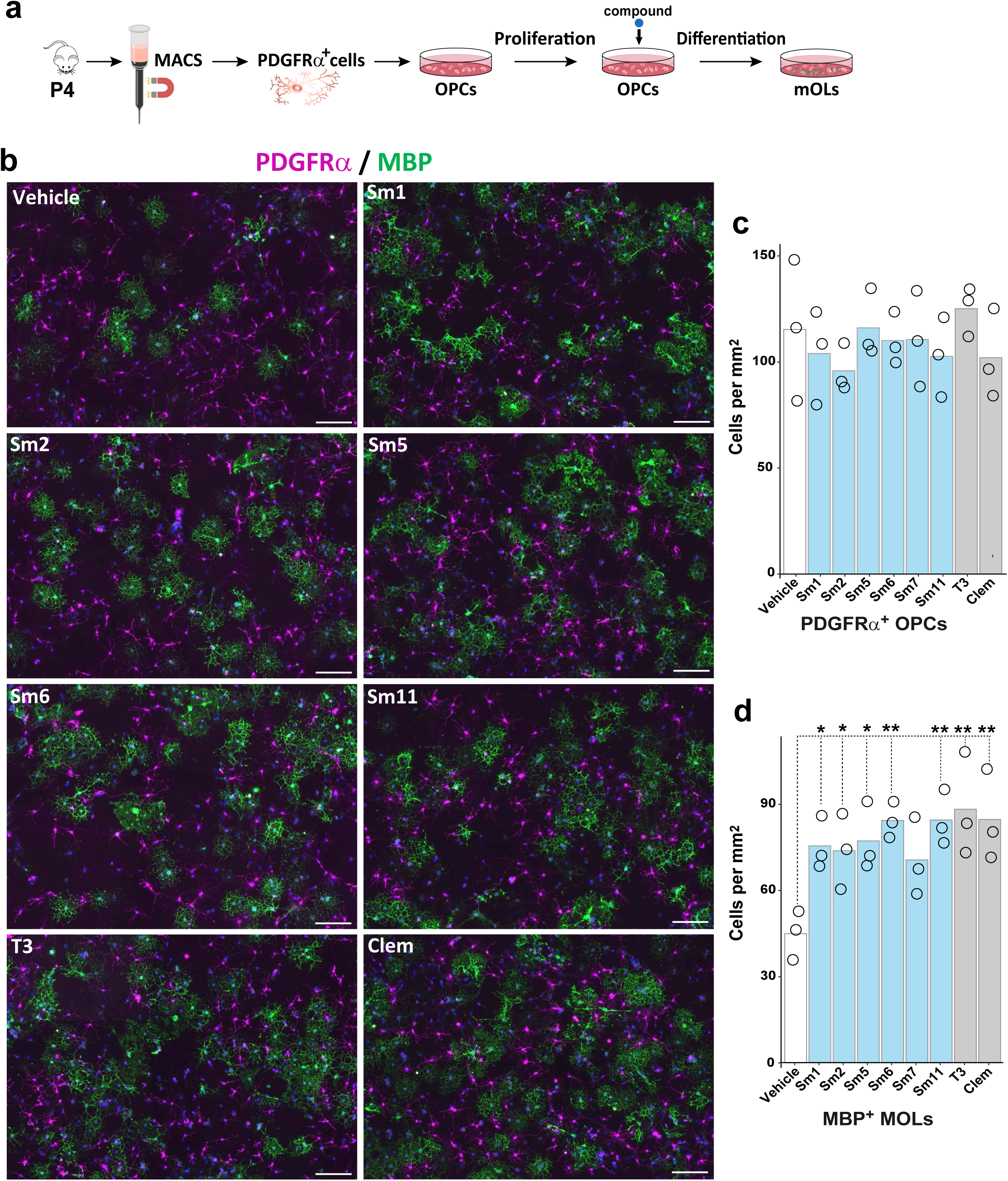
Cell-autonomous effect of selected compounds in OPC differentiation. (**a**) Schematic representing the protocol of OPC purification, culture, and compound treatment. (**b**) Representative images illustrating the immunodetection of OPCs (PDGFRα^+^ cells, magenta) and differentiating OLs (MBP^+^ cells, green). (**c, d**) Quantification of the number of OPCs (PDGFRα^+^ cells, **c**) and number of OLs (MBP^+^ cells, **d**) per mm^2^ in different treated conditions, showing that most compounds present a significant increase in the number of differentiating OLs compared to the vehicle treatment. Data are presented as mean +/− SEM from 3 independent experiments. * p <0.05, ** p <0.01. ANOVA with Dunnett’s post hoc test. Scale bars: 20 μm.

### Selected compounds promote oligodendrocyte differentiation and myelination *ex vivo* in cerebellar explant cultures

Validation of these compounds as a potential treatment for (re)myelination requires further proof of concept before translation into pre-clinical animal models. To this end, we used cerebellar slices (Baudouin et al., 2021) to identify the best compounds having pro-oligodendrogenic and pro-myelinating activities *ex vivo*. This model allowed us to better assess the effect of the selected compounds on OL maturation, myelination, and cytotoxicity in a richer cellular system, also containing immune cells (microglia/macrophages) that are known to influence oligodendrogenesis and (re)myelination (Hagemeyer et al., 2017; Hughes and Appel, 2020; Kirby et al., 2019; Lloyd et al., 2019; Miron et al., 2013; Nemes-Baran et al., 2020; Shen et al., 2021). The onset of myelination in this model takes place after 7 days in culture (Fig. 5a). Therefore, we incubated the cerebellar explants for three days (7-10 days *in vitro*, DIV) in the presence of the six compounds showing the highest pro-oligodendrogenic activity in culture (Sm1, Sm2, Sm5, Sm6, Sm7, Sm11), as well as two positive controls (T3 and clemastine), and their associated negative control (vehicle). We analyzed the effect on OL numbers (CC1^+^/So×10^+^ cells) and myelination (CaBP^+^ Purkinje cell axons co-labeled with MBP), by immunodetection of all four markers in the same sections. A ‘differentiation index’ was calculated as the ratio of So×10^+^ cells being CC1^+^, and the ‘myelination index’ as the ratio of CaBP^+^ axons being MBP^+^ (Baudouin *et al*., 2021). Remarkably, all compounds presented an increased differentiation index compared to their negative controls, but only Sm1, Sm2, Sm5, Sm11, and clemastine reached statistical significance (Fig. 5b, c). Moreover, Sm2, Sm5, and Sm11 also induced a robust increase in the myelination index, comparable to the effect of T3 (Fig. 5d, e). Therefore, these results show the pro-oligodendrogenic and pro-myelinating activities of Sm2, Sm5, and Sm11 in cerebellar explant cultures, and together with previous results, demonstrate the pro-oligodendrogenic capacity of these compounds in different brain regions (cerebral cortex and cerebellum). Altogether, based on the global effects of these molecules combining our *in vitro* and *ex vivo* experiments (summarized in Fig. 5 extended data 1a), and their intended clinical application, we selected Sm5 (dyclonine) and Sm11 (leucovorin) as the best candidates to test their efficiency to promote oligodendrogenesis in pre-clinical models of myelin pathologies.

**Figure 5.**
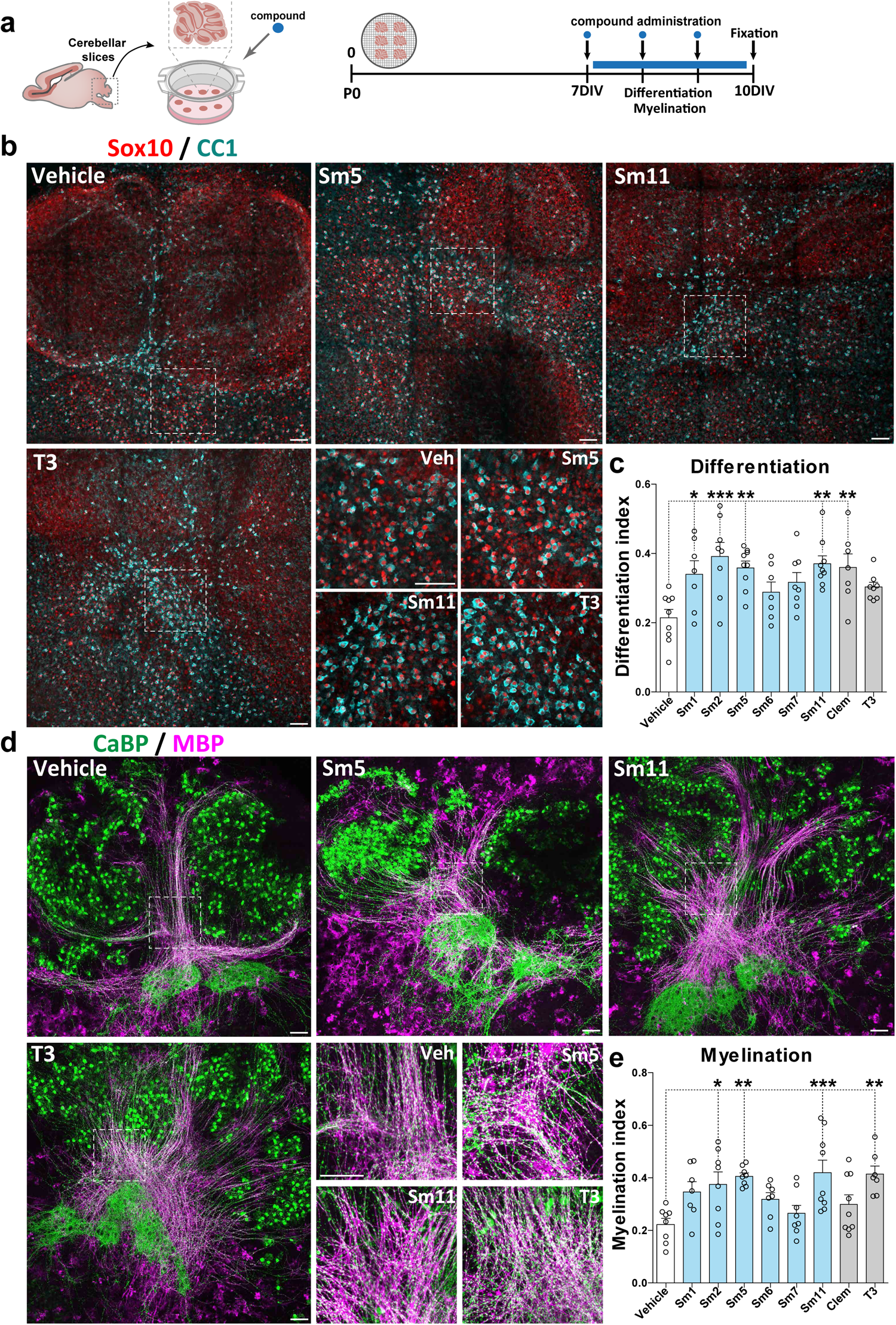
Selected compounds promote oligodendrocyte differentiation and myelination *ex vivo* in cerebellar explant cultures. (**a**) Schematic illustrating the protocol of the cerebellar explant culture model and timing of compound administration. (**b**) Images of explants illustrating effects of compounds (Sm5, Sm11, and T3) on oligodendrocyte differentiation by immunodetection of So×10^+^ oligodendroglia (red) and CC1^+^ OLs (blue). (**c**) Quantification of the differentiation index (SOX10^+^CC1^+^/SOX10^+^ cells) showing an increase following treatment with Sm1, Sm2, Sm5, Sm11, and clemastine compounds. (**d**) Immunofluorescence of explants illustrating compound effects (Sm5, Sm11, and T3) on myelination of Purkinje axons by immunodetection of CaBP^+^ axons/cells (green) and MBP (pink). (**e**) Quantification of the myelination index (surface MBP^+^ CaBP^+^/ surface CaBP^+^) showing an increase following treatment with Sm2, Sm5, Sm11, and T3 compounds. Data are presented as mean ± SEM. N= 5 independent experiments (1-3 confocal acquisitions for each cerebellum). *p < 0.05; **p < 0.01; ***p < 0.001. Statistical unpaired bilateral Wilcoxon Mann Whitney test. Scale bars: 100 μm.

### Dyclonine and leucovorin promote oligodendroglial regeneration in a mouse model of preterm birth brain injury

Chronic hypoxia is a well-established clinically relevant model of very early preterm birth (Jablonska et al., 2012; Salmaso et al., 2014; Scafidi et al., 2013). It is induced by subjecting pups to a low (i.e. 10%) oxygen environment from P3 to P11 (Fig. 6a), which results in diffuse grey and white matter brain injuries, albeit without eliciting a drastic inflammatory response. This period of chronic neonatal hypoxia induces marginal cell death of both neuronal and glial cells along with a delay of OL maturation that persists into adulthood (Scafidi et al., 2009). We assessed the effects of dyclonine (Sm5) and leucovorin (Sm11), as pro-oligodendroglial treatments, by performing intranasal administration of these compounds immediately following the period of hypoxia (i.e. from P11 to P13, see Methods; Fig. 6a and Fig. 6 extended data 1a). This strategy represents a very promising noninvasive technique for drug administration, that was already demonstrated to impact brain cell behavior and eventually their proliferation (Azim *et al*., 2017; Scafidi *et al*., 2013). To examine the impact of dyclonine and leucovorin on proliferation, brains were analyzed at P13, corresponding to the end of the treatment period, after EdU administration one hour before sacrifice to label cells in S-phase. Quantification of EdU^+^ cell density within the dorsal V-SVZ revealed an overall increase in proliferation following hypoxia, with no marked additive effects of the treatment (Fig. 6b, c). Notably, the proportion of Olig2^+^ cells labeled with EdU was significantly increased upon dyclonine and leucovorin treatment (Fig. 6d, e), suggesting an increase in the proliferation of oligodendroglial committed progenitors. Furthermore, the proportion of EdU^+^ cells expressing Olig2 was increased by dyclonine and leucovorin treatment (Fig. 6h), suggesting that these compounds also promote the oligodendroglial fate acquisition from NPCs, in line with their pro-oligodendrogenic activity in neonatal NPC cultures (Fig.3). We then investigated the capacity of dyclonine and leucovorin to rescue the reduction in OL differentiation and maturation induced by hypoxia (Scafidi *et al*., 2013). To this end, OL density and maturation were analyzed within the cortex six days following the end of the treatment, at P19, when the rate of cortical OL maturation and myelination is highest. While the density of oligodendroglial cells (Olig2^+^ cells) was similar between normoxic (Nx; control condition) and hypoxic groups, with no major effects attributable to compounds administration (Fig. 6 extended data 1b, c), their differentiation into CC1^+^/Olig2^+^ OLs was impaired by hypoxia but this difference with normoxic group was rescued by both dyclonine and leucovorin treatments, with leucovorin also reaching statistical difference with the hypoxic group (Fig. 6g, h). We next assessed the effect of hypoxia and treatments on the number of myelinating OLs (mOLs), identified as Olig2^+^ cells expressing GSTπ, a marker restricted to mOLs [(Zhou et al., 2021); Fig. 6i, j, and Fig. 6 extended data 1d, e]. Quantification of Olig2^+^/GSTπ^+^ cell density revealed a marked effect of hypoxia on OL maturation that was fully rescued by leucovorin treatment. In order to substantiate these observations, we extended this analysis by automatically quantifying Olig2^+^/GSTπ^+^ cells to other forebrain regions. At this age, mOLs showed the highest density in ventral brain regions such as the thalamus and hypothalamus when compared to dorsal regions such as the isocortex (Fig. 6k). Further, normalized Olig2^+^/GSTπ^+^ cell densities on Nx mice (Fig. 6l) revealed a stronger effect of hypoxia on OL maturation in ventral brain regions. We confirmed these qualitative observations by performing a manual quantification within the hypothalamus (Fig. 6I). Those confirmed a stronger effect of hypoxia on OL maturation in the hypothalamic lateral zone, compared to cortical region, as well as a full rescue by leucovorin treatment (Fig. 6i-l). In contrast, in this region again, dyclonine treatment did not increase Olig2^+^/GSTπ^+^ cells density when compared to hypoxic mice (Fig. 6k, l). We finally investigated the effect of treatments on MBP immunoreactivity within this ventral region (Fig. 6k). Whereas MBP levels have been shown to be reduced following hypoxia at P11 in several brain regions (Foucault et al., 2024; Chourrout et al., 2022), our results show a return to baseline by P19 in the hypothalamus (Fig. 6 extended data 1f, g). Analyses of MBP optical density and Olig2^+^/GSTπ^+^ cell densities suggest an increase of MBP expression per mOL following dyclonine treatment, while the MBP/mOLs ratio was back to normal in leucovorin-treated animals (Fig. 6m). These results, in line with our previous *in vitro* and *ex vivo* results, confirm the therapeutic capacity of dyclonine and leucovorin to promote oligodendrogenesis in a model of preterm birth brain injury and validate the intranasal approach to administer these compounds and their capacity to cross the blood-brain barrier (BBB). Further, our results show a superior capacity of leucovorin to promote OL maturation restoring normal myelinating OL numbers following injury.

**Figure 6.**
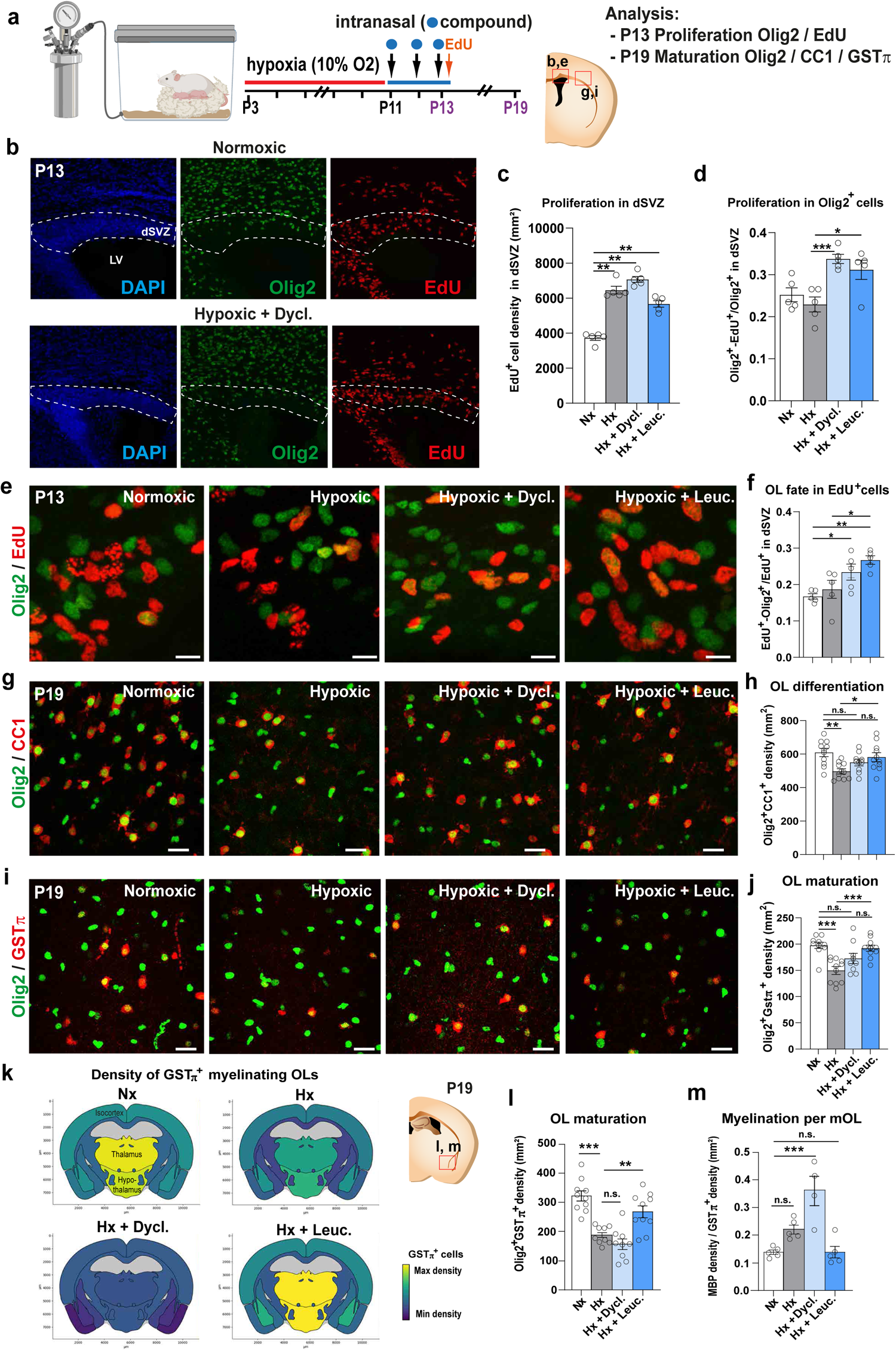
Dyclonine and leucovorin promote oligodendroglial regeneration in a mouse model of preterm birth brain injury. (**a**) Schematic illustrating the workflow used to assess the capacity of dyclonine and leucovorin to promote OPC proliferation and rescue OL maturation following neonatal chronic hypoxia. (**b**) Images of dorsal SVZ at P13, delimited by DAPI counterstaining, showing Olig2 and EdU immunodetection in brain sections of control animals (i.e., normoxic) and following hypoxia and dyclonine treatment. (**c**) Quantification of EdU^+^ cell density (i.e., proliferative cells) in the dorsal SVZ at P13, illustrating the increase in proliferation observed following hypoxia with or without treatment. (**d-f**) Olig2/EdU immunodetection and corresponding quantifications showing that dyclonine and leucovorin increase (**d**) the ratio of proliferative OPCs (Olig2^+^EdU^+^ cells) and (**f**) the OL fate of EdU^+^ cells in the dorsal SVZ at P13. (**g**) Olig2 and CC1 immunodetection and quantification (**h**) showing that dyclonine and leucovorin rescue the reduced density of differentiating OLs (CC1^+^ cells) induced by neonatal chronic hypoxia within the cortex at P19, compared to the density found in the normoxic group, with leucovorin reaching statistical difference with the hypoxic group. (**i**) Olig2 and GST*π* immunodetection and quantification (**j**) showing that only leucovorin rescues the density of myelinating OLs (GST*π*^+^ cells) following hypoxia within the cortex at P19. (**k**) Heatmap representations depicting quantifications of Olig2^+^/GST*π*^+^ cell density in distinct regions of P19 coronal brain sections. Note the reduced density induced by hypoxia (Hx) in most brain regions with a pronounced effect in the thalamus and hypothalamus, compared to normoxic (Nx) control brains, and the rescue of Olig2^+^/GST*π*^+^ cell density following leucovorin (Hx + Leuc.) but not dyclonine (Hx + Dycl.) treatment. (**l**) Quantification of Olig2^+^/GST*π*^+^ cells in the hypothalamic lateral zone of P19 animals confirming that leucovorin, but not dyclonine, treatment rescues the reduced density of hypoxic animals. (**m**) Quantification of the ratio between MBP immunodetection (shown in Fig. 6 extended data **f, g**) and GST*π*^+^ cell density showing an increased myelination per mOL in dyclonine-treated animals. Dyc., dyclonine; Leuc., leucovorin; Data are presented as Mean ± SEM. *p < 0.05; **p < 0.01; ***p < 0.001, n.s., non-significant; scale bars: 100 µm in b, 10 µm in e; 20 µm in g and i.

### Dyclonine and leucovorin promote OPC differentiation while maintaining the OPC pool in a mouse model of adult demyelination

To further assess the pro-oligodendrogenic capacity of dyclonine (Sm5) and leucovorin (Sm11) *in vivo* in the context of myelin pathologies occurring later in life, we used the mouse model of adult focal de/remyelination induced by lysolecithin (LPC) injection into the corpus callosum (Fig. 7a). Compounds were administered via drinking water (see Methods section), in line with oral administration protocols previously used in adult mice and humans (Frye et al., 2018; Okazaki et al., 2018; Rossignol and Frye, 2021; Sahdeo et al., 2014). Notably, no alterations in the drinking behavior of treated mice were observed, ensuring the expected intake of compounds (Fig. 7 extended data 1b). We first analyzed the lesions at 7 days post-lesion (7 dpl) induction, when newly formed OLs have started to remyelinate the lesion (Nait-Oumesmar et al., 1999; Nakatani et al., 2013). The lesion area was identified by the high cellular density (DAPI staining), abundance of microglia/macrophages (Iba1^+^ cells), and reduction of myelin content (using myelin oligodendrocyte glycoprotein, MOG) (Fig. 1 extended data 1c). We determined the effect of the compounds to promote oligodendrogenesis at the lesion site by combinatory immunodetection of Olig2, CC1, and Olig1, allowing to distinguish OPCs (Olig2^high^/CC1^-^/Olig1^nuclear-cyto^ cells) and three stages in OL differentiation (Marie *et al*., 2018; Nakatani *et al*., 2013): immature OL (iOL) 1 (iOL1, Olig2^high^/CC1^high^/Olig1^-^ cells), iOL2 (Olig2^high^/CC1^high^/Olig1^high-cyto^ cells), and mOL (Olig2^low^/CC1^low^/Olig1^low^ cells; Fig. 7b, c). First, we found that dyclonine, and to a greater extent, leucovorin, increased the number of oligodendroglial cells (Olig2^+^ cells) in and around the lesion (Fig. 7c-e_1_, f). Quantification of oligodendroglial stages showed that while the administration of these compounds did not change the density of OPCs (Olig2^high^/CC1^-^/Olig1^nuclear-cyto^ cells) within the lesion area (Fig. 7g), both compounds increased the number of immature OLs. Specifically, leucovorin increased the number of iOL1s (Fig. 7c, h), while both leucovorin and dyclonine increased iOL2s (Fig. 7c, i). Given that both compounds increased the number of differentiating OLs without diminishing the pool of OPCs in the lesion area, we looked at the proliferative status of OPCs, using Mcm2 proliferation marker (Fig. 7i-l), and found that both compounds promote a two-fold increase in the density of proliferative OPCs (Fig. 7m). Finally, calculation of the differentiation ratio (number of iOL2 per OPC) showed that whereas this ratio was 1 OL for 4 OPCs in the vehicle-treated lesions, it increased by 2 folds in dyclonine-treated (2 OLs for 4 OPCs) and by more than 3-fold in leucovorin-treated (3 OLs for 4 OPCs) lesions (Fig. 7n). We then performed an additional experiment to assess for possible improved effects using higher doses of the compounds (10-fold for leucovorin and 2-fold for dyclonine) and to compare their efficacy with that of clemastine, a pro-myelinating compound (Mei *et al*., 2014). At either dose, we observed a similar increase in oligodendroglial (Olig2^+^ cells) and immature OL (iOL1 and iOL2) densities, suggesting that we had reached an optimal dose for eliciting the pro-oligodendrogenic effects of our compounds in this model (Fig. 7 extended data 1d-h). Moreover, while clemastine induced an increase in iOL2 density similar to dyclonine and leucovorin (Fig. 7 extended data 1h), both leucovorin and dyclonine additionally increased the density of proliferating OPCs (Fig. 7m, o), an effect that was absent for clemastine (Fig. 7o).

**Figure 7.**
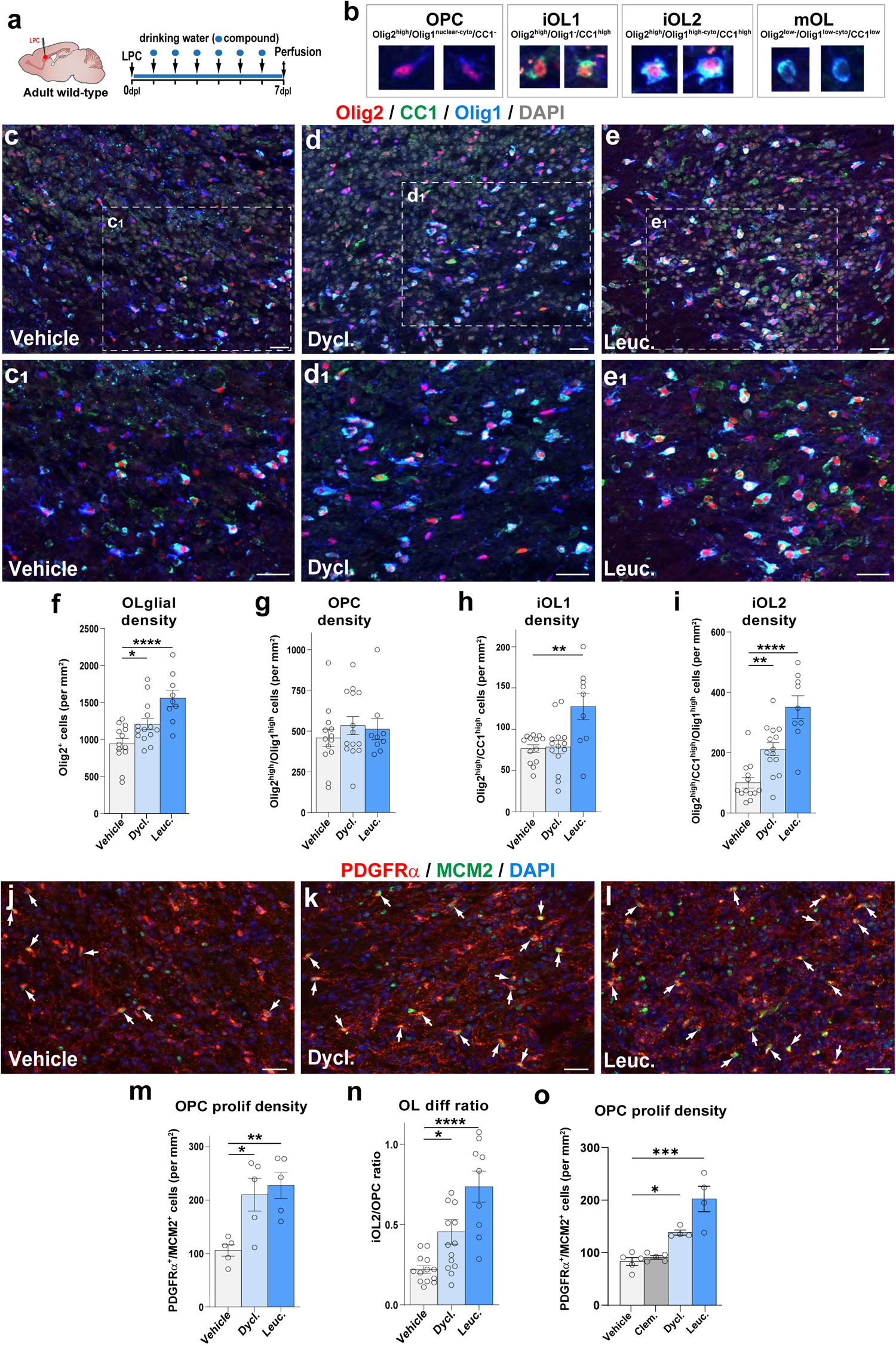
Dyclonine and leucovorin promote both OPC proliferation and differentiation in a mouse model of adult demyelination. (**a**) Schematics illustrating the protocol for LPC demyelination in the corpus callosum, the timing of compound administration in drinking water and analysis. (**b**) Images illustrating the 4 oligodendroglial stages obtained combining Olig2/CC1/Olig1 immunofluorescence: OPCs (Olig2^high^/Olig1^high^), iOL1 (Olig2^high^ /CC1^high^), iOL2 (Olig2^high^/CC1^high^/Olig1^high^) and mOLs (Olig2^low^/CC1^low^/Olig1^low^). (**c**-**e_1_**) Representative images of Olig2/CC1/Olig1 immunofluorescence in the lesion site (depicted by the high cellularity with DAPI) in control mice (c,c_1_, N=13), dyclonine-treated (d,d_1_, N=14), and leucovorin-treated (e,e_1_, N=9) representative of the quantifications shown in f-i. (**f-i**) Quantification showing the increase in the density of Olig2^+^ oligodendroglia in dyclonine- and leucovorin-treated lesions (**f**), no changes in OPC density (**g**), the increase in iOL1 density only in leucovorin treated lesions (**h**), and the increased iOL2 density in dyclonine- and leucovorin-treated lesions (**i**). (**j-l**) Representative images showing the increase in proliferation (Mcm2^+^ cells, green) of OPCs (PDGFRα^+^ cells, red) in the lesion area in dyclonine-treated (**k**) and leucovorin-treated (**l**) compared to vehicle controls (**j**) (N=5 per group). White arrow indicates proliferating OPCs (PDGFRα^+^/Mcm2^+^ cells). (**m**) Quantification of the proliferating OPC density showing a 2-fold increase in the compound treated groups. (**n**) Quantification of the OL differentiation ratio showing 2 to 3 times differentiation increase in dyclonine- and leucovorin-treated lesions respectively. (**o**) Quantification of proliferating OPC density in a replicated experiment comparing with clemastine, showing that contrary to clemastine, both dyclonine and leucovorin increase the OPC proliferation density in the lesion. Dycl., dyclonine; Leuc., leucovorin; Clem., clemastine. Data are presented as Mean ± SEM. *p < 0.05; **p < 0.01; ***p < 0.001. One-way ANOVA statistical test. Scale bars: 20 μm.

### Dyclonine and leucovorin accelerate oligodendrocyte formation and remyelination in a mouse model of adult demyelination

To obtain further proof that the increase in newly formed OLs contributes to accelerated remyelination, we fluorescently labeled adult OPCs and their progeny (OPCs and OLs) by administering tamoxifen to *Pdgfra-CreER^T^; Rosa26^stop-YFP^* mice for five consecutive days before inducing the LPC lesion (Fig 8a). At 10 dpl, we identified by immunofluorescence newly formed OLs (YFP^+^ cells) being either Bcas1, a marker of immature/pre-myelinating OLs (Fard et al., 2017), or GSTπ, a marker restricted to myelinating OLs (Zhou *et al*., 2021). Interestingly we did not find overlap between YFP^+^/Bcas1^+^ cells and YFP^+^/GSTπ^+^ cells, suggesting that indeed Bcas1 and GSTπ identify immature/pre-myelinating and mature/myelinating OLs, respectively. This analysis showed that, similar to clemastine, dyclonine and leucovorin increased more than 2-fold the number of newly formed myelinating OLs (YFP^+^/GSTπ^+^ cells) in the lesion area (Fig. 8b, d) without significant reduction in the lesion volume at this time point likely due to lesion variability (Fig 8c). Performing electron microscopy imaging, we confirmed by ultrastructural analysis the increase in myelinating OLs (identified by their round- or oval-shape nucleus having densely packed chromatin and processes wrapping around myelinated axons; Fig. 8 extended data 1b) in animals treated with leucovorin, dyclonine, and clemastine compared to vehicle-treated animals (Fig. 8e). Moreover, quantification of myelinated axons in the lesion area showed a tendency to decrease the g-ratio of axons in leucovorin-, dyclonine-, and clemastine-treated animals compared to vehicle-treated controls, suggesting an increased remyelination (more wrapping) induced by the compound treatment, that reaches significance in the case of leucovorin treatment for axons larger than 1 μm (Fig. 8f, g). Altogether, these results show that, similar to clemastine, leucovorin and dyclonine promote OL differentiation, thus remyelination, *in vivo* in the context of adult brain demyelination, with leucovorin showing the strongest effect. The possibility of leucovorin and dyclonine directly inducing myelin formation would require further investigation. Moreover, the increased number of oligodendroglial cells in the lesion area together with the increase in OPC proliferation indicate that in the context of adult demyelinating lesions, these compounds are capable of promoting OL differentiation while maintaining the pool of OPCs by fostering their proliferation, an effect not found with clemastine (Fig. 7o).

**Figure 8.**
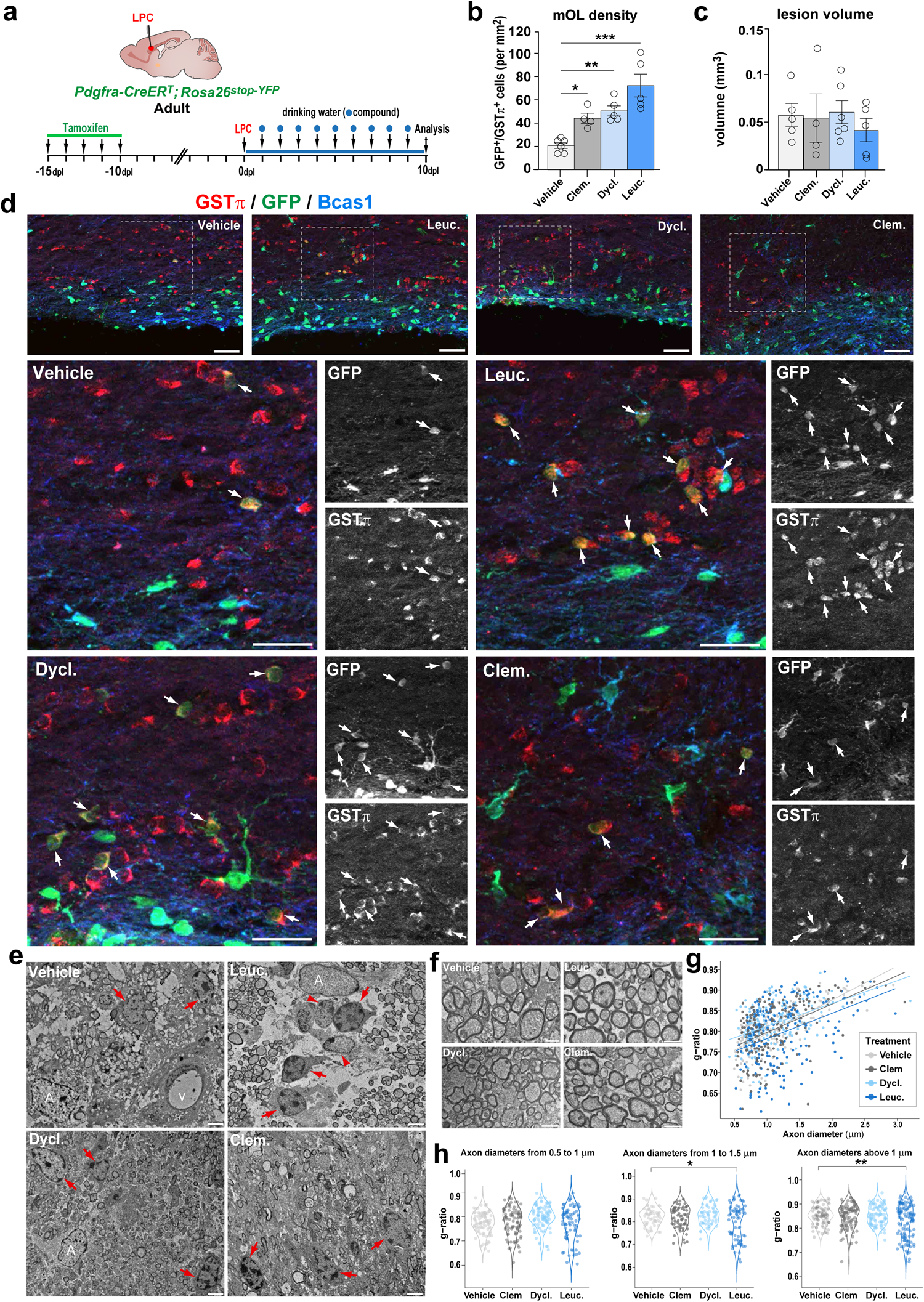
Leucovorin and dyclonine accelerate oligodendrocyte formation and remyelination in a mouse model of adult demyelination. **(a)** Schematics illustrating the protocol used for tracing OPCs and newly formed OLs by YFP-reporter induction prior to LPC demyelination by tamoxifen mediated Cre-recombination of a stop cassette in *Pdgfra-CreER^T^; Rosa26^stop-YFP^* mice, followed by compounds’ administration in the drinking water and analysis at 10 days post-lesion (dpl). **(b)** Quantification of the density of newly formed (GFP^+^) myelinating OLs (GSTπ^+^), showing that dyclonine and leucovorin, alike clemastine, increase the generation of remyelinating OLs compared to vehicle. **(c)** Quantification of the lesion volume indicating no major reduction at 10 dpl in treated animals. (**d**) Representative images of adult generated YFP^+^ immature OLs (Bcas1^+^ cells, blue) or myelinating OLs (GSTπ^+^ cells, red). (**e**) Electron microscopy images at 10 dpl illustrating the increase in mOLs (red arrows) identified by their typical ultrastructural traits (round- or oval-shape nucleus having densely packed chromatin and processes wrapping around) in compounds’ treated lesions compared to controls (Vehicle). Note that leucovorin-treated lesions have more frequent iOLs (arrowheads) in the lesion area, characterized by showing less densely packed chromatin. (**f**) Representative micrographs illustrating remyelinated axons in the lesion area in different treatments. (**g**) Scatter plot representing the quantification of myelin sheath thickness (g-ratio) per axon diameter in the lesion area of vehicle-(light grey), clemastine-(grey), dyclonine-(light blue), and leucovorin-treated (blue) animals indicating a tendency to increase myelin thickness (lower g-ratios) in compound-treated animals compared to vehicle, with leucovorin-treated animals showing the strongest effect. **(h)** Violin plots quantifications of g-ratios in axons with different thicknesses indicating a significant decrease in g-ratio (thicker myelin) of axons above one micrometer in the lesion area of leucovorin-treated animals. A, astrocytes. V, vessel. Clem., clemastine; Dycl., dyclonine; Leuc., leucovorin. Data are presented as Mean ± SEM. *p <0.05; **p <0.01; ***p < 0.001. One-way ANOVA statistical test. Scale bars: 20 μm.

### Dyclonine and leucovorin accelerate myelin clearance and microglial transition from pro-inflammatory to pro-regenerative profiles in a mouse model of adult demyelination

Finally, given the recognized pro-remyelinating properties of microglia (Lloyd and Miron, 2019) and the capacity of our compounds to repress inflammatory genes such as Interferon gamma (Fig. 2d), we investigated their potential anti-inflammatory and pro-regenerative activity mediated by microglia/macrophages (Iba1^+^ cells). We assessed various microglial profiles, including phagocytic (CD68^+^/Iba1^+^ cells), pro-inflammatory (Cox2^+^ and iNOS^+^ cells), and pro-regenerative (Arg1^+^/Iba1^+^ cells) microglia, known to follow the dynamics of lesion repair in this model (Lloyd *et al*., 2019; Miron *et al*., 2013). Interestingly, within the lesion area at 7 dpl, there was a reduction in the fraction of phagocytic microglia/macrophages (CD68^+^ cells; Fig. 9a, c) and a reduction in the area of myelin debris labeled by dMBP (Shen *et al*., 2021) accompanied by an increased proportion of the dMBP phagocyted by CD68^+^ cells (Fig. 9a, d, e), suggesting faster myelin clearance. Furthermore, we found a decrease in Cox2^+^ and iNOS^+^ pro-inflammatory microglial profiles (Fig. 9f-h), paralleled with a strong increase in pro-regenerative microglial phenotype (Arg1^+^ cells; Fig. 9i, j), suggesting that both dyclonine and leucovorin promote an earlier transition between these microglia/macrophage profiles.

**Figure 9.**
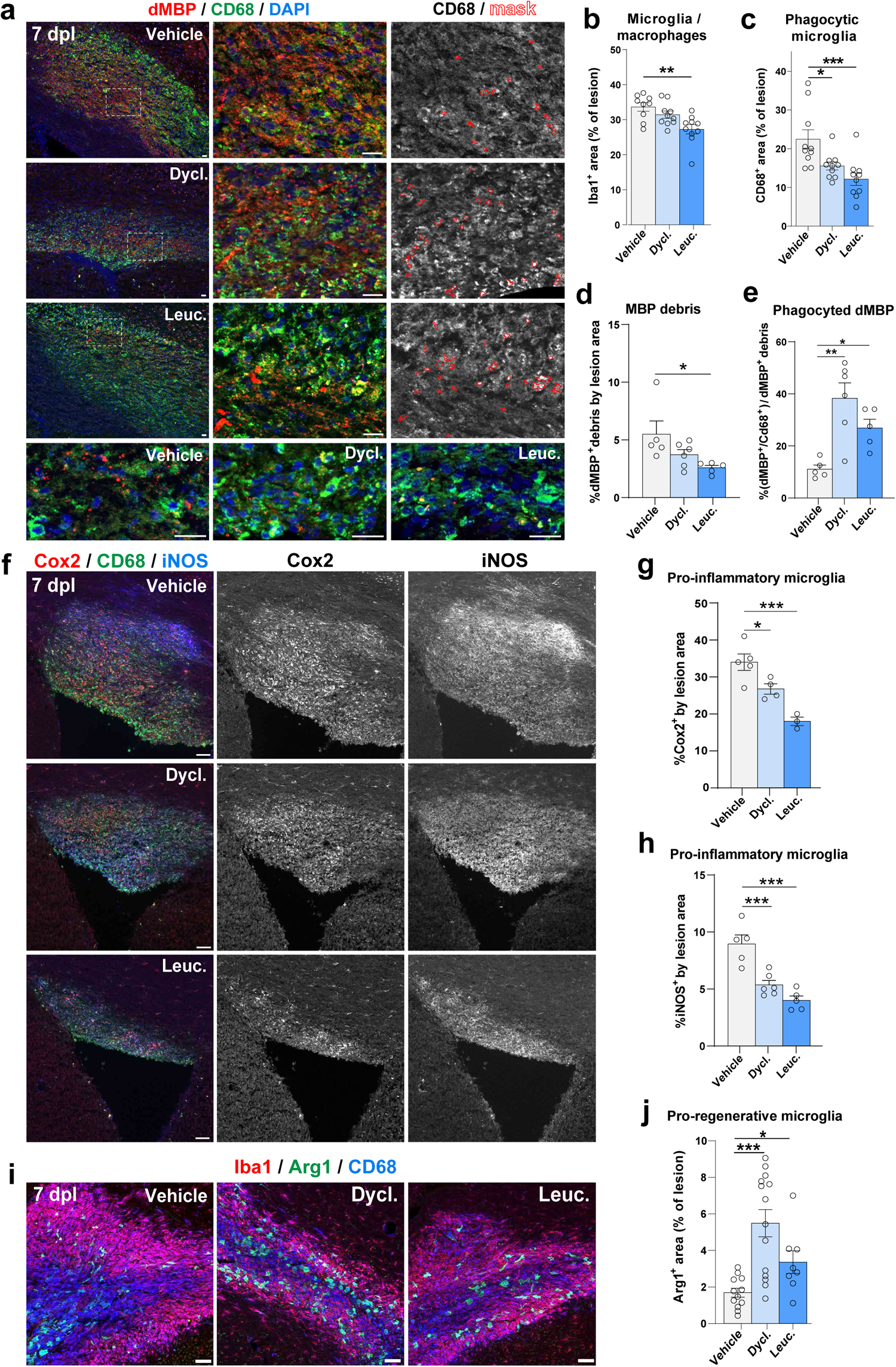
Leucovorin and dyclonine accelerate myelin clearance and microglial transition from pro-inflammatory to a pro-regenerative profiles in a mouse model of adult demyelination. **(a)** Representative images of the lesion territory immunodetecting myelin debris with MBP (dMBP, red) and phagocytic microglia with CD68 (green), showing increased dMBP signal inside CD68^+^ cells in dyclonine- and leucovorin-treated lesions (yellow dots) and *vs.* more dMBP outside CD68^+^ cells (red dots) in vehicle-treated lesions at 7 days post-lesion (dpl). Right panels illustrate the mask of automatic quantification for CD68 and dMBP colocalization (red labels). Bottom panels are higher magnification images for dMBP dot visualization in different treated lesions. (**b-e**) Barplots and dotplots representing the quantification in the lesion of Iba1^+^ microglial area (**b**), CD68^+^ phagocytic area (**c**), myelin debris as percentage of dMBP area (**d**), and phagocyted myelin debris as the percentage of dMBP area inside CD68^+^ cells from the total dMBP area (**e**). Note the strong increase of phagocyted myelin debris within dyclonine- and leucovorin-treated lesions. **(f)** Representative pictures showing immunodetection of microglia/macrophages (CD68^+^ cells, green) in the lesion area expressing inflammatory markers (Cox2 in red, and iNOS in blue). (**g**-**h**) Quantification of the percentage of lesion area labeled by Cox2 (**g**) and iNOS (**h**) immunofluorescence, in dyclonine-treated (N=5) and leucovorin-treated (N=4) mice compared to vehicle (N=5). Note the decrease in the pro-inflammatory profiles of microglia in leucovorin and dyclonine-treated conditions. (**i**) Representative pictures showing immunodetection of microglia/macrophages (Iba1^+^ cells, red) in the lesion area presenting phagocytic (CD68^+^ cells, blue) and pro-regenerative (Arg1^+^ cells, green) profiles. Note the increase in the pro-regenerative profiles of microglia in leucovorin and dyclonine-treated conditions. (**j**) Histograms representing the density of pro-regenerative microglia, in dyclonine-treated and leucovorin-treated mice compared to vehicle. Data are presented as Mean ± SEM. *p < 0.05; **p < 0.01; ***p < 0.001. One-way ANOVA statistical test. Scale bars: 20 μm.

In summary, these results demonstrate that in the context of adult brain de/remyelination, oral administration of dyclonine and leucovorin promotes the generation of newly formed OLs while preserving the OPC pool size. Furthermore, in addition to their direct effects on OLs, these compounds also appear to improve lesion repair by accelerating myelin clearance and promoting the transition from pro-inflammatory to pro-regenerative microglial profiles.

## Discussion

To date, no medication demonstrating convincing remyelination efficacy in humans has been approved for treating myelin pathologies, including preterm-birth brain injuries (PBI) and multiple sclerosis (MS). While most previous studies have followed a gene/pathway candidate approach, here we used a more comprehensive strategy to fill this gap. Leveraging transcriptomic datasets through a pharmacogenomics analysis and developing an expert curation of genes previously involved in oligodendroglial biology (provided as a resource for the scientific community: [OligoScore](https://oligoscore-staging.icm-institute.org/), https://oligoscore.icm-institute.org), we identified and ranked novel small bioactive molecules (compounds) fostering transcriptional programs associated with various aspects of oligodendrogenesis, including OPC proliferation, differentiation, and (re)myelination. We then demonstrated the pro-oligodendrogenic activity of some of these compounds in mice using several approaches: i.e. (1) *in vitro,* in cultures of neural progenitor cells and primary oligodendroglial progenitors, (2) *ex vivo,* in cerebellar explant cultures, and (3) *in vivo,* using both a model of perinatal chronic hypoxia and a model of adult focal demyelination with spontaneous remyelination. Compounds repurposed to promote remyelination in MS currently in phase III clinical trials, such as clemastine and thyroid hormone analogs (Gingele and Stangel, 2020; Green *et al*., 2017; Mei *et al*., 2014), generate OLs at the expense of OPCs, thus potentially depleting OPCs in the long term. Remarkably, the compounds we identified here, leucovorin and dyclonine, promote the generation of new OLs while maintaining the pool of OPCs stable by inducing their concomitant proliferation and differentiation, an effect not found using clemastine (Fig. 7). The dual capacity of leucovorin and dyclonine to promote at the same time OPC proliferation and differentiation in the context of adult remyelination, is supported by our pharmacogenomics analysis predicting that both leucovorin and dyclonine induce the expression of *Olig2, Ascl1, Sox2, and Myt1* (Supplemental Tables S7, S11, S12), key transcription factors promoting both proliferation and differentiation programs (Nielsen et al., 2004; Ligon et al., 2007; Nakatani et al., 2013; Zhang et al., 2018; Vue et al., 2020). Finally, in the context of an adult demyelination model, leucovorin and dyclonine also improve lesion repair by promoting in parallel a microglia-mediated myelin debris clearance, and a faster transition from pro-inflammatory to pro-regenerative microglial profiles. Therefore, these compounds represent promising candidates for a more comprehensive and sustained regenerative response in various oligodendroglial pathologies, with leucovorin showing the largest beneficial effects in all conditions tested here.

Selected for their broad transcriptional effects, leucovorin and dyclonine may increase oligodendrogenesis and (re)myelination by targeting multiple cell types and mechanisms. Concerning targeted cell types, our results obtained in neonatal neural progenitor cultures and in the neonatal hypoxia model *in vivo* indicate that leucovorin and dyclonine act onto neural progenitor cells to promote their differentiation into oligodendroglial cells. Second, using purified OPC primary cultures, we demonstrate that they can act cell-autonomously in OPCs to promote their differentiation/maturation, in agreement with a recent study showing that folic acid, also involved in folate metabolism, can enhance oligodendrocyte differentiation (Weng et al., 2017). Third, in the *in vivo* context of a neonatal hypoxia model of PBI, both compounds can promote oligodendroglial proliferation, with only leucovorin efficiently restoring the number of myelinating OLs (Fig. 6). It is interesting to note that whereas dyclonine did not recover normal myelinating OL density, myelination however returned to baseline as revealed by optical density measurement of MBP expression. This suggests that dyclonine did not impede compensatory myelin production by the remaining OLs. The long-term consequences of this incomplete recovery and the possible increase in myelination capacity of individual OLs upon dyclonine treatment would need further investigation. Fourth, in the *in vivo* model of adult focal demyelination with spontaneous remyelination, both leucovorin and dyclonine accelerate the generation of newly formed and remyelinating OLs, while maintaining the pool of OPCs by increasing their proliferation (Figs. 7 and 8). This capacity to maintain the pool of adult OPCs, not found using clemastine, has important implications for the long-term treatment of MS patients, whose remyelination capacity decreases with age and disease progression (Neumann et al., 2019a; Neumann et al., 2019b). Finally, leucovorin and dyclonine also target microglia/macrophages, accelerating their phagocytosis of myelin debris and their transition from pro-inflammatory (Cox2^+^ and iNOS^+^ cells) to pro-regenerative (Arg1^+^/Iba1^+^ cells) profiles. These effects, which have also been described in clemastine treatments (Apolloni et al., 2016; Su et al., 2018), are known to favor OPC differentiation and remyelination in this model (Hagemeyer *et al*., 2017; Hughes and Appel, 2020; Kirby *et al*., 2019; Lloyd *et al*., 2019; Miron *et al*., 2013; Nemes-Baran *et al*., 2020; Shen *et al*., 2021; Van Steenwinckel et al., 2019).

Dyclonine and leucovorin have been approved by the FDA in 1955 and 1952, respectively. Dyclonine has been used as an oral anesthetic administered in throat lozenges and is known to inhibit Nav1.8, a voltage-dependent Na channel primarily expressed on small, unmyelinated peripheral sensory neurons, most of which are nociceptors (Akopian et al., 1996). Recent findings identified dyclonine’s ability to enhance synaptic activity in cultured hippocampal mouse neurons, with chronic treatment of mice over several months in their drinking water leading to increased respiration and ATP production in brain mitochondria (Varkuti et al., 2020). In a study of drug repositioning in the context of Friedreich Ataxia (Sahdeo *et al*., 2014), dyclonine was found to confer protection against diamide-induced oxidative stress through binding to the transcription factor NRF2 (nuclear respiratory factor 2) and activation of the NRF2 pathway (Rufini et al., 2022). Finally, dyclonine may also mediate pro-oligodendrogenic effects through its suggested regulation of different hormones, including estrogen receptors, thyroid hormone receptor β, and the androgen receptor (Varkuti *et al*., 2020) which represent possible therapeutic targets in MS (Zahaf et al., 2023).

Leucovorin (aka folinic acid or folinate), is a naturally occurring folate (from Latin *folia,* foliage), a general term designing molecularly-related metabolites participating in the folate cycle, a metabolic pathway that, depending on cell requirements, can lead to nucleotide synthesis, mitochondrial tRNA modification, or methylation, respectively impacting proliferation, mitochondrial respiration, and epigenetic regulation (Zheng and Cantley, 2018). Abnormal folate metabolism has been causally linked to a myriad of diseases, but it is often unclear which biochemical processes and cellular functions are affected in each disease (Zheng and Cantley, 2018). Molecularly different from folic acid, a synthetic folate, leucovorin has a different entrance and metabolism in the folate cycle. Both leucovorin and folic acid can be transformed by different enzymes into 5-methyl-tetrahydro-folate, which efficiently crosses the BBB using the folate receptor alpha (FRα) transporter, and is thought to be the main active folate in the CNS (Menezo et al., 2022; Scaglione and Panzavolta, 2014). Leucovorin has the advantage over folic acid of using other transporters present in the choroid plexus to get into the CNS, i.e., the proton-coupled folate transporter (PCFT) and the reduced folate carrier (RFC) (Mafi et al., 2020). Moreover, folic acid must be metabolized by dihydrofolate reductase (DHFR) to enter the folate cycle, with this reaction being slow and easily reaching saturation, thus limiting the therapeutic use of high doses of folic acid (Menezo *et al*., 2022; Scaglione and Panzavolta, 2014). For these reasons, leucovorin has been used to treat epileptic patients having mutations in the folate receptor alpha gene, FOLR1 (Mafi *et al*., 2020), as well as in the context of cancer chemotherapy to decrease the toxic effects of methotrexate, an inhibitor of the DHFR. Our results showing that leucovorin increases oligodendrogenesis and (re)myelination both *in vitro* and *in vivo* suggest that it promotes these processes at least in part by fostering high levels of folate metabolism. Moreover, the current clinical administration of leucovorin to adults for large periods guarantees its long-term administration to MS patients without major side effects. Finally, some studies point out that molecules involved in the folate cycle can reduce the inflammatory response by either inhibiting TNFα, IL-1β, or iNOS-dependent NO production (Cianciulli et al., 2016; Tommy et al., 2021). These anti-inflammatory effects parallel our pharmacogenomics strategy indicating that leucovorin downregulates the gene coding for Interferon gamma (*IFNG*), a key inflammatory cytokine, and our results in adult remyelinating lesions indicating that leucovorin acts in microglia/macrophages accelerating the transition from pro-inflammatory to pro-regenerative profiles. Altogether, our current understanding of dyclonine and leucovorin activities warrants the broad pro-regenerative activities found in our mouse models of myelin pathologies, with superior effects of leucovorin. Nevertheless, future studies will be required to elucidate the specific mechanisms of action of these compounds in oligodendroglial development, remyelination, and lesion repair.

In conclusion, our study underscores pharmacogenomic analysis as an efficient and low-cost approach to identify pro-oligodendrogenic compounds. By developing a scoring system for genes implicated in oligodendrogenesis ([OligoScore](https://oligoscore-staging.icm-institute.org/)), we could rank the most promising compounds to demonstrate their pro-oligodendrogenic and pro-myelinating activities using both *in vitro* and *ex vivo* murine culture systems. This resulted in the selection of leucovorin and dyclonine, two FDA-approved compounds, which we demonstrated to have brain repair and pro-oligodendrogenic activities in both preterm birth brain injury and adult demyelination mouse models, with leucovorin demonstrating broader beneficial effects. This work paves the way to clinical trials repurposing these compounds to promote brain repair in the context of both early- and late-life-onset myelin pathologies such as PBI and MS.

### Limitations of the study

Despite the cellular (neural precursor cells, oligodendroglia, and microglia) and molecular mechanisms (folate cycle and anti-oxidative stress metabolism) underlying the beneficial effects of leucovorin and dyclonine in oligodendrogenesis and myelin repair shown here, follow up studies are crucial to unravel the particular mechanisms underlying the effects of each compound in each pathological context, and in cell types, including neurons and astrocytes. For clinical translation in treating preterm birth brain injuries, the capacities of leucovorin and dyclonine to rescue other aspects of preterm birth brain injury, such as inflammation and malnutrition, will need further investigation. Also, the increase in OPC proliferation and numbers mediated by dyclonine in the hypoxia model do not rescue the density of myelinating OLs, contrasting with its effects in adult brain de/remyelination, calling for a better understanding of the environmental differences of each pathological model, such as microglial involvement. The proposed increased myelination capacity per OL (e.g. increased number of internodes) mediated by dyclonine treatment in the hypoxia model will require direct experimental evidence. Finally, further studies are needed to explore the capacity of leucovorin and dyclonine in promoting myelin repair in the context of the aging brain, which is only obtained with clemastine or T3 treatment when combined with metformin (Neumann et al., 2019a). Thus, considering the possible synergy between leucovorin and dyclonine with metformin in the context of the aging brain constitutes an essential follow-up pre-clinical research before starting clinical trials in the context of MS lesion repair.

## Methods

### Data Processing and Oligodendroglial Gene Enrichment

Data Selection: transcriptomic datasets were selected from single-cell and bulk-transcriptomic studies generated in neural cell types during mouse development at the embryonic, postnatal, and adult stages (Falcão *et al*., 2018; Jäkel *et al*., 2019; Marques *et al*., 2018; Marques *et al*., 2016; Saunders *et al*., 2018; Zhang *et al*., 2014). Methods **Table 2** summarizes the datasets used, criteria for selecting DEGs, p-values, fold-changes, GSE numbers, and data details. For most of the datasets, enriched genes were identified using supplementary tables provided in the publications, selecting DEGs (≥ 1.5-fold change) or genes with enriched expression (p < 0.05). Detailed methods for specific dataset analyses are provided below.

#### Postnatal Dorsal NSCs and TAPs

we used data from Raineteau’s lab (GSE60905) focusing on neural stem cells (NSCs) and transit amplifying precursors (TAPs) isolated at P4 from the dorsal and lateral SVZ regions, using transgenic mice and specific markers for each cell type (i.e., *Hes5::GFP* and *prominin-1* mice for NSCs and *Ascl1::GFP* mice for TAPs). The microarray data were processed in two phases (see ‘*dNSCs/TAPs*’ script): Normalization: Using R and the ‘affy’ package, data were normalized with the Robust Multiarray Averaging (RMA) method (Irizarry et al., 2003). Differential Analysis: Limma analysis (Ritchie et al., 2015) identified DEGs with a filter of FC > 1.2 and p < 0.05.

#### Oligodendrocyte Lineage in Postnatal Brain

we used data from Ben Barres’ lab (GSE52564), and using RNA-Seq from purified cells, we identified oligodendrocyte-specific genes. Cells were isolated at P7 (astrocytes and neurons) and P17 (oligodendrocyte lineage cells). Filtering criteria included FPKM > 0.5 to remove low-expression genes and a final filter for overexpressed/repressed genes in the oligodendrocyte lineage (FC > 1.2 to 9.2). The ‘*dNSCs/TAPs*’ transcriptional signature was combined with the ‘*OL lineage*’ transcriptional signatures to obtain lists of intersection genes (‘*OL specification*’ signature) and exclusion genes (‘*OL differentiation*’ signature) (see results section).

#### Single-cell RNA-seq Oligodendroglial Clusters

using Marques and colleagues’ (2016) supplemental table ‘aaf6463 Table S1’, sheet ‘specific genes’ we pooled all genes except those of VLMC column obtaining 532 unique genes. To extract more information, raw data from 5072 cells were processed with the Seurat package v2 (Stuart et al., 2019) using the following criteria: min.cells = 10; min.genes = 500; mitochondrial percentage <0.10, 20 dimensions to find neighbors, a resolution of 0.9 to find clusters, selecting the top 150 genes from each cluster from FindAllMarkers function, identifying 1598 DEGs (Methods **Table 2**).

### Pharmacogenomics Using the SPIED Platform

The transcriptional signatures obtained were used to query the CMAP 2.0 database via the wSPIED platform. The CMAP 2.0 (CMAP2.0) database consists of transcriptional profiles corresponding to the effects of small molecules at various concentrations and treatment times on panels of human cell lines (∼100). This enables the identification of therapeutic molecules capable of inducing transcriptional changes similar to those observed during oligodendrogenesis. Query files were simple text files with columns containing gene IDs and their enrichment/repression (+1 or −1, respectively). The output table contained identifiers of small molecules (correlative and anti-correlative), ranked by Z-score (Pearson-type regression analysis), and p-value.

### Gene Ontology and Cell-specific Enrichment Analysis

GO analysis of selected genes was performed using the Cluster Profiler R-package (2012), [Metascape website](https://metascape.org/gp/index.html#/main/step1) (Zhou et al., 2019), [EnrichR website](https://maayanlab.cloud/Enrichr/) (Chen et al., 2013), [Panther website](https://pantherdb.org/) (Mi et al., 2019).

### Expert Curation and Scoring of Genes Regulating Oligodendrogenesis

We implemented a knowledge-driven scoring procedure by curating over 1000 publications involving functional studies demonstrating the role of specific genes in oligodendrogenesis (OPC specification, proliferation, migration, survival, differentiation, myelination, and remyelination) under physiological or pathological conditions. This curation implicated 430 genes, scored for each process from 1 to 3 (low, medium, strong), either positively or negatively, based on their function in the process. To share our expert curation scoring strategy, we established a user-friendly platform OligoScore (OligoScore platform) evaluating the regulatory impact on transcriptional programs involved in oligodendrogenesis and (re)myelination.

### Validation of the OligoScore Strategy

We validated the OligoScore strategy using genes enriched in specific oligodendroglial stages and genes deregulated in OPCs upon perturbations. First, we queried OligoScore with the top 2000 genes enriched in OPCs and myelinating oligodendrocytes (mOLs), obtained by comparing their transcriptomes from purified brain cells (RNAseq (Zhang *et al*., 2014)) and sorting them by their expression ratio between OPCs and mOLs, respectively. Using oligodendroglial single-cell transcriptomes (Marques *et al*., 2018; Marques *et al*., 2016), we performed a similar analysis using genes enriched in OPCs (723 genes, logFC > 0.5 in OPC and cycling OPC clusters versus mOL1/2 clusters) and in mOLs (1428 genes, logFC > 0.5 in mOL1/2 clusters versus OPC and cycling OPC clusters). Results confirm expectations, showing that genes enriched in OPCs are involved in several processes of oligodendrogenesis (specification, proliferation, differentiation, and (re)myelination; **Fig. 2 Extended Data 2a, c**), while genes enriched in mOLs are mainly involved in differentiation and myelination (**Fig. 2 Extended Data 2b, d**). Second, we queried OligoScore with genes differentially expressed in OPCs upon genetic (P7 Chd7-deleted OPCs, p-value < 0.01; Marie et al., 2018), or environmental perturbation (P10 OPCs in the systemic IL1β-mediated neonatal neuroinflammatory model, FDR < 0.05; Schang et al., 2022). This identified the affected processes in agreement with these studies: reduced survival and differentiation of Chd7-iKO OPCs (**Fig. 2 Extended Data 2e**), and increased proliferation and reduced differentiation of P10 OPCs in the IL1β-model (**Fig. 2 Extended Data 2f**). Additionally, this analysis provided mechanistic insights, identifying genes responsible for the deregulation of each oligodendroglial process. Gene sets used to query OligoScore and the full results are provided in the Excel file named ‘OligoScore validation tables’.

### Hub Genes and Small Molecules List Generation

To obtain either hub genes or small molecules, we interrogated the SPIED3 version (a searchable platform-independent expression database III; SPIED3; GeneHubs and CmapG function). To generate a list of hub genes, we uploaded gene symbols corresponding to the large oligodendroglial gene subset (3372 genes), whereas for small molecules, we used gene symbols corresponding to either the large oligodendroglial gene sets (1898, 2099 and 3372 genes) and to oligodendroglial curated gene sets for each process regulating oligodendrogenesis (with their positive or negative score attributed for their respective influence in each process). Output for a hub gene or a compound contains a similarity score (correlation score with the query) and a statistical significance of the similarity score, associated with a probability of finding the same genetic association.

### Pharmacogenomic Analysis

The 156 compounds obtained in common between both approaches were scored using <u>OligoScore</u> and ranked by their scores in the curated target genes for each process of oligodendrogenesis. We also performed a systematic analysis based on gene ontology databases and cell-specific enrichment (see methods for GO analysis) to exclude compounds showing potential side effects (e.g., effects onto other lineages and/or cell death). Finally, we also selected compounds based on their functional properties and signaling pathways modulated using publicly available drug repository websites ([DrugBank](https://go.drugbank.com/); [KEGG Drug](https://www.genome.jp/kegg/drug/); [GeneCards](https://www.genecards.org/)).

### Pharmacological Analysis

The ADMET parameters were calculated by the ACD/Percepta software (v.2019.1.2 built 3200) ([ACD/Labs Percepta](https://www.acdlabs.com/products/percepta-platform/)). Predictive models for BBB permeability are based on the rate and extent of brain penetration (respectively log PS and log BB) constants, built using non-linear least squares regression validated by an internal validation set, and two other experimental external validation sets (Kalvass et al., 2007; Summerfield et al., 2008). Physicochemical properties such as lipophilicity (Log P), number of hydrogen bond acceptors and donors, ion form fraction at pH 6.5, and McGowan volume were calculated with the ACD/Labs Algorithm Builder 1.8 development platform and used for the modeling.

The combination of brain/plasma equilibration rate (Log (PS * fraction unbound, brain)) with the partitioning of compounds at equilibrium (Log BB) classifies compounds as either active or inactive on the CNS. The model is validated using experimental data from more than 1500 compounds with CNS activity (Adenot and Lahana, 2004). This software uses a human expert rules system (Benigni and Bossa, 2006; Kazius et al., 2005; Kazius et al., 2006) to predict mutagenicity, clastogenicity, carcinogenicity, and reproductive toxicity. The model predicting “health effects on particular organs or organ systems” is based on RTECS and ESIS databases for more than 100.000 compounds. The experimental data have been collected from RTECS and ESIS databases, and diverse publications.

Health effects predictions are known to be very sensitive, so we reduced the weight of these predictions in our selection strategy for the 11 molecules. These results are completed by a reliability index (RI) ranging from 0 to 1, where 0 is unreliable for the prediction and 1 is a fully reliable prediction. This RI value tends to zero in two cases: first when the overall similarity between the tested compound and the most similar compounds used for the correction of the global model is weak, and second when an inconsistent variability between the predicted and experimental values was observed among the most similar compounds to the tested compound. Validation sets were used to evaluate result accuracy by computing the Root Mean Square Error (RMSE) between the studied parameter’s experimental value and its final predicted value. Results with a reliability index under 0.3 (cut-off value) were discarded. Finally, 11 compounds were selected (**Supplementary Table 8**).

### Animals

We used RjOrl:SWISS mice (Janvier), OF1 mice (Charles Rivers; France), and *Pdgfra-CreERT*; *Rosa26stop-floxed-YFP* mice (RRID:IMSR_JAX:018280; RRID:IMSR_JAX:006148). Both females and males were included in the study. Mice were maintained under standard conditions with food and water available *ad libitum* in the ICM animal facilities. All animal studies were conducted following protocols approved by local ethical committees and French regulatory authorities (#03860.07, Committee CELYNE APAFIS#187 & 188, APAFIS#5516-2016053017288535, APAFIS #38705-2022092718027606).

### Neonatal Neural Progenitor Cultures

NPC proliferating medium: prepared using the following components: DMEM/F12 GlutaMAX™ supplement (life technology, 31331028), 0.6% glucose (Sigma, G8769), 1% penicillin/streptomycin (life technology, 15140122), 5mM HEPES buffer (life technology, 15630056), 1% N2 supplement (life technology, 17502048), 2% B27 supplement (life technology, 17504044), 20µg/ml Insulin (Sigma, I6634), 20ng/ml EGF (Peprotech, AF-100-15), and 10ng/ml FGF-basic (Peprotech, 100-18B).

#### Preparation and Maintenance of NPC Cultures

brains from RjOrl:SWISS mice (Janvier) at postnatal days 0-1 (P0-P1) were collected and washed three times in PBS-1X (Invitrogen, 14080055). Subventricular zones (SVZs) were dissected, transferred to fresh NPC medium, and dissociated using a pipette. Cells were amplified and maintained in a humidified atmosphere at 37 °C and 5% CO2. After 3 days, floating neurospheres were collected by centrifugation at 500g for 5 minutes, dissociated, and resuspended in proliferating NPC medium.

#### Plating and Treatment

after two passages, 30.000 cells were plated in 24-well plates with coverslips pre-coated with poly-L-ornithine (sigma, P4957). Compounds were added at concentrations of 250nM, 500nM, and 750nM. Positive controls were added including 3,3′,5-Triiodo-L-thyronine sodium salt (Sigma, T6397) at 30mM and clemastine fumarate (Sigma, SML0445) at 500nM with their respective vehicle (DMSO and PBS). The medium (including compounds or vehicles) was changed every 2 days.

#### Differentiation and Fixation

after 4 days in NPC proliferating medium, a differentiation medium (without supplemented growth factors) was added for two days. Cells were then washed once in PBS-1X, fixed for 10 minutes in 4% PBS-paraformaldehyde (Electron Microscopy Sciences, 50-980-495), and washed three times in PBS-1X.

### Oligodendrocyte Precursor Cell Sorting and Cultures

#### Brain Dissection and Cell Sorting

brains from P4 mice, cortices, and corpora callosa were dissected and dissociated using a neural tissue dissociation kit (P) (Miltenyi Biotec, 130-092-628) with the gentleMACS Octo Dissociator (Miltenyi Biotec, 130-096-427). OPCs were isolated via magnetic cell sorting (MACS) using anti-PDGFRα-coupled-beads (CD140a-PDGFRα MicroBead Kit, Miltenyi biotec, 130-101-502) and the MultiMACS Cell24 Separator Plus (Miltenyi biotec, 130-095-691).

#### Culture Conditions

40,000 cells were plated on coverslips pre-coated with poly-L-ornithine (sigma, P4957) and amplified for two/three days in a proliferating medium containing DMEM High Glucose (Dutscher, L0103-500), Ham′s Nutrient Mixture F12 (Merck, 51651C), L-Glutamine (Thermo, 200 mM), Hormone mix (Gritti, A., et al., 2001), Penicillin/Streptomycin (life technology, 15140122, 1X), 20ng/ml EGF (Peprotech, AF-100-15), 10ng/ml FGF-basic (Peprotech, 100-18B), and 10ng/ml PDGF-AA (PeproTech, 100-13A). Differentiation and Treatment: following the proliferation phase, cells were shifted to the differentiation medium for 3 days (without growth factors). Compounds were added to the medium at a concentration of 750nM. Positive controls including T3 (3,3′,5-Triiodo-L-thyronine sodium salt, Sigma, T6397) at 30nM and clemastine fumarate (Sigma, SML0445) at 500nM were used as positive controls, with their respective vehicles (DMSO and PBS) as negative controls. Cells were fixed for 10 minutes in 4% PBS-paraformaldehyde (Electron Microscopy Sciences, 50-980-495) and washed three times with PBS-1X.

### Cerebellar organotypic cultures

#### Culture Preparation

cerebellar organotypic cultures were prepared from newborn (P0) RjOrl:SWISS mice as previously described (Bouslama-Oueghlani et al., 2003). Briefly, cerebellar parasagittal slices (350 μm thick) were cut on a McIlwain tissue chopper and transferred onto 30 mm diameter Millipore culture inserts with 0,4 µm pores (Millicell, Millipore, PIHP03050).

#### Culture Conditions

slices were maintained in incubators at 37°C, under a humidified atmosphere containing 5% CO2 in six-well plates containing 1 ml of slice culture medium, containing basal Earle’s salts medium (BME, sigma, MFCD00217343), 25% Hanks’ balanced salt solution (Sigma, H9394), 27mM glucose (Sigma, G8769), 1% penicillin/streptomycin (life technology, 15140122), 1mM glutamine (Sigma, 228034), and 5% horse serum (New Zealand origin, heat-inactivated; ThermoFisher, 16050122). The medium was renewed every 2 to 7 days to support OPC differentiation and myelination.

#### Treatment and Fixation

at this time point, small molecules were added daily for 3 days, including Sm1, Sm2, Sm5, Sm6, Sm7, and Sm11 at 750nM. Positive controls including triiodothyronine (T3; Sigma, T6397-100MG) at 30nM and clemastine fumarate (SelleckChem, Houston, TX) at 500nM were added, with their respective vehicles (DMSO and PBS) as negative controls. After treatment, slices were collected and fixed for 1 hour at room temperature in 4% PBS-paraformaldehyde (Electron Microscopy Sciences, 50-980-495) and incubated for 20 minutes at 4°C in Clark’s solution (95% ethanol/5% acetic acid), then washed 3 times with PBS.

### Compounds nomenclature

Small molecule (Sm)

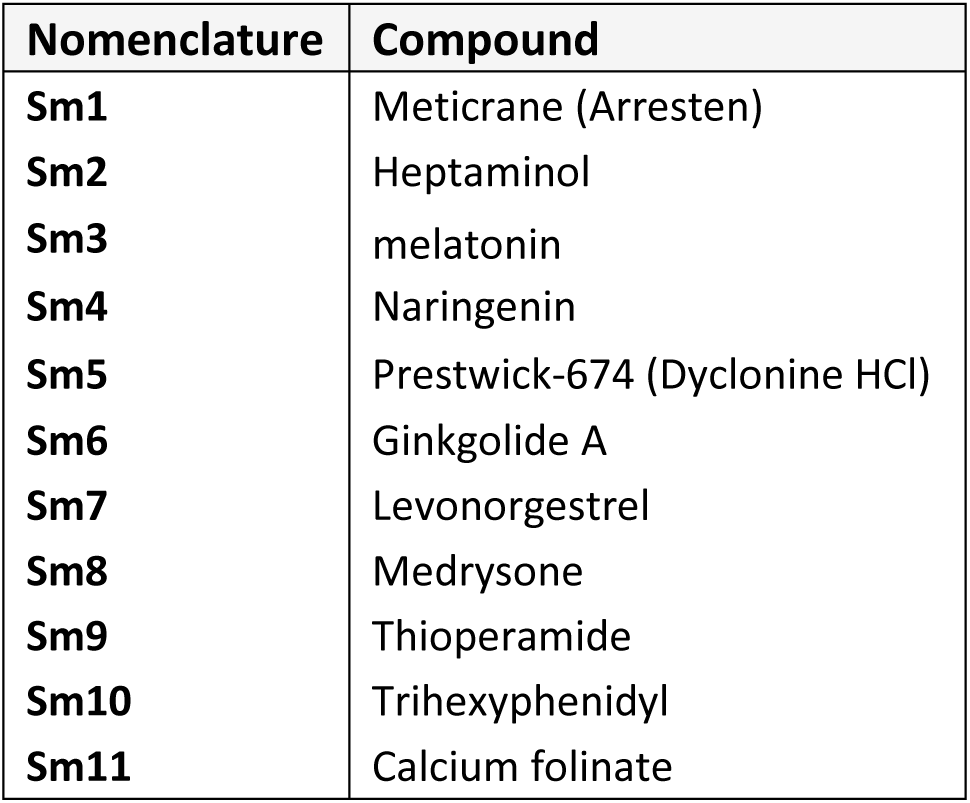

### Neonatal Hypoxia

#### Hypoxic Rearing Conditions

P3 OF1 mice were placed for 8 days until P11 in a hypoxic rearing chamber maintained at 10% O2 concentration by displacement with N2 as previously described (Fagel et al., 2005). A separate group was maintained in a normal atmosphere (normoxic group).

#### Compound Administration

compounds were administered intranasally, with doses corresponding to the oral administration in the adult mice Dyclonine/Sm5 at 5 mg/kg (Okazaki *et al*., 2018; Sahdeo *et al*., 2014) and Leucovorin/Sm11 at 0,5 mg/kg in humans (Frye *et al*., 2018; Rossignol and Frye, 2021). Mucus was first permeabilized using type IV hyaluronidase, then 10 μl of compounds (Sigma) were administered 3 times daily from P11 to P13 (starting at the end of the hypoxic period and repeated every 24 hours) in sterile PBS (control).

#### Proliferation and Maturation Analysis

for proliferation analysis, mice were injected with EdU (Sigma) to label cells in the S-phase of the cell cycle. Mice were perfused 1 hour post-injection at P13. For analysis of OL maturation, mice were perfused at P19.

#### Perfusion and Tissue Preparation

All perfusions were performed with Ringer’s solution, followed by an ice-cold solution of 4% paraformaldehyde (Thermo Fisher). Mice were sacrificed at P13 or P19 by an intraperitoneal overdose of pentobarbital, followed by perfusion with Ringer’s lactate solution and 4% paraformaldehyde (PFA; Sigma) dissolved in 0.1M phosphate buffer (PB; pH 7.4). Brains were removed and post-fixed for 24 hours at 4°C in 4% PFA and vibratome-sectioned in 50-µm thick coronal serial free-floating sections.

### Intracerebral Demyelination in the Adult Mouse

#### Induction of Recombination in Adult OPCs

tamoxifen (Sigma) was administered orally (gavage) to *Pdgfra-CreERT; Rosa26stop-YFP* mice for five consecutive days (210 g/kg per day, dissolved in corn oil at 20 mg/ml), 10 days before lesion induction to optimize the YFP-labelling of newly-generated OLs in the lesion area.

#### Lesion Induction

before surgery, 4-month-old RjOrl:SWISS mice (Janvier) and *Pdgfra-CreERT; Rosa26stop-YFP* mice were administered with buprenorphine (30 mg/g) to prevent postsurgical pain. Mice were anesthetized with isoflurane (ISO-VET). Eye protection (Ocrygel, Tvm) and cream lidocaine (Anesderm 5%) were applied to prevent eye dryness and pain from ear bars. A small incision was made on the head, and liquid lidocaine was applied to the site. Focal demyelinated lesions were induced by injecting 0.5μl of a 1% lysolecithin solution (L-a-lysophosphatidylcholine, Sigma L4129) diluted in 0.9% NaCl into the corpus callosum. A glass capillary connected to a 10μl Hamilton syringe was fixed and oriented through stereotaxic apparatus (coordinates: 1 mm lateral, 1.3 mm rostral to Bregma, 1.7mm deep to brain surface). Animals were left to recover for a few hours in a warm chamber.

#### Compound Administration

Dyclonine/Sm5 (5 mg/kg) and Leucovorin/Sm11 (0.5 mg/kg) were administered daily in 5% glucose drinking water according to the reported oral administration in adult mice (dyclonine; (Okazaki *et al*., 2018; Sahdeo *et al*., 2014)) and humans (leucovorin; (Frye *et al*., 2018; Rossignol and Frye, 2021)). Given the reported solubility in water and the half-life of these compounds, we diluted them in the drinking water and renewed the treatment daily. Given that a 40 g mouse drinks approximately 6mL per day, we diluted respectively 0.2 mg of dyclonine/Sm5 (0.2 mg in 40 g = 5 mg/kg) and 0.02 mg of leucovorin/Sm11 (0.02 mg in 40 g = 0.5 mg/kg) in 6mL, scaling these drug concentrations to a total volume of 200mL per bottle. We added 5% glucose to the preparation to increase the appetite of the mice.

#### Monitoring

Daily intake volumes were monitored, ensuring no significant differences in consumption between treatments, and confirming the appropriate dosing.

#### Perfusion and Tissue Preparation

mice were perfused 7 dpI and 10 dpl with 25 mL of 2% PBS-paraformaldehyde freshly prepared from 32% PFA solution (Electron Microscopy Sciences, 50-980-495). Perfused brains were dissected out, and dehydrated in 10% sucrose, followed by 20% sucrose overnight. Brains were then embedded in OCT (BDH), frozen, and sectioned into 14-µm thick sagittal sections using a cryostat microtome (Leica).

### Immunofluorescence Staining

#### Coated Coverslips

coverslips were blocked for 30 min at room temperature in a moist chamber using a solution of 0.05% Triton X-100 and 10% normal goat serum (Eurobio, CAECHVOO-OU) in PBS. Coverslips were incubated with primary antibodies for 30 min at room temperature. Following 3 washes with 0.05% Triton X-100/PBS, coverslips were incubated for 30 min at room temperature with secondary antibodies (Molecular Probes or Thermo Fisher) and DAPI (Sigma-Aldrich®, D9542). Coverslips were washed 3 times with 0.05% Triton X-100 in PBS and mounted with Fluoromount-G® (SouthernBiotech, Inc. 15586276).

#### *Ex Vivo* Cerebellar Slices

slices were incubated for 20 min at 4°C in Clark’s solution (95% ethanol/5% acetic acid) and washed 3 times with PBS. Then, sections were incubated for 1 hour in a blocking solution (0.2% Triton X-100, 4% bovine serum albumin, and 4% donkey serum in PBS). Slices were incubated with primary antibodies for 2 hours at room temperature. Slices were washed 3 times with 0.1% Triton X-100 in PBS and incubated with secondary antibodies (Molecular Probes or Thermo Fisher) and DAPI (Sigma-Aldrich®, D9542) in blocking solution at room temperature for 2 hours. Slices were then washed 3 times with 0.1% Triton X-100 in PBS and mounted in Fluoromount-G® (SouthernBiotech, Inc. 15586276).

#### *In vivo* Cryosections of Lesions

sections (14-μm thick) of lesions were dried for 30 min at room temperature. For MOG staining, sections were treated with 100% ethanol for 10 minutes at room temperature. Sections were permeabilized and blocked in a solution of 0.05% Triton X-100 and 10% normal goat serum in PBS for 1 hour. Sections were incubated overnight with primary antibodies at 4°C. Sections were washed 3 times with 0.05% Triton X-100 in PBS and incubated with secondary antibodies (Molecular Probes or Thermo Fisher) and DAPI (Sigma-Aldrich®, D9542) during 1 hour at room temperature. Sections were washed 3 times with 0.05% Triton X-100 in PBS and mounted with Fluoromount-G® (SouthernBiotech, Inc. 15586276).

#### Forebrain Sections Following Neonatal Hypoxia

free-floating 50µm coronal sections were collected. Antigen retrieval was performed for 20 minutes in citrate buffer (pH 6.0) at 80°C, cooled for 20 minutes at room temperature, and washed with PBS. Blocking was performed in a TNB buffer (0.1 M PB; 0.05% Casein; 0.25% Bovine Serum Albumin; 0.25% TopBlock) with 0.4% triton-X (TNB-Tx). Sections were incubated with primary antibodies overnight at 4°C under constant agitation. Following washing in 0.1 M PBS? with 0.4% triton-X (PB-Tx), sections were incubated with secondary antibodies (Jackson or Invitrogen) and DAPI (Life Technologies; D1306) for 2 hours at room temperature. The revelation of EdU was done using Click-it, EdU cell proliferation Kit (Thermo Fisher Scientific).

The list of primary and secondary antibodies used can be found in Methods **Tables 4 and 5**.

### Image Acquisition and Analysis

#### Visualization and Acquisition

cryosections and immunocytochemistry were imaged using Zeiss® Axio Imager M2 microscope with Zeiss® Apotome system at 20X/0.8 NA dry objective (Plan-apochromat), including deconvolution and Z-stack. Acquisition for cerebellar slices was done using a confocal SP8 X white light laser (Leica), at ×40 magnification with a Z-axis range of 10–15 μm, and nine fields (1024 × 1024) were captured and merged into a mosaic.

#### Quantifications

for neonatal hypoxia sections, quantification was performed by automatic quantifications of Olig2 signal using QuPath software (V0.3.0) in manually defined regions of the corpus callosum and cortex (based on DAPI counterstaining). Other analyses were performed using ZEN (Zeiss) and ImageJ (Fiji) software packages from z-stack images, with macros for *ex vivo* experiments to determine myelination and differentiation index (methods previously described in (Baudouin *et al*., 2021). Olig2+ cells were detected in five equally spaced coronal sections (50 µm, 250 µm apart). Identification of Olig2+ cells co-expressing GSTπ was performed in manually defined regions covering the entire forebrain (Isocortex, Olfactory areas, Striatum, Fiber tracts, Cortical subplate, Thalamus, Hypothalamus) using the Allen adult mouse brain reference atlas (CCF v3). Manual quantification was done on z-stack confocal images acquired from the hypothalamic lateral zone, using ImageJ-Win-64. *In vivo,* images of lesions were quantified manually in a double-blinded manner. Illustrations were performed using Illustrator and Adobe Photoshop (Adobe System, Inc). For electron microscopy analysis, axons were quantified using Fiji software packages.

### Electron Microscopy

#### Sample Preparation

mice were perfused with 2.5% glutaraldehyde in 0.1M sodium cacodylate buffer 0.1M (Caco buffer) at pH 7.4. Brains were collected and left overnight in the fixative at 4°C. After rinsing in Caco buffer, brains were sliced into 1mm-thick sections. Sections were selected and rinsed 3 times with Caco buffer and post-fixed with 2% osmium tetroxide Caco buffer for 1h at room temperature. After an extensive wash (3×10 min) with distilled water, they were incubated overnight in 2% aqueous uranyl acetate. Dehydration was done in a graded series of ethanol solutions (2×5min each): 30%, 50%, 70%, 80%, 90%, and 100% (X3). Final dehydration was performed twice in 100% acetone for 20 min. Samples were then progressively infiltrated with an epoxy resin, Epon 812® (EMS, Souffelweyersheim, France) for 2 nights in 50% resin 50% acetone at 4°C in an airtight container, then twice for 2h in pure fresh resin at room temperature. They were embedded in molds and put at 56°C for 48h in a dry oven for polymerization of the resin. Blocks were cut with a UC7 ultramicrotome (Leica Microsystemes). Semi-thin sections (0.5μm thick) were stained with 1% toluidine blue and 1% borax. Ultra-thin sections (70nm thick) of the region were recovered on copper grids (200 mesh, EMS, Souffelweyersheim) and contrasted with Reynold’s lead citrate (Reynolds, ES (1963)).

#### Microscopy

ultrathin sections were observed with a Hitachi HT7700 electron microscope (Milexiaé) operating at 100 kV. Images (2048×2048 pixels) were taken with an AMT41B camera.

### Statistics and Representations

#### Software and Analysis

all analyses were performed using GraphPad Prism 6.00 (San Diego California, www.graphpad.com). For electron microscopy, plot visualizations, and statistics were performed in R. Scripts used are available on demand.

#### Statistics

The assumption of normality was made for data distribution; formal testing for normality was not performed. Statistical significance was determined using One-Way ANOVA by multiple comparisons to evaluate differences among groups. Dunnett’s Post-hoc test was used following one-way ANOVA for pairwise comparisons against a control group, and unpaired t-tests were used for comparisons between two independent groups. Data are presented as Mean ± SEM (Standard Error of the Mean). Significance levels were set as: p-value < 0.05: * (one asterisk), p < 0.01: ** (two asterisks), p < 0.001: *** (three asterisks), p < 0.0001: **** (four asterisks).

A summary of statistical tests used, the number of replicates, and p-values can be found in Methods **Table 3**.

## Acknowledgements

This work was supported by funding from the Agence Nationale de la Recherche [ANR NeoRepair (ANR-17-CE16-0009) and ANR NeoReGen (ANR-22-CE17-0029) grants], the French Multiple Sclerosis Foundation (ARSEP-1284), the ICM Carnot projects (#137, #142, and #148), and BDL Capital Management. The research leading to these results has received funding from the national program “Investissements d’avenir” ANR-10-IAIHU-0006 and the national program “Investissements d’avenir” ANR-11-INBS-0011 – NeurATRIS: Translational Research Infrastructure for Biotherapies in Neurosciences. OR lab is supported by the Fondation pour la Recherche Médicale (FRM: EQU202203014658) as well as by the LABEX CORTEX (ANR-11-LABX-0042) of Université de Lyon, within the program “Investissements d’Avenir” (decision n° 2019-ANR-LABX-02) operated by the French National Research Agency (ANR). We thank Severine Candelier and Mathilde Lannes for creating the OligoScore website, and François Xavier Lejeune for overseeing the statistical analysis. We thank the following ICM (Paris Brain Institute) core facilities: PHENOPARC, CELIS, Histomics, and ICM.QUANT, and all personnel involved for their contribution and support.

## Author contributions

Conceptualization, supervision, funding acquisition: C.P., and O.R. Design of the experiments: C.P., O.R., and J.-B.H. Achievement of the experiments and data analysis: bio-informatics, J.-H. B., C.P., O.R., and R.A.; culture models, J.-H. B., L.-M.G, C.M., L.B., and A.A.-N.; hypoxia model, L.B.-O.; L.F., N.V., and O.R.; adult demyelination model, J.-H. B., L.-M.G, C.P., C.M., N.I., A.A.-N., and M.P.; pharmacology, J.-A.C., M.-A.D., R.T., and F.G. Writing of the original version of the manuscript: C.P. with main contributions from J.-H.B, O.R., B.A.-H., and help from all other authors.

---

**Figure 1 extended data 1.**
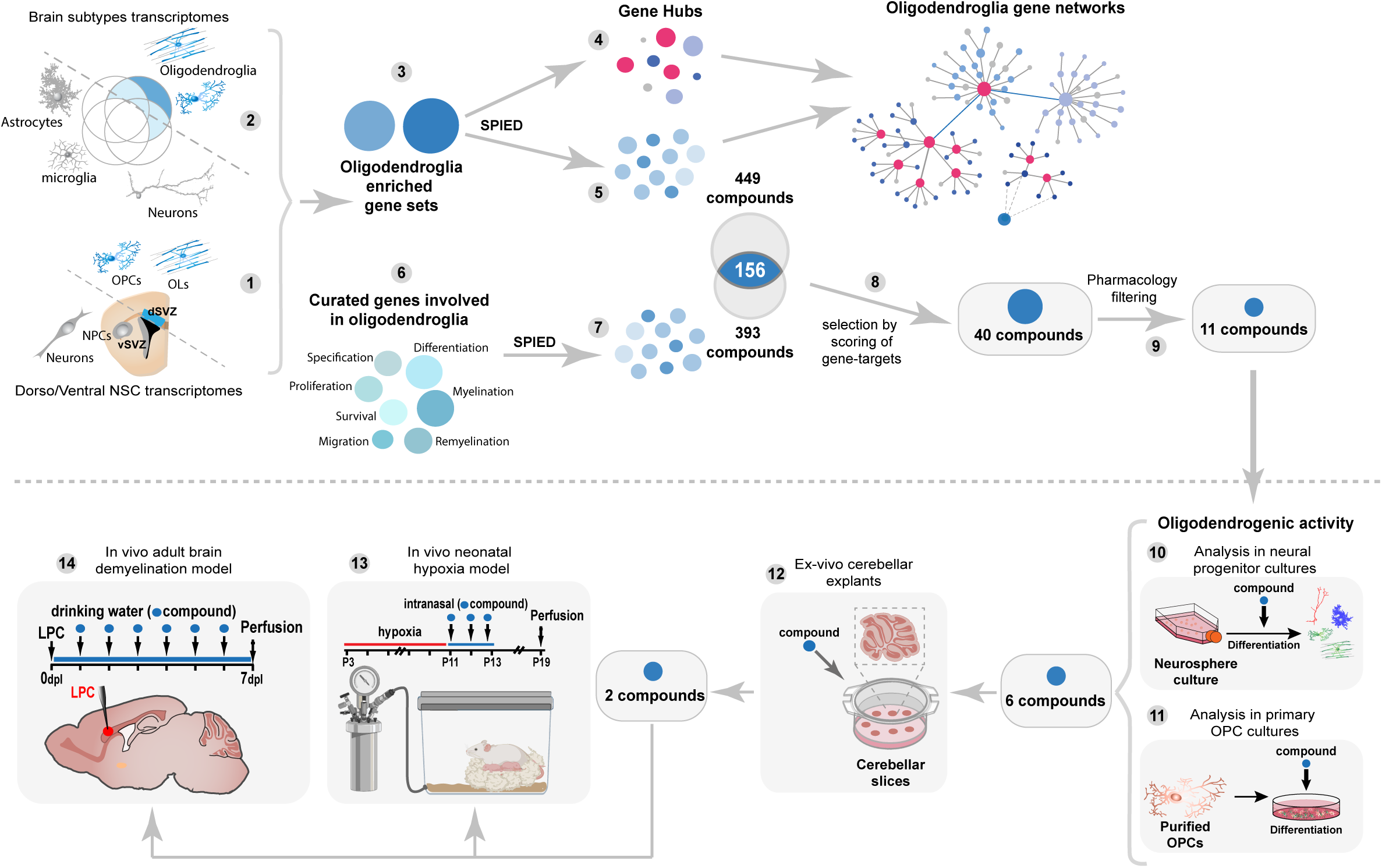
Summary scheme of the screening strategy and compound selection process. Schematics representing the in-silico (top) and wet-lab (bottom) strategies. In-silico bioinformatic strategy allowed the generation of a large oligodendroglial transcriptional signature (steps 1-3) and gene hubs of the oligodendroglial gene-networks (step 4) used to generate a list of compounds with putative pro-oligodendrogenic and (re)myelinating activities (step 5). Compounds were ranked by their large pro-oligodendrogenic activities through expert curation of their regulated target genes (steps 6-8), and then by their pharmacological properties (step 9), with a final selection of the top 11 compounds. Wet-lab strategy to assess the pro-oligodendrogenic activity of the selected compounds in the culture of neural progenitor cells (step 10) and OPCs (step 11), resulting in 6 compounds being selected for further activity assessment in *ex vivo* cerebellar explant cultures (step 12). Finally, the top two compounds were used *in vivo* in a model of preterm birth injury (neonatal hypoxia, step 13) and a model of adult de/remyelination (step 14). Numbers in grey circles indicate the different steps of the project as detailed in the main text.

**Figure 2 extended data 1.**
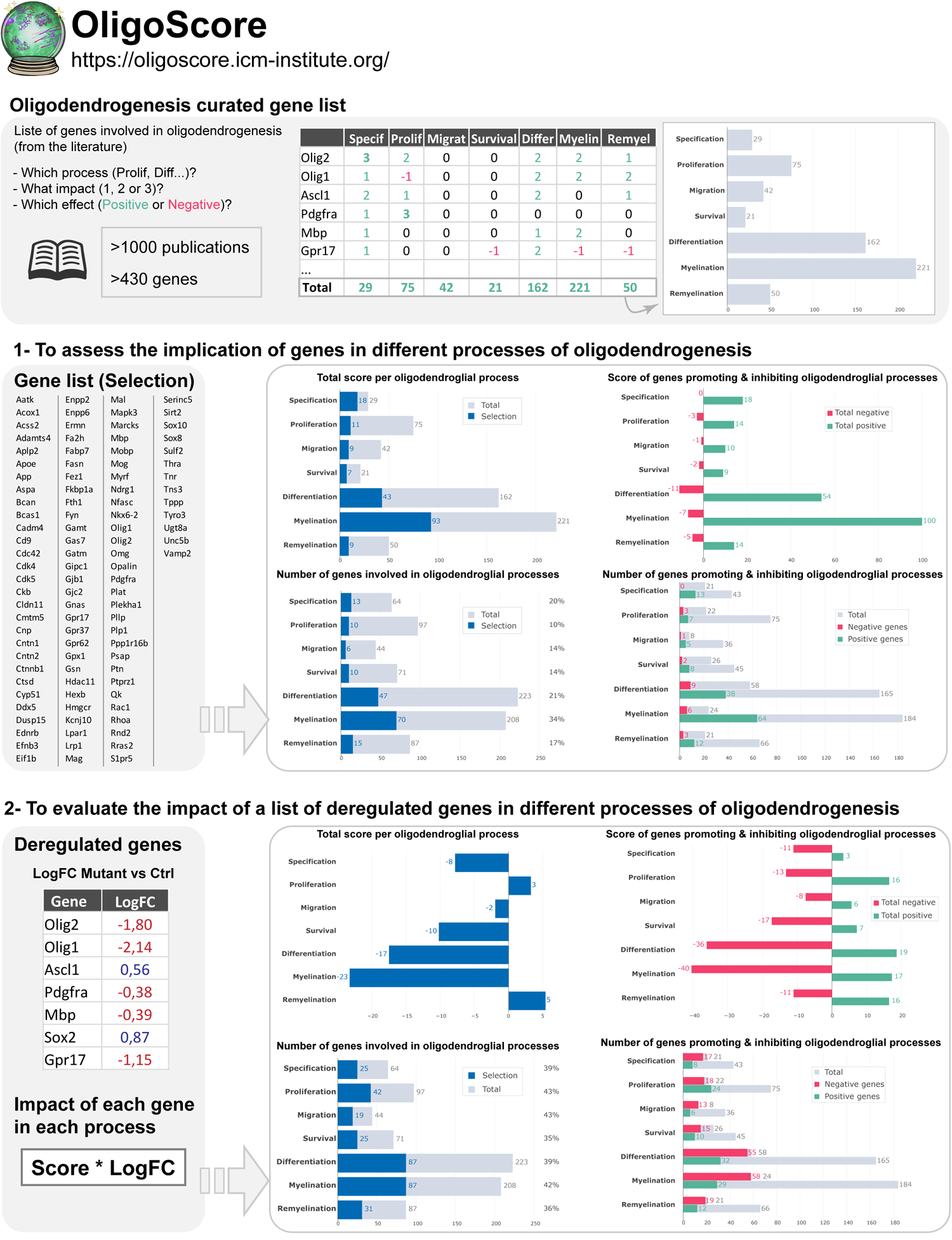
OligoScore, a new web resource to evaluate gene sets for their involvement in oligodendrogenesis. Schematics illustrating the OligoScore resource, a curation strategy currently encompassing over 430 genes and 1000 publications that report the functional implication (either by loss-of-function or gain-of-function experiments) of a given gene in each aspect (process) of oligodendrogenesis (i.e., specification, proliferation, migration, survival, differentiation, myelination, and remyelination). Detailed statistics obtained either from queries using either (1) a gene set (i.e, list of genes) or (2) a list of deregulated genes (i.e, genes and logarithmic fold-changes, logFC) are provided as barplots.

**Figure 2 extended data 2.**
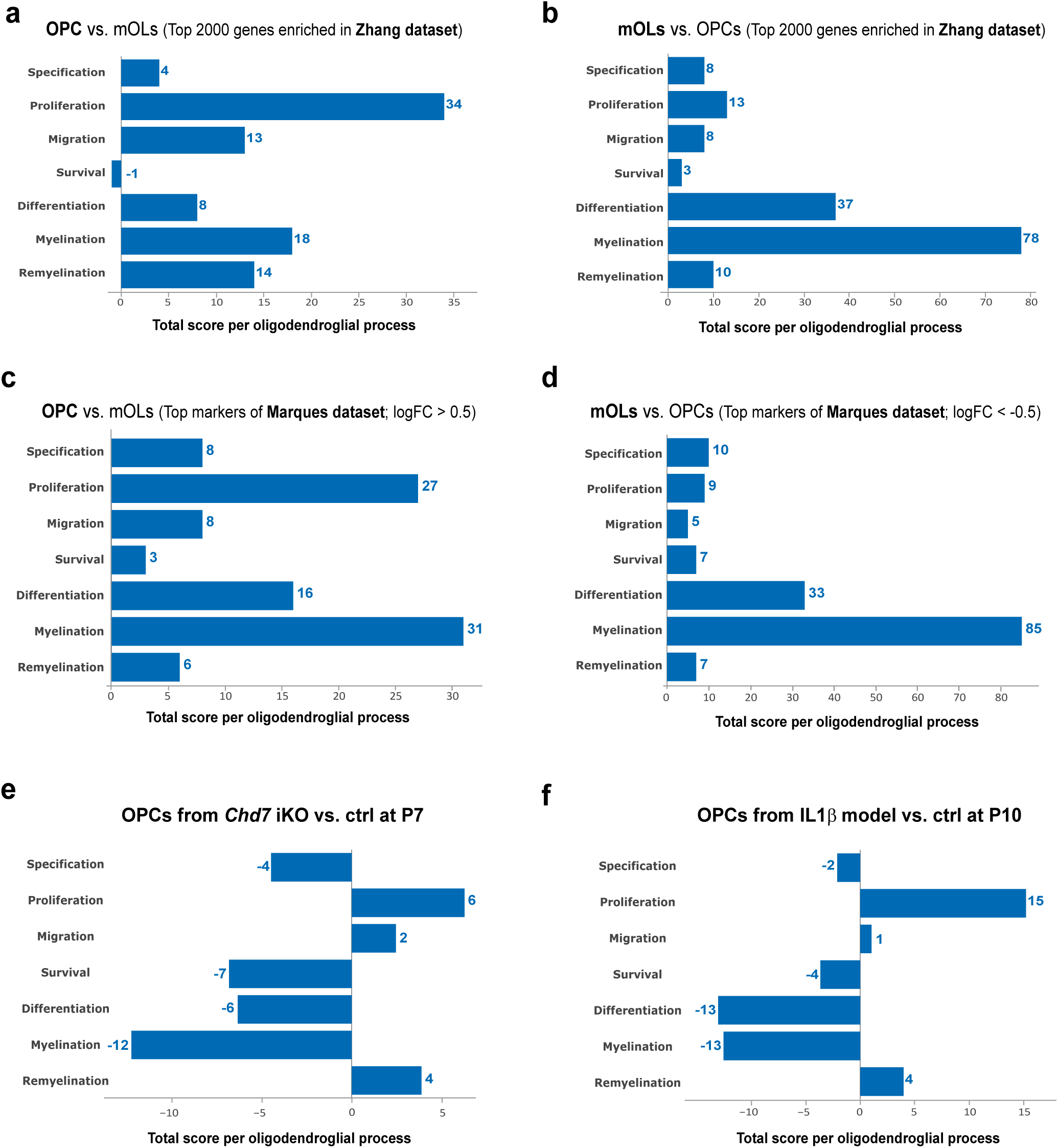
OligoScore validation using expression and deregulated transcriptional signatures. (**a-d**) Querying of OligoScore with genes enriched in OPC vs. myelinating OLs (mOLs), and vice versa, using either (**a**) postnatal purified brain cell transcriptomes (Zhang et al., 2024) or (**c**) oligodendroglial single-cells transcriptomes (Marques et al., 2016, 2018; see also Methods and *OligoScore_validation_tables* file). In both cases, OligoScore highlights that while OPC gene programs are involved in several processes of oligodendrogenesis, including specification, proliferation, migration, differentiation, and myelination (**a, c**), mOL-enriched genes are mainly involved in differentiation and myelination (**b, d**). (**e, f**) Querying OligoScore (genes and fold-changes) with genes dysregulated in OPCs upon a genetic (postnatal day 7 OPCs with an induced knockout for *Chd7,* compared to controls; Marie et al., 2018), or an environmental (P10 OPCs in the systemic IL1β-mediated neonatal neuroinflammatory model, Schang et al., 2022) perturbation. This analysis highlights the deregulated processes identified in these studies, such as reduced survival, and differentiation of *Chd7* iKO OPCs (**e**), and increased proliferation and reduced differentiation of OPCs in the IL1β-model (**f**).

**Figure 3 extended data 1.**
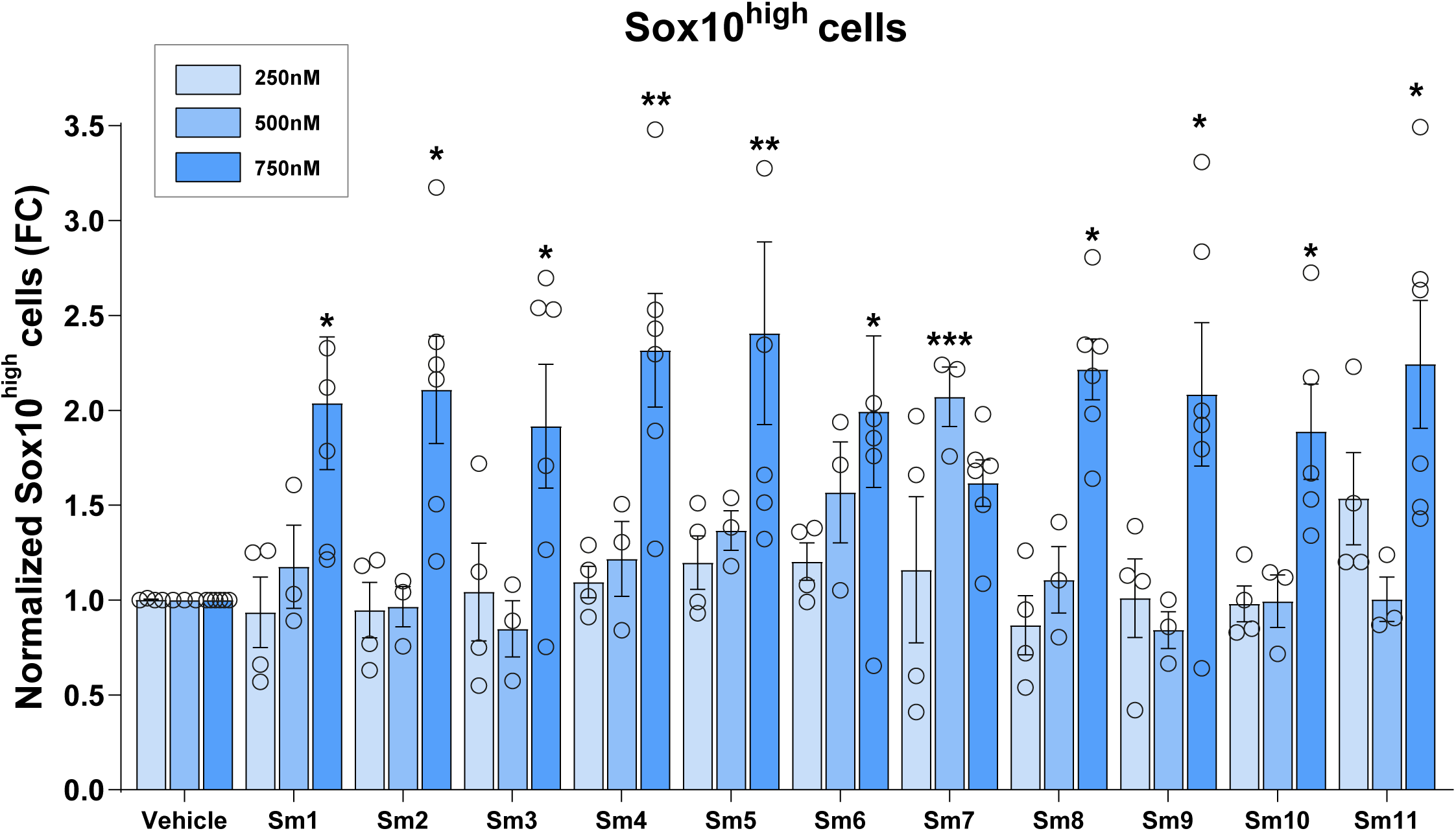
Dose effect of compounds’ pro-oligodendrogenic activity in neonatal neural progenitor cultures. Quantifications of So×10^high^ cells (iOLs) in cultures treated with compounds at 250nM (N=4), 500nM (N=3) and 750nM (N=6), showing the best effect at 750nM. Data are presented as mean ± SEM of fold change normalized to vehicle. *p < 0.05; **p < 0.01; ***p < 0.001. Statistical analysis used linear mixed-effects models followed by Type II Wald chi-square tests (ANOVA).

**Figure 4 extended data 1.**
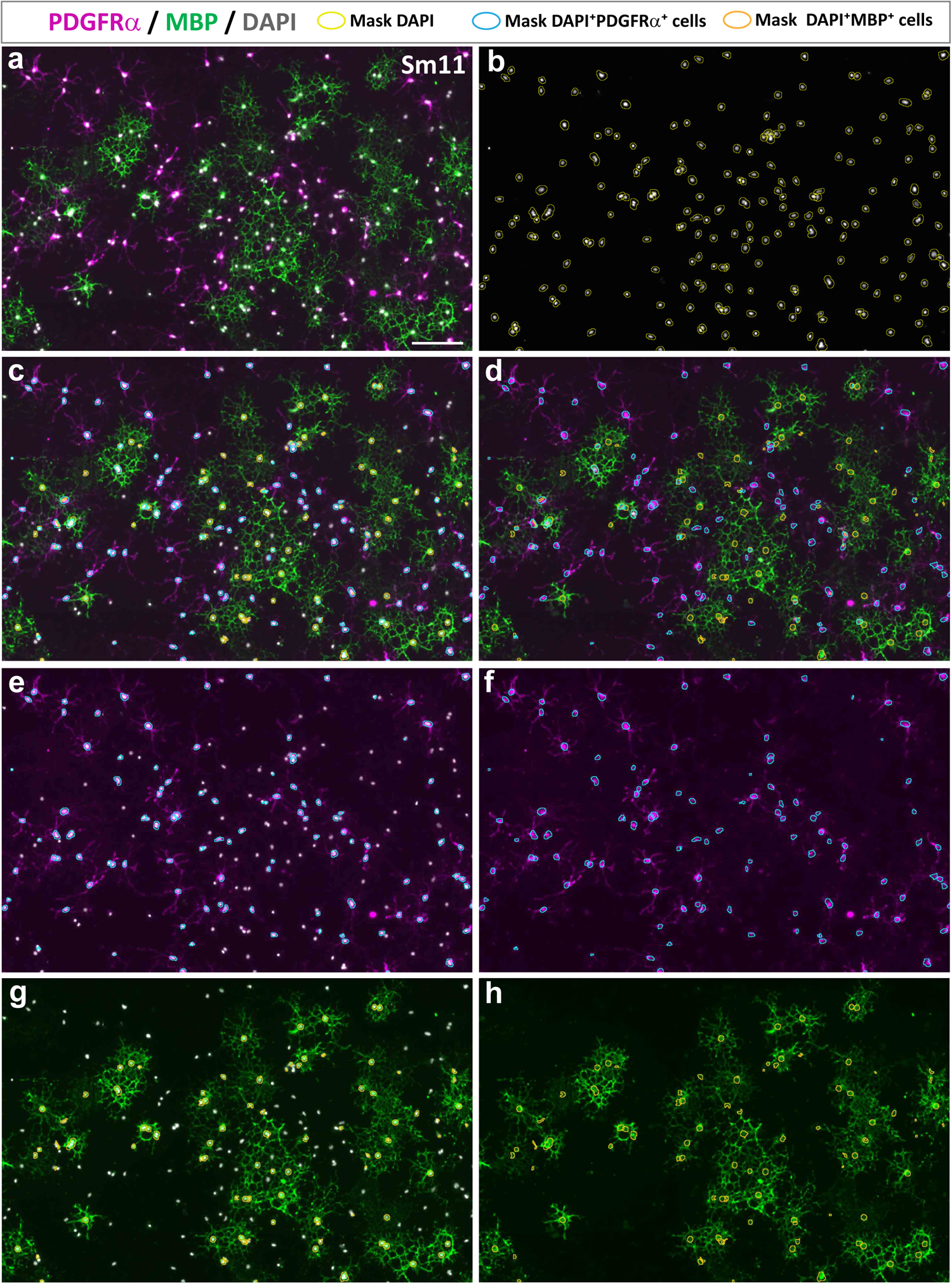
Automatic quantification of OPC differentiation cultures. (**a**) Representative image illustrating the immunodetection of OPCs (PDGFRα^+^ cells, magenta) and differentiating OLs (MBP^+^ cells, green) together with nuclear DAPI staining (grey) from OPC differentiation cultures, used to illustrate automatic segmentation and quantification shown in following panels. (**b**) Image illustrating with yellow masks the segmentation of DAPI nuclei (grey) used for automatic quantification. (**c-h**) Images illustrating with cyan masks the automatic identification of OPCs, DAPI nuclei having PDGFRα labeling (magenta), and with orange marks the identification of OLs, DAPI nuclei having MBP labeling (green), shown with (c, e, g) or without the DAPI channel (d, f, h), and showing PDGFRα and MBP labeling together (c, d) or as separate channels (e-h) to facilitate mask visualization.

**Figure 5 extended data 1.**
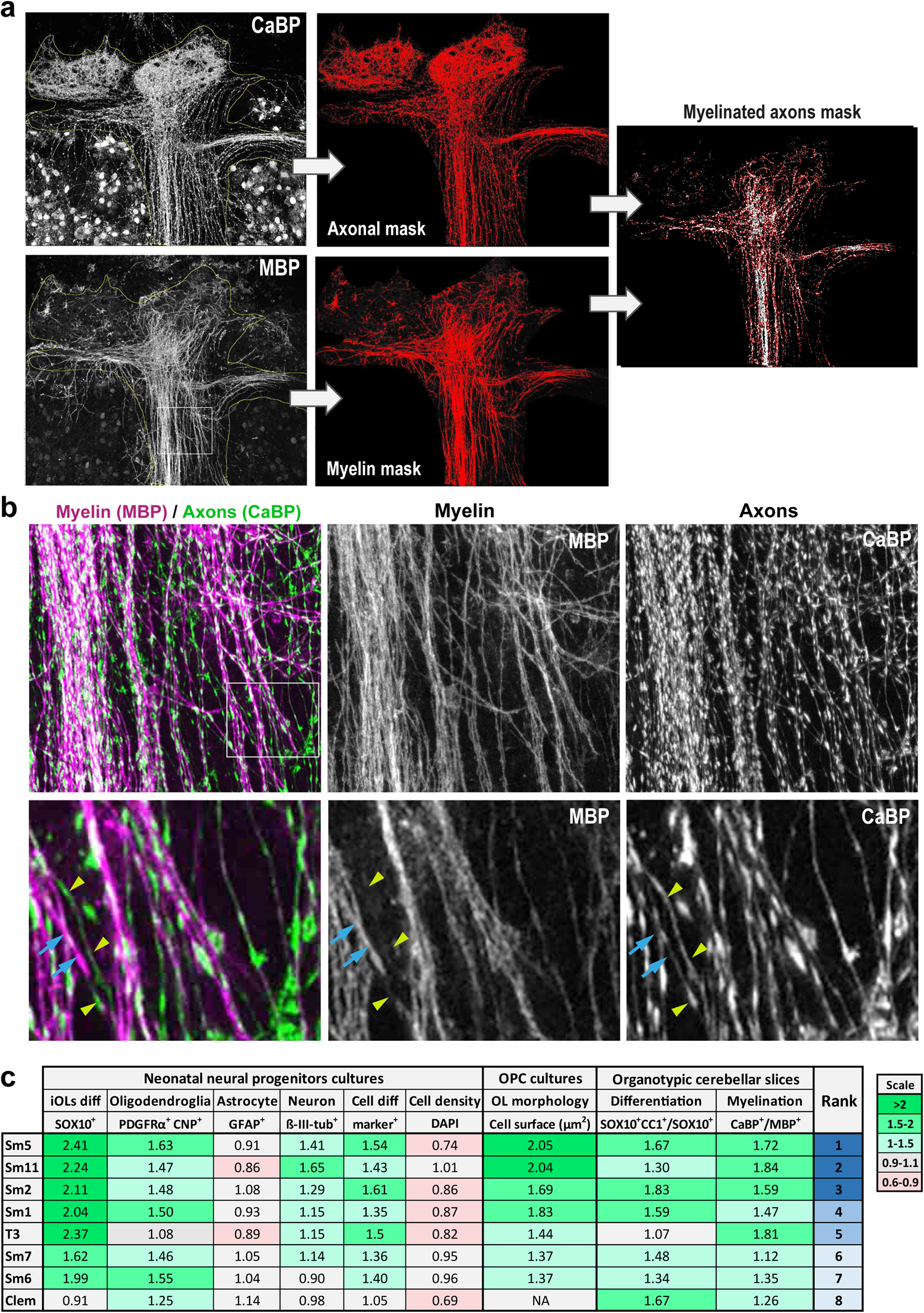
Automatic quantification of differentiation and myelination index in ex vivo cerebellar cultures, with a ranking of selected drugs for their pro-oligodendrogenic effects. (**a**) Illustration images of the automatic quantification approach to obtain the myelinating index (as optimized in Baudouin et al., 2021). (**b**) High magnification images illustrating the overlap between myelin, MBP immunodetection (magenta), and Purkinje neuronal axon immunodetected with CaBP antibody (green). Examples of myelinated-(blue arrows) and non-myelinated (yellow arrowheads) axonal segments are indicated in the lower panels. (**c**) Table integrating and ranking the effects of selected compounds on oligodendrogenesis. Biological effects of the small molecules and positive controls (T3, clem) determined through in vitro and ex vivo systems allow ranking them for their pro-oligodendrogenic activity to select the top two candidates for in vivo analysis. Data are presented as fold change compared to their respective vehicle. diff: differentiation; NA: not assessed; Clem, clemastine.

**Figure 6 extended data 1:**
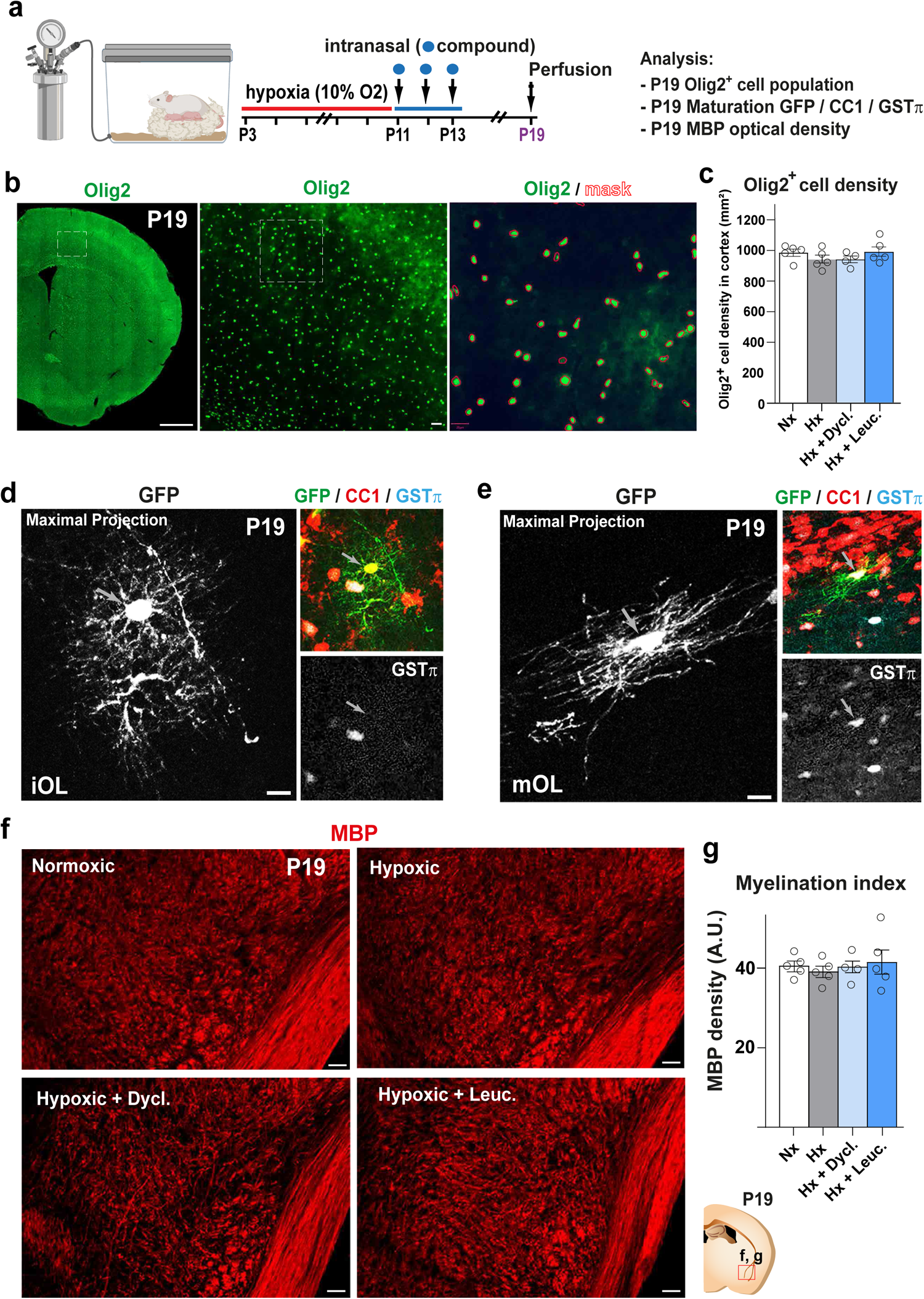
Dyclonine and leucovorin promote oligodendroglial regeneration in a mouse model of preterm birth brain injury. (**a**) Schematic illustrating the workflow used to assess the capacity of dyclonine and leucovorin to promote OPC proliferation and rescue OLs maturation following neonatal chronic hypoxia. (**b, c**) Olig2 immunodetection and quantification showing that the density of OLs is not changed by hypoxia nor by treatment within the cortex at P19 (**c**). Quantification was performed via automated detections (QPath) as illustrated by the mask used for cellular segmentation in the third illustration (mask). (**d, e**) Illustrations validating the use of CC1 and GST*π* markers to identify OLs and mature/myelinating OLs respectively. Note that during OL differentiation, immature OLs (labeled with GFP) present a stellar morphology and are positive for CC1 but not for GST*π* (**d**), while mature/myelinating OLs having myelin segments co-express CC1 and GST*π* (**e**). (**f, g**) Representative images illustrating myelination by MBP immunodetection in the hypothalamic lateral zone, showing no difference between groups at P19 (f) and quantification of MBP immunofluorescence in this zone is represented in arbitrary units (A.U.) (g). N≥ 4; Data are presented as Mean ± SEM. Scale bars: 100 µm in b left panel, 20 µm in b right panels, and f; e; 4 µm in d and e.

**Figure 7 Extended data 1.**
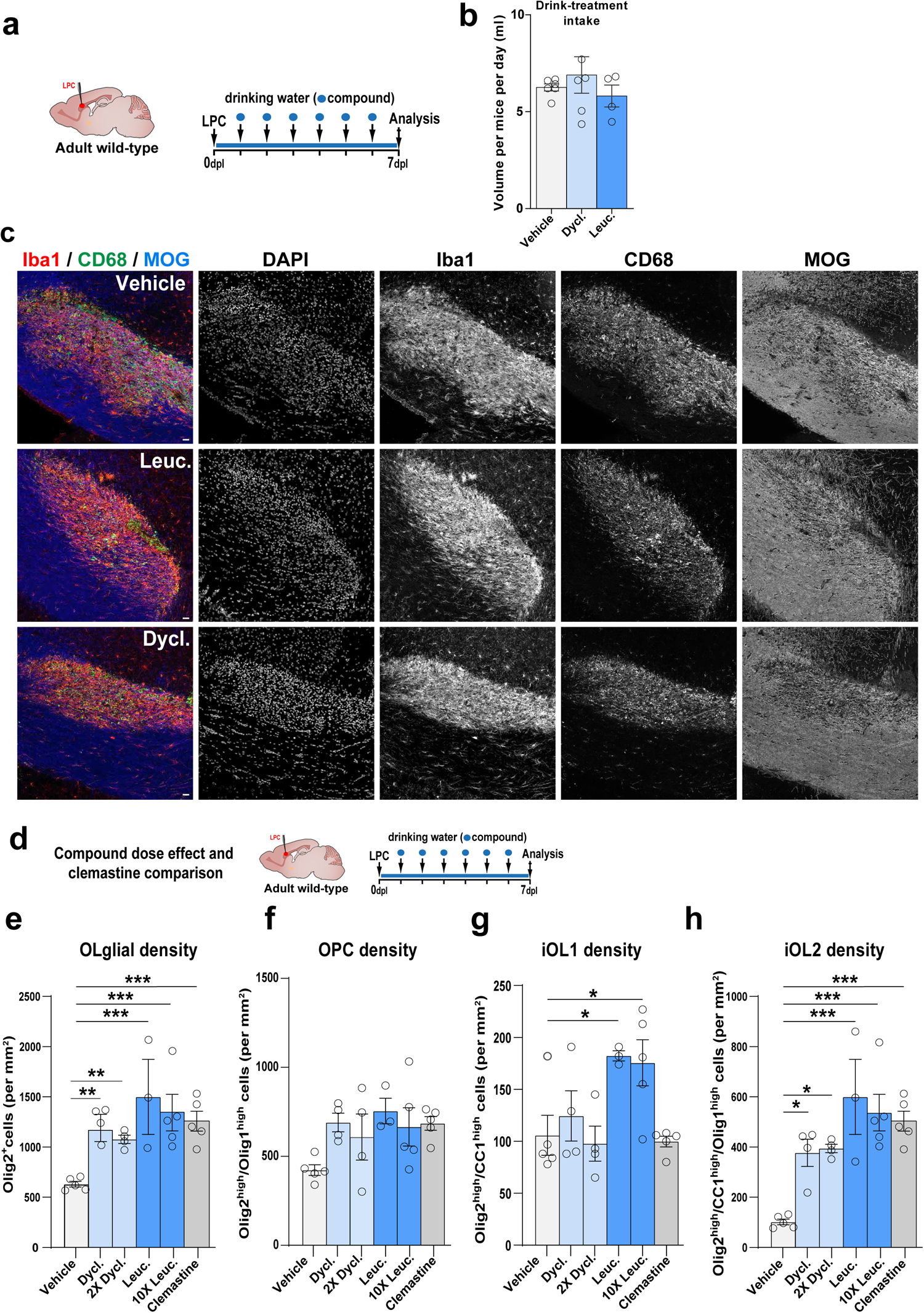
Dyclonine and leucovorin promote both OPC proliferation and differentiation in a mouse model of adult demyelination. (**a**) Schematics illustrating the protocol for LPC demyelination in the corpus callosum, the timing of compound administration in drinking water and analysis. (**b**) Quantification of drinking intake per day (ml) during treatment showing no difference between groups and thus confirming expected compound dose administration. (**c**) Single channel panels for the immunofluorescence in the lesion site identified by cell density (DAPI), microglia/macrophages (Iba1^+^ cells, red) and their phagocytic profile (CD68^+^ cells, green), and the reduction in myelin immunodetection (MOG, blue) in different treatment conditions. (**d**) Schematics illustrating the protocol of a replicated experiment to compare different doses of leucovorin and dyclonine, and clemastine as a pro-oligodendrogenic positive control. (**e-h**). Quantification of Olig2^+^ oligodendroglial density (**e**), OPC density (**f**), iOL1 density (**g**) and iOL2 density (**h**) showing a similar increase at both doses tested of leucovorin and dyclonine in the density of oligodendroglia (Olig2+ cells), iOL1s (only increased by leucovorin treatment), and iOL2s. Note that clemastine induces a similar effect to dyclonine, with only leucovorin increasing the density of iOL1s. Dycl., dyclonine; Leuc., leucovorin; 2X Dycl. 2-fold dyclonine dose; 10X Leuc., 10-fold leucovorin dose; Clem., clemastine. Scale bars: 20 μm.

**Figure 8 Extended data 1.**
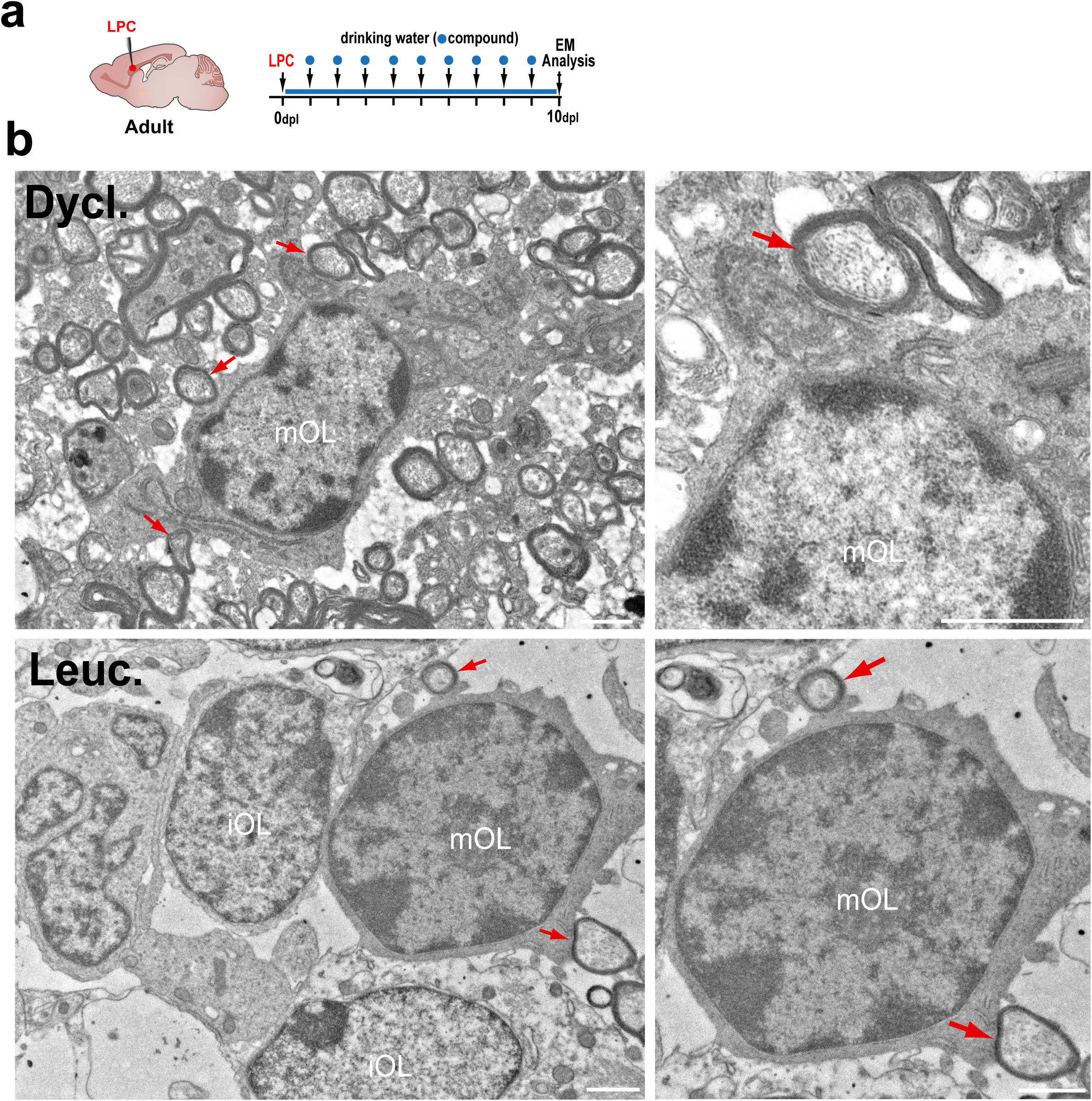
Leucovorin and dyclonine accelerate the generation of myelinating oligodendrocytes in a mouse model of adult demyelination. **(a)** Schematics illustrating the protocol used for compounds’ administration in the drinking water and ultrastructural analysis by EM at 10 days post-lesion (dpl). **(b)** High magnification micrographs illustrating newly formed (re)myelinating OLs (mOL) in the lesion area, identified by their typical ultrastructural traits, i.e., oval-shape nucleus having densely packed chromatin with several heterochromatin spots and large cytoplasm in continuity with axons presenting compact myelin ultrastructure (red arrows) in dyclonine- and leucovorin-treated animals. Two examples of immature OLs (iOL), characterized by less compacted chromatin (lighter nuclear contrast than mOLs) and large cytoplasmic processes, are also shown in the bottom-left panel. The right panels show higher magnifications to better visualize the continuity between the OL cytoplasm and the myelinated axons (red arrows). Dycl., dyclonine; Leuc., leucovorin. Scale bars: 1 μm.

## Point by point response to reviewers’ comments (1^st^ round)

### Answers in blue

#### Reviewers’ comments

##### Reviewer #1 (Remarks to the Author)

This manuscript aimed to find new therapeutic solution for some very challenging diseases, namely, early preterm-birth brain injury and adult multiple sclerosis. **This is very clinically important**. The authors c**reatively proposed a very novel strategy** to identify the drug targets for these diseases by the **full utilization of genetic data and bioinformatics**. These efforts indeed generate some potential lead compounds, and their effect were subsequently proved by both in vitro and in vivo experiments. Generally, this study has **very good creativity** which was supported by **solid experiment data**. The findings of this manuscript **could also provide useful hints for the development of new medication of early preterm-birth brain injury and adult multiple sclerosis**. Whereas the quality of this manuscript could be further improved in the following aspects.

We thank the reviewer for her/his enthusiastic comments concerning the clinical relevance, creativity, and solidity of the experimental data of our study. We answer to the point raised below.

1) Line 153, the authors developed a very novel scoring system useful for the identification of effective compounds using the genetic data in this study. I am wondering whether this scoring system has been validated somewhere else before it was employed in this study.

To answer the reviewer: (1) in the context of a recent collaboration (Nat. Commun., DOI: 10.1038/s41467-023-36846-w), we used OligoScore to analyze bulk RNA-seq data from spinal cord of EAE mice subjected to different treatments, not only confirming at the transcriptional level the deregulated processes of oligodendrogenesis previously found by immunofluorescent analysis, but also identifying key genes responsible for these deregulations; (2) in our revised manuscript, we added two further validations. First, we queried OligoScore with genes enriched in OPCs and myelinating OLs (mOLs), using transcriptomes from purified brain cells (Zhang et al., 2014, RNAseq) or from single cells (Marques et al., 2016, 2018). With both datasets, as expected, we found that genes enriched in OPCs are involved in several processes of oligodendrogenesis (specification, promoting proliferation, differentiation, and (re)myelination), while genes enriched in mOLs are mainly promoting differentiation and myelination (Figure 2 extended data 2a-d). Second, we queried OligoScore with genes deregulated in OPCs upon either a genetic (*Chd7*-deletion in OPCs; Marie et al., PNAS 2018) or environmental perturbation (IL1β-mediated neonatal neuroinflammatory model, Schang et al., 2022). Remarkably, OligoScore analysis not only highlighted the deregulated processes of oligodendrogenesis previously found by immunofluorescent analyses in these studies (that is, reduced survival and differentiation of *Chd7-iKO* OPCs, and reduced OPC differentiation paralleled by an increase proliferation of OPCs from IL1β-induced neuroinflammatory model), but also provided mechanistic insights identifying some of the genes responsible for the deregulation of each process. We have added these analyses to the manuscript as follows: (a) results section lines 165-170 and Figure 2 Extended Data 2, (b) gene sets used to query and validate OligoScore in the Excel file named ‘OligoScore validation tables’, (c) a method’s section named ‘Validation of OligoScore strategy’ describing these validations.

2) Line 177, the authors ranked the potential of compounds as a lead compound by evaluate various properties of druggability such as toxicology, pharmacokinetics and pharmacodynamics. This is a correct and smart strategy. After checking Supplementary table 8, we can see that the authors have paid too much attention on the issue of toxicology. Most of evaluation items in Supplementary table 8 are related to toxicology rather than pharmacokinetics and pharmacodynamics. In my opinion, the identification of suitable candidates for the proof of concept should focus on their pharmacokinetics and pharmacodynamics characteristics. However, I found very few items of scoring related to pharmacokinetics and pharmacodynamics characteristics. Even an important scoring item related to pharmacokinetic of CNS drug, namely, BBB penetration is just a very simple qualitative prediction provided by the software of ACDsee. Actually, many of the compounds in Supplementary table 8 are old drugs. I guess it is possible to collect the BBB penetration data in animals from the literature. Such quantitative information may be more valuable for such scoring purpose.

According to the reviewer’s suggestion, we searched the literature for those compounds experimentally tested for their BBB penetration and found data supporting most of the predictions (83%, 23 out of 28 molecules). For the 10 molecules with no experimental data, we have now labeled them as ‘(Predicted)’. We included this data and the bibliographic references for the 28 molecules with available experimental values in the supplemental table S8 columns D and E, and provide the details in the Word file named *Reference S8 BBB*.

In more general aspects, we selected compounds based on: (1) their larger pro-oligodendrogenic activities (high pharmacogenomic total scores), (2) their crossing of the BBB, and (3) not having major predicted toxicity. Finally, for the *in vivo* studies, we selected the best two being FDA-approved, so that their pharmacokinetics and pharmacodynamics did not need to be repeated. We have modified the main text to make this point clear in **lines 180-186**.

3) Line 621, please specify the names of the compounds here. Please also clarify the compounds information in other parts of this manuscript such as line 300.

As suggested by the reviewer in our revised manuscript we have added a Method section called ‘Compounds nomenclature’ and added the dyclonine (Sm5) and leucovorin (Sm11) in the text of the *in vivo* studies.

4) Line 637, in the animal experiment, the authors used a dose of 30 mg/g buprenorphine. This is a very high dose considering the human dose is usually less than 1 mg. How can the mice tolerate such a high dose in the experiment? Additionally, the oral bioavailability of buprenorphine is low due to the extensive first pass effect. Thus this drug is administered by injection or sublingual administration. The authors didn’t describe the administration method. This should be clearly described.

We thank the reviewer for spotting this error in the methods which we have corrected. Indeed, buprenorphine was administered by intraperitoneal injection at 0.1µg/g to prevent postsurgical pain, in agreement with our approved “animal experimentation procedure” (APAFIS #38705-2022092718027606 v3).

5) Line 645-647, why did the authors choose to administer the drug via the drinking water? Because it is difficult to control the exact dose in this way. What is the actual dose for mice in this study? I can’t get the accurate information from the description like this: (sm5 at 5 mg/kg and sm11 at 0,5 mg/kg).

Oral administration is a common option for administering leucovorin (aka calcium folinate/Sm11) in humans and is often well tolerated by patients, and dyclonine hydrochloride (Sm5) has been used as a local oral anesthetic in humans for more than 50 years. We privileged the oral administration via the bottle of drinking water to post-lesion adult mice to **(1) prevent stress induced inflammation as a confounding parameter** (scientific rational), **and (2) avoid unnecessary stress/suffering to post-operatory animals** as demanded by the ethical guidelines approved by our institution (ethical rational). **Importantly, as we indicated in the Fig. 7 Extended Data 1, we monitored daily the intake volume, finding no differences in the amount of drinking intake between the different treatments, thus confirming the daily oral posology of 5mg/kg of dyclonine/Sm5 and 0.5mg/kg of leucovorin/Sm11, per mouse per day**.

The previously reported dose and frequency of oral administration for leucovorin and dyclonine are variable between studies. The typical concentration of leucovorin used in humans ranges between 0.5 to 2.5 mg/kg per day. However, other studies have demonstrated the compound’s effectiveness at higher doses (2 to 10 mg/kg per day) in the context of ASD patients (Frye et al., 2018; Rossignol and Frye, 2021). We thus chose to administer leucovorin at a concentration of 0.5mg/kg per day over 7 days. For dyclonine, with two studies administering adult mice in the range of 1 to 10 mg/kg (Sahdeo et al; 2014) or at 5 mg/kg (Okazaki et al; 2018), we chose to administer the mean dose of 5 mg/kg. We used the following rationale for the treatment: considering the reported solubility in water and half-life of these compounds, we diluted them in the drinking water and renewed the treatment daily. Given that a 40 g mice drinks approximately 6mL per day, we diluted respectively 0.2mg of dyclonine/Sm5 (0.2mg in 40 g = 5 mg/kg) and 0.02mg of leucovorin/Sm11 (0.02mg in 40 g = 0.5 mg/kg) in 6mL. We then scale these concentrations to a total volume of 200mL per bottle. We added 5% glucose to the preparation to increase appetite for the mice. According to the reviewer comment, we have rephrased this part of the material and methods in **lines 817-828**.

6) It is important to set different dose levels of compounds for both in vitro and in vivo studies to demonstrate the dose or exposure/efficacy relationship to some extent. I am not sure if the authors could improve the quality of the manuscript in this aspect.

In line with the reviewer comment, we used NPC cultures to perform a systematic testing the pro-oligodendrogenic activity **dose effect *in vitro*** of the **11 selected compounds** at **3 different concentrations** (250 nM, 500 nM, and 750 nM) based on the previous reports (Mei et al., 2014; Najm et al., 2015; Eleuteri et al., 2017) and in **4 replicated experiments** (Figure 3 Extended data 1). Whole coverslip automatic quantifications of Sox10+ cells indicated that all selected compounds presented the strongest effects to promote Sox10+ oligodendroglia at 750nM. Given the results obtained in this extensive *in vitro*, we carry out following *in vitro* and *ex vivo* experiments at this optimal dose (750 nM). We have added a paragraph in the main text of make this point clearer **in lines 191-198**.

Concerning the ***in vivo*** compounds’ treatments showing robust pro-oligodendrogenic effects in two pathological models, the concentrations used were based on previously reported *in vivo* administration to mice and humans (Sahdeo et al; 2014; Okazaki et al; 2018; Frye et al., 2018; Rossignol et al., 2021).

In this context and considering the 3R ethical rule guidelines for animal experimentation, it is worth considering that in our first version of the study, we already used ∼150 adult and ∼60 postnatal animals. Nevertheless, taking into account that these experiments are laborious due to the complexity of both models of oligodendrocyte pathology, **in our revised study we addressed both the dose and the mode of administration** (see reviewer 2 point 2.3), performing a **new experiment with 30 adult animals**, and **also using** the pro-remyelinating drug **clemastine** to compare with our compounds. Considering that the doses used before where in the lower range of those previously reported (Sahdeo et al; 2014; Okazaki et al; 2018; Frye et al., 2018; Rossignol et al., 2021), **we tried oral administration by gavage** (theoretically better controlling the dose administered) at two concentrations, the same concentration (to compare with the drinking water treatment) **and a higher dose**. This corresponded to 2X-dose for dyclonine/Sm5 (10mg/kg to be at the highest range used in adult mice) and 10X-dose of leucovorin/Sm11 (5mg/kg, to be in the higher range used in humans).

After two days of gavage, we found clear signs of stress in all groups, including weight loss. Therefore, from the third day onward, we came back to the posology through the drinking water, with mice showing no more signs of stress and weight loss. Immunofluorescent analysis at 7 days post-lesion, to compare with our previous experiment, showed similar pro-oligodendrogenic effects at both doses tested (Fig. 7 extended data 1d-h), and confirmed our previous experiment (Fig. 7f-i). Moreover, while clemastine induced an increase in iOL2 density similar to dyclonine and leucovorin (Fig. 7 extended data 1h), **both leucovorin and dyclonine increased the density of proliferating OPCs** (Fig. 7m, o), **an effect that was absent for clemastine** (Fig. 7o). We have added this data in **lines 352-361**.

In the context of the hypoxia model, the intranasal administration of these compounds immediately following hypoxia is a strategy representing a noninvasive technique for drug administration, that was already demonstrated to impact brain cell behavior and eventually their proliferation (Scafidi et., 2013; Azim et al., 2017). With this treatment, we showed robust increase in the number of proliferating OPCs, progenitors fated to oligodendroglia, and an increase in the number of myelinating OLs in the leucovorin/Sm11 treatment (Fig. 6). Of course, this is without any intention to use intranasal administration in human preterm babies, which receive intravenous administration for most treatments. Our follow up project, NeoReGen, utilizes different mouse models of preterm brain injury to assess in more details the translational capacity of leucovorin and dyclonine to foster brain repair in the context of early preterm brain injury.

#### Reviewer #2 (Remarks to the Author)

Huré et al. present a pharmacogenomic approach to identify small bioactive molecules with pro-oligodendrogenic activity. Using a curation strategy (OligoScore), the identify a series of small bioactive molecules with pro-oligodendrogenic activity, based on their effect on transcriptional pathways linked to oligodendrogenesis and (re)myelination. They demonstrate that these compounds are able to induce NPC differentiation to oligodendroglial fate, boost oligodendrocyte differentiation, and promote myelination. They then selected Sm5 (dyclonine) and Sm11 (leucovorin) as candidates to move forward in preclinical models of myelin pathologies. They found that both were able to promote oligodendroglial regeneration in a mouse model of preterm brain injury and enhance oligodendrogenesis in the LPC model of de/remyelination in the corpus callosum.

### Major Points

1. While the authors do a nice job showing the effect of the compounds on several metrics in various models, the claim that the compounds **promote myelination needs to be strengthened**. The only evidence provided is the myelin index in cerebellar slice cultures.

1.a. For the in vivo models, the authors show effects on a variety of OL lineage markers. However, it would be important to look at myelination (by IHC and EM, for instance) in both the preterm brain injury model and the LPC model. For instance, reports have shown hypomyelination in the preterm brain injury model (e.g. Yuen TJ, Silbereis JC et al., Cell 2014), so an increase in myelination here with compound treatment would be interesting. In addition, in Figure 7, the title says that compound treatment promotes myelin repair, but there is no data on this to corroborate this claim.

We thank the reviewer for the comments. It is worth mentioning that we do not claim that our compounds directly increase the process of myelin formation but we now provide further evidence showing that they boost the generation of newly-formed OLs and myelinating OLs [expressing GSTπ, a marker restricted to myelinating OLs (Zhou et al., 2021)] *in vivo* in two models of oligodendrocyte/myelin pathology.

According to the reviewer’s comment, to strengthen this aspect of the study, **we performed a new experiment using 30 additional adult mice genetically labeling and quantifying the newly-formed OLs in the lesion area.** To this aim, we fluorescently labeled adult OPCs and their progeny (OPCs and OLs) by administering tamoxifen to *Pdgfra-CreER^T^; Rosa26^stop-YFP^* mice just before inducing the LPC lesion. At 10 days post-lesion, when newly-formed (re)myelinating OLs are found in the lesion area, the quantification of the number of YFP^+^ (newly-formed) oligodendroglia expressing GSTπ, a marker restricted to myelinating OLs (Zhou et al., 2021), showed that dyclonine and leucovorin increased more than 2-fold the number of newly-formed myelinating OLs (YFP^+^/GSTπ^+^ cells) in the lesion area, similar to clemastine (Fig. 8a-d). We have added this data in the result section lines 365-373.

Moreover, balancing the reviewer comment and the 3R ethical guidelines, we have also performed ultrastructural analyses of the effect of our compounds in (re)myelination in the adult model of myelin pathology, using 20 adult mice, including a group treated with clemastine as positive control. At 10 days post-lesion, we confirmed by ultrastructural analysis the increase in myelinating OLs (identified by their round- or oval-shape nucleus having densely packed chromatin and processes wrapping around myelinated axons) in animals treated with leucovorin, dyclonine, and clemastine compared to vehicle-treated animals (Fig. 8e). Moreover, quantification of myelinated axons in the lesion area showed a tendency to decrease the g-ratio of axons in leucovorin-, dyclonine-, and clemastine-treated animals compared to vehicle-treated controls, suggesting an increased remyelination (more wrapping) induced by the compound treatment, that reach significance in the case of leucovorin treatment on axons larger than 1 μm (Fig. 8f-h). We have added this in **lines 372-380**.

1.b. The in vivo evidence of compounds boosting OL differentiation and maturation could be stronger. For instance, Fig 6I/J show that Hx + compound is not significantly different from Nx, but only leucovorin treatment is significantly boosting OL differentiation / maturation over the Hx condition, which seems to be the more relevant comparison here.

To answer the reviewer point, in the context of the **neonatal hypoxia model**, **we now provided more complete analyses**, increasing the number of animals, and performing automatic and manual quantifications in different brain regions, now provided in Figure 6 and Fig.6 extended data 1f. and detailed in **lines 290-320**.

1.c. Figure 4 – it would be helpful to also look at number of Plp+ (or another mature OL marker, like MBP) to see if these compounds affect numbers of differentiated cells, versus an effect on morphology, which the authors report.

According to the reviewer comment, we now present automatic quantifications of total cells in OPC differentiation cultures after three days of compound’s treatment (Figure 4 extended data 1), showing that, while no significant changes were seen in the variable density of PDGFRα+ OPCs (Fig. 4b, c), except for Sm7, both T3, clemastine, and all tested compounds (Sm1, Sm2, Sm5, Sm6, and Sm11) increased the number of MBP+ OLs compared to vehicle-treated controls (Fig. 4b, d). These results demonstrate the capacity of most selected compounds to cell-autonomously foster OPC differentiation. We have included this in **lines 226-231**.

2.1 The compound dosing paradigm for in vivo studies need corroboration. For instance, in the LPC studies, compounds were dosed in the drinking water, so there is no controlled way to monitor the amount of drug being delivered.

Oral administration is a common option for administering leucovorin (aka calcium folinate/Sm11) in humans and is often well tolerated by patients, and dyclonine hydrochloride (Sm5) has been used as a local oral anesthetic in humans for more than 50 years. We privileged the oral administration via the bottle of drinking water to post-lesion adult mice to **(1) prevent stress induced inflammation as a confounding parameter** (scientific rational), **and (2) avoid unnecessary stress/suffering to post-operatory animals** as demanded by the ethical guidelines approved by our institution (ethical rational). **Importantly, as we indicated in the Fig. 7 Extended Data 1, we monitored daily the intake volume, finding no differences in the amount of drinking intake between the different treatments, thus confirming the daily oral posology of 5mg/kg of dyclonine/Sm5 and 0.5mg/kg of leucovorin/Sm11, per mouse per day**.

The previously reported dose and frequency of oral administration for leucovorin and dyclonine are variable between studies. The typical concentration of leucovorin used in humans ranges between 0.5 to 2.5 mg/kg per day. However, other studies have demonstrated the compound’s effectiveness at higher doses (2 to 10 mg/kg per day) in the context of ASD patients (Frye et al., 2018; Rossignol and Frye, 2021). We thus chose to administer leucovorin at a concentration of 0.5mg/kg per day over 7 days. For dyclonine, with two studies administering adult mice in the range of 1 to 10 mg/kg (Sahdeo et al; 2014) or at 5 mg/kg (Okazaki et al; 2018), we chose to administer the mean dose of 5 mg/kg. We used the following rationale for the treatment: considering the reported solubility in water and half-life of these compounds, we diluted them in the drinking water and renewed the treatment daily. Given that a 40 g mice drinks approximately 6mL per day, we diluted respectively 0.2mg of dyclonine/Sm5 (0.2mg in 40 g = 5 mg/kg) and 0.02mg of leucovorin/Sm11 (0.02mg in 40 g = 0.5 mg/kg) in 6mL. We then scale these concentrations to a total volume of 200mL per bottle. We added 5% glucose to the preparation to increase appetite for the mice. According to the reviewer comment, we have rephrased this part of the material and methods in **lines 817-828**.

2.2 Were any measures of PK and/or target engagement done?

As mentioned in the introduction, results, and discussion, our drugs/compounds have been selected by their large pro-oligodendrogenic activity involving different cellular (neural progenitor/stem cells, OPCs, and microglia) and molecular target mechanisms (including the one carbon/ folic acid metabolism in the case of leucovorin/sm11 and NRF2 antioxidant activities in the case of dyclonine/sm5). Furthermore, for the *in vivo* models of myelin pathologies, we selected FDA-approved compounds previously used in human and animal models. Therefore, it is clearly beyond the scope of this large and extensive study to perform further detailed measures of pharmacokinetics in animal models.

2.3 This would help strengthen things. Could the authors try PO dosing?

Concerning the ***in vivo*** compounds’ treatments showing robust pro-oligodendrogenic effects in two pathological models, the concentrations used were based on previously reported *in vivo* administration to mice and humans (Sahdeo et al; 2014; Okazaki et al; 2018; Frye et al., 2018; Rossignol et al., 2021).

In this context and considering the 3R ethical rule guidelines for animal experimentation, it is worth considering that in our first version of the study, we already used ∼150 adult and ∼60 postnatal animals. Nevertheless, taking into account that these experiments are laborious due to the complexity of both models of oligodendrocyte pathology, **in our revised study we addressed both the dose and the mode of administration** (see reviewer 2 point 2.3), performing a **new experiment with 30 adult animals**, and **also using** the pro-remyelinating drug **clemastine** to compare with our compounds. Considering that the doses used before where in the lower range of those previously reported (Sahdeo et al; 2014; Okazaki et al; 2018; Frye et al., 2018; Rossignol et al., 2021), **we tried oral administration by gavage** (theoretically better controlling the dose administered) at two concentrations, the same concentration (to compare with the drinking water treatment) **and a higher dose**. This corresponded to 2X-dose for dyclonine/Sm5 (10mg/kg to be at the highest range used in adult mice) and 10X-dose of leucovorin/Sm11 (5mg/kg, to be in the higher range used in humans).

After two days of gavage, we found clear signs of stress in all groups, including weight loss. Therefore, from the third day onward, we came back to the posology through the drinking water, with mice showing no more signs of stress and weight loss. Immunofluorescent analysis at 7 days post-lesion, to compare with our previous experiment, showed similar pro-oligodendrogenic effects at both doses tested (Fig. 7 extended data 1d-h), and confirmed our previous experiment (Fig. 7f-i). Moreover, while clemastine induced an increase in iOL2 density similar to dyclonine and leucovorin (Fig. 7 extended data 1h), **both leucovorin and dyclonine increased the density of proliferating OPCs** (Fig. 7m, o), **an effect that was absent for clemastine** (Fig. 7o). We have added this data in **lines 352-361**.

In the context of the hypoxia model, the intranasal administration of these compounds immediately following hypoxia is a strategy representing a noninvasive technique for drug administration, that was already demonstrated to impact brain cell behavior and eventually their proliferation (Scafidi et., 2013; Azim et al., 2017). With this treatment, we showed robust increase in the number of proliferating OPCs, progenitors fated to oligodendroglia, and an increase in the number of myelinating OLs in the leucovorin/Sm11 treatment (Fig. 6). Of course, this is without any intention to use intranasal administration in human preterm babies, which receive intravenous administration for most treatments. Our follow up project, NeoReGen, utilizes different mouse models of preterm brain injury to assess in more detail the translational capacity of leucovorin and dyclonine to foster brain repair in the context of early preterm brain injury.

3. The link to microglia, particularly the change in phagocytic / pro-regenerative microglial numbers, is intriguing but also needs to be strengthened. Did the authors look at the presence of myelin debris (e.g. dMBP staining)? Were changes in inflammatory signals observed?

In agreement to the reviewer points, to complement our study of microglial inflammatory, phagocytic and regenerative profiles, we now provide the quantification for myelin debris using dMBP staining (Shen et al., 2021) at 7 days post-lesion (dpl) showing that both leucovorin/Sm11 and dyclonine/Sm5 treatments reduce the amount of myelin debris in the lesion area and increase the proportion of dMBP phagocytosed by CD68^+^ microglia (Fig. 9a-e). With respect to changes in inflammatory microglial profiles, using iNOS and Cox2 as pro-inflammatory markers, we now show that both leucovorin- and dyclonine-treatments strongly reduce the inflammatory microglial profile at 7 dpl (Fig. 9f-h), in parallel to the increase in Arg1^+^ pro-regenerative microglia profile (Fig. 9i, j). Altogether these results strongly support that leucovorin and dyclonine promote/accelerate the lesion repair also by inducing microglia beneficial activities, including a faster clearance of myelin debris (dMBP by CD68^+^ cells) and an earlier transition microglial profiles from inflammatory (iNOS^+^/Cox2^+^) towards pro-regenerative (Arg1^+^). We have now added this data in Figure 9 and **lines 394-403**.

In Fig 7O, it also appears that lesion size might vary. Was this examined?

According to the reviewer comment, we reconstructed the lesion volume at 10 dpI to assess the impact of compounds’ treatment on the lesion size finding no significant reduction of the lesion volume upon clemastine-, dyclonine-, and leucovorin-treatment, likely due to lesion variability. We now provide this data in Fig. 8c and in the main in **line 371**.

4. The bioinformatics data analysis needs clarification:

4a. The authors mention that raw sequencing data was downloaded from GEO, but it appears that the authors applied a microarray data processing pipeline. Can this be clarified?

According to the reviewer comment, we now clarify this in the methods in lines 559-565. We also now provide the bioinformatic analyses of the expression microarrays deposited in GEO of our previous publications (Azim et al., 2015; Azim et al., 2017) and detailed below.

4b. In addition, no details were documented about how the authors applied Seurat for single-cell RNAseq data analysis and for bulk RNAseq analysis. For instance, how did the authors correct for different batches for data integration? How many cells were integrated? What statistical method was used for differential analysis? Did the authors use adjusted p-value or a raw p-value as cutoff?

To obtain an oligodendroglial transcriptomic signature, we used expression data already processed by the respective authors from different sorts of datasets (i.e., expression microarrays, bulk-RNA-seq, and single cell RNA-seq datasets; see methods corresponding to all cited papers). For most datasets, the supplementary tables provided were used to select genes with enriched expression in each cell type (p-value < 0.05) or differentially expressed genes between different cell-subtypes/stages/clusters (>= 1.5-fold change compared to other cell types). As mentioned above, Methods table 2 summarizes all this information. We now also provide the script for the microarray analyses (dNSCs_dTAPs_Azim et al.R and OLglia_Zhang et al.R) and the provide a more detailed description in the Methods section *Data processing and oligodendroglial gene enrichment* in **lines 555-601.**

### Minor Points

1. The sentence in the abstract “An unmet challenge is the discovery of medications presenting convincing pro-oligodendrogenic activity for the eventual clinical treatment of these pathologies” is somewhat misleading as there are several compounds currently being tested (e.g. clemastine, analogs of thyroid hormone) that promote OPC differentiation and remyelination. Of note, this is nicely discussed in the introduction.

According to the reviewer suggestion we have modified this sentence of the abstract to: [‘No medication presenting convincing repair capacity in humans has been approved for these pathologies’]

2. Figure 3 – Why was 750nM chosen as the dose to test all compounds (also 250 and 500nM in Figure 3, extended data 1)?

As mentioned in our answer to reviewer#1 question 6, we used NPC cultures to perform a systematic *in vitro* testing the pro-oligodendrogenic activity **dose effect** of our **11 selected compounds** at **3 different concentrations** (250 nM, 500 nM, and 750 nM) based on the previous reports (Mei et al., 2014; Najm et al., 2015; Eleuteri et al., 2017) and in **4 replicated experiments** (Figure 3 Extended data 1). Whole coverslip automatic quantifications of Sox10+ cells indicated that all selected compounds presented the strongest effects to promote Sox10+ oligodendroglia at 750nM. Given the results obtained in this extensive *in vitro* experiment, we carry out following *in vitro* and *ex vivo* experiments at this optimal dose (750 nM). We have made this point clearer in **lines 191-197**.

3. Figure 5 – CaBP staining is in green, but the CaBP+ axons are shown in white?

The overlap between CaBP^+^ axons labeled **in green** and MBP^+^ myelin labeled **in magenta,** gives the overlapping white color of ‘myelinated axons’ quantified in the myelination index. We now provided higher magnification pictures illustrating this point in Fig. 5 extended data 1b.

4. For the slice cultures, it would be helpful to have another metric of myelination, perhaps Caspr staining to axons, which would mark compact myelin and not be confounded by MBP+ oligodendrocytes on axons.

In many papers (including Yuen TJ, Silbereis JC et al., Cell 2014 cited by the reviewer and the method book chapter entitled *Investigating demyelination, efficient remyelination and remyelination failure in organotypic cerebellar slice cultures*, Gorter et al., 2022; DOI: 10.1016/bs.mcb.2021.12.011) myelination in explant cultures is quantified by the overlap between the axonal and the myelin segments. We now also provide higher magnification pictures making this more visible in Figure 5 extended data 1b, c). It would be very challenging to optimize a whole explant automatic quantification of Caspr^+^ paranodes (as we do in our experiments for all the markers used), and will not reinforce the messages of the paper. Therefore, given that it is mentioned by the reviewer as a minor point, we have concentrated our efforts and limited the number of animals sacrificed to address the major points. Note that we have now provided immunofluorescence and electron microscopy data showing that at 10 days post-lesion, both leucovorin- and dyclonine-treatment increased the number of newly-formed myelinating oligodendrocytes in the lesion area and, that leucovorin-treatment significantly increased the myelin thickness of axons larger than 1 μm (Figure 8).

It is intriguing that clemastine promotes OL differentiation, but not myelination, while T3 does not promote differentiation, but promotes myelination.

It is noteworthy that both T3 and clemastine treatments increase the differentiation and myelination indexes (Figure 5c, e), even though the experimental variability makes that these differences reach significance in one or the other case. Indeed, as shown in the graphic below, g-test between T3 and clemastine indicates that the differences between their sample values are non-significantly different (n.s.) in either differentiation index (Mann-Whitney test; p-value = 0.1893) or myelination index (Mann-Whitney test; p-value = 0.0907).

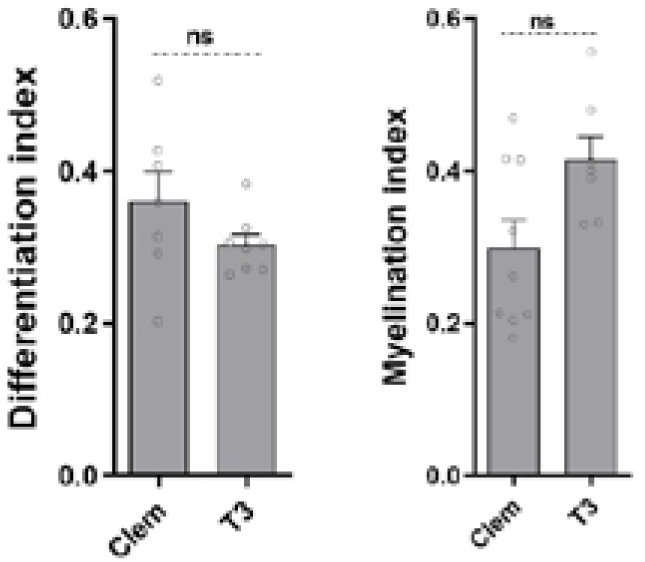

5. Discussion on why Sm2 was not selected to move forward to in vivo studies would be helpful. Is it simply that it was ranked #3 (as shown in Figure 5, extended data 1), or was there additional rationale?

Sm2/Heptaminol is myocardial stimulant and vasodilator, thus having a risk of side effects in the context of preterm brain injury, what made us not select it in the top two compounds. Its applications in the context of adult MS lesion repair will require future *in vivo* animal studies.

6. The manuscript could benefit from proofreading throughout (some grammar errors and typos).

We thank the reviewer for his/her comments, and we have carefully paid attention to this aspect in our revised manuscript checked by English native-speakers.

#### Reviewer #3 (Remarks to the Author)

**This manuscript tackles an important aspect of debilitating diseases** such as newborn brain injury and multiple sclerosis, that of identifying therapeutic agents which promote remyelination in injury. Several other studies have used cell screening approaches in vitro and in vivo to identify compounds which promote remyelination. **The novelty of this paper lies in the demonstration of an informatics approach to predict compounds** which will induce oligodendrogenesis. The authors use this approach to identify predicted compounds which will promote remyelination after injury, and use two of them in vivo to promote injury repair. **I like the idea of this novel informatics approach a lot**, but have quite a few issues with its execution:

We thank the reviewer for the enthusiasm in the clinical relevance of our study and the novelty of our pharmacogenomic approach. We answer below to the questions/points raised.

1) It seems unclear from the description **how SPIED is predicting bioactive molecules** which induce transcriptional changes that induce oligodendrogenesis. A more thorough description of how this tool is used would be useful.

Following the suggestion of the reviewer, we have added a paragraph to the Methods section entitled *Pharmacogenomics using SPIED platform* in **lines 603-613**.

2) The main novelty of this paper is the informatics approach used. The authors discuss different gene sets for different processes of oligodendrocyte biology, specification, proliferation, migration, survival, differentiation, myelination. But it seems as if they lump all processes together to identify drugs that are ‘oligodendrogenic’?

As indicated in the introduction, we chose ‘**a more comprehensive strategy**’ to find **drugs acting on several processes of oligodendrogenesis**, thus ranking and selecting drugs by the total score (sum of individual process’ scores) which explain our results showing that dyclonine/Sm5 and leucovorin/Sm11 can act in different cell types and promote different oligodendroglial processes: (1) NPC differentiation into oligodendroglia, (2) OPC proliferation, (3) OPC differentiation, and (4) accelerates microglial phagocytosis and its faster transition from pro-inflammatory to pro-regenerative microglial profiles. We have state this aspect more clearly as follows:

##### Introduction lines 91-97

[‘While most previous studies have followed a gene/pathway candidate approach, here **we aimed at using a more comprehensive strategy** (Fig. 1 extended data 1). To this end, we developed an *in-silico* approach combining both unbiased identification (i.e., transcriptomes) and knowledge-driven curation of transcriptional changes associated with oligodendrogenesis (i.e., OligoScore). We then identified small bioactive molecules (compounds) **capable of globally mimicking this transcriptional signature.**’],

We have now made this point clearer in the abstract and first paragraph of the discussion, as follows:

##### Abstract

[‘Here, we present a pharmacogenomic approach leading to the identification of small bioactive molecules with a **large pro-oligodendrogenic activity**, selected through an expert curation scoring strategy (OligoScore) **of their large impact on transcriptional programs controlling oligodendrogenesis and (re)myelination**]

##### Discussion lines 413-420

[‘While most previous studies have followed a gene/pathway candidate approach, **here we used a more global strategy to fill this gap**. Leveraging transcriptomic datasets through a pharmacogenomics analysis and developing an expert curation of genes previously involved in oligodendroglial biology (provided as a resource for the scientific community: [OligoScore](https://oligoscore-staging.icm-institute.org/), https://oligoscore.icm-institute.org), we identified and ranked novel small bioactive molecules (compounds) **fostering transcriptional programs associated with various aspects of oligodendrogenesis**, including OPC proliferation, differentiation, and (re)myelination.’]

Were compounds assessed that effect specific individual processes?

Yes. Figure 2B illustrates the genes in each process of oligodendrogenesis used to query SPIED, obtaining a total of 393 different compounds, with 156 of them overlapping with those obtained using the oligodendroglial transcription signature (Figure 2c). For these 156 selected compounds, we now provide the processes identifying each compound (**Supplemental table 7, column C**). It is worth noting that, given that each compound is scored by their curated genes involved in each process of oligodendrogenesis (columns G to M), most of the compounds on the upper part of the list (with higher total scores) were identified using the curated genes of several (2 or 3) oligodendrogenic processes (as indicated in column C). As mentioned above, in our study we focused on compounds having a large pro-oligodendrogenic activity (top of the list), and indeed, our experimental data validates this strategy given that using different experimental models, we demonstrate the capacity of Sm5/Dyclonine and Sm11/Leucovorin to increase oligodendroglial specification (Fig. 3c, g and Fig. 6f), OPC proliferation (Fig 6d, 7m,n), OPC differentiation (Fig. 4d, 5c, 7h,i), and the number of myelinating OLs (Fig. 5e and Fig. 8).

3) A somewhat similar point, of the compounds selected, which transcriptional hubs are activated by each and which individual processes are predicted to be affected by this?

Even though these questions beg for follow up studies, according to the reviewer request, we now provide the list of 252 up- and down-regulated transcriptional hubs associated with the genes regulated by dyclonine/Sm5 (**Supplemental table 9**) and leucovorin/Sm11 (**Supplemental table 10**), together with the enriched biological process (Gene Ontology) associated with them. It is worth noting that the biological processes associated with correlative gene hubs are mainly related to oligodendroglial development, while those associated with anti-correlative genes hubs are mainly related to immune and inflammation processes. We also provide now the OligoScore analysis of correlative hub genes regulated by dyclonine/Sm5 (**Supplemental table 11**) and leucovorin/Sm11 (**Supplemental table 12**) indicating the processes of oligodendrogenesis affected by the curated hub genes (i.e., those with already known function in oligodendrogenesis).

4) In both in vivo hypoxia and lysolecithin models, the chosen compounds are described as increasing OPC proliferation as well as the number of differentiated OL. It is difficult to understand how a drug is affecting both of these two very different processes. It is important to describe gene hubs affected by the drugs and how this leads to effects on proliferation and differentiation.

According to the reviewer’s suggestion, we investigated how the hub genes regulated by Sm5 (Supplemental table 11) and Sm11 (Supplemental table 12) influence the processes of proliferation and differentiation by querying OligoScore for the known activities of these genes. We found 6 genes promoting proliferation, with 5 of them commonly regulated by Sm5 and Sm11 (*Cntnn1, Olig2, Ascl1, Myt1,* and *Sox2)* and many genes promoting differentiation (18 for Sm11, and 14 for Sm5; see Supplemental tables 11 and 12). Interestingly, four of these genes encode for key transcription factors (*Olig2, Ascl1, Sox2,* and *Myt1*) promoting both proliferation and differentiation programs (see references associated to each gene and process in OligoScore, thus allowing a balance between both cell fates (proliferation and differentiation) in OPCs. These results explain, at least in part, how Sm5 and Sm11 promote parallel OPC proliferation and differentiation. We discuss this point in the discussion section lines 431-436.

It is also worth mentioning that the current evidence based in long term *in vivo* live-imaging of adult OPCs in physiological or demyelinating conditions, suggests that OPCs start differentiation without previous division, with the ‘*hole’* left by their differentiation being ‘*filled’* by the proliferation of neighboring OPCs (Hughes et al., 2013, DOI: 0.1038/nn.3390). This is also supported by single cell transcriptomics showing different subsets (/clusters) of OPCs (see Marques et al., 2016, 2018; Spitzer et al., 2018, DOI: 10.1016/j.neuron.2018.12.020). Therefore, it is conceivable that compounds fostering the metabolism of OPCs, like leucovorin promoting the one-carbon metabolism, can on the one hand promote the differentiation those OPCs in a state prompt to differentiate, while on the other hand, induce OPCs in different state to proliferate more actively.

5) The most confusing part of the manuscript comes at the very end. The point of the paper is to identify compounds which affect oligodendrocyte transcriptional networks, but then at the very end the authors seem to be suggesting that the compounds they chose also alter a microglial signature in vivo in injury, raising the possibility that the promotion in remyelination has nothing to do with the effects on OPCs but due to altered microglia.

As stated above, our strategy aimed to select compounds having a large pro-oligodendrogenic activity, based on their regulatory impact on several transcriptional programs. It is thus not surprising that the selected compounds, particularly sm5/dyclonine and sm11/leucovorin studied in more detail *in vivo*, can also impact other cell-types than oligodendroglia, including neural progenitor cells (NPCs) and microglia. During our study we demonstrated: (1) using neonatal neural progenitor cells’ cultures (thus in the absence of microglia), the capacity of all selected compounds (Sm1 to Sm11) to foster differentiation of (NPCs) into oligodendroglia (figure 3); (2) using purified OPC cultures (absence of microglia), the direct capacity of Sm5 and Sm11 to promote OPC differentiation (figure 4); and (3) the *in vivo* capacity of Sm5 & Sm11 to foster neonatal NPC differentiation into oligodendroglia (figure 6, a process never described before to involve microglia). Therefore, these results demonstrate a direct activity of Sm5 and Sm11 in NPCs, OPCs, and likely differentiating OLs to promote oligodendrogenesis. Thus, the fact that Sm5 and Sm11 have effects in other cell types, such as microglia, is also not unexpected given that we selected the also for their: (a) previous reports of their anti-inflammatory activities (Rufini et al., 2022; Cianciulli et al., 2016; Tommy et al., 2021), (b) some of their pharmacogenomics’ target genes (e.g., Sm11/leucovorin downregulates *IFNG,* the gene coding for the inflammatory cytokine Interferon gamma), and (c) their abovementioned anti-correlative genes hubs are mainly related to immune and inflammation processes. In this line of argument, the anti-inflammatory effect of Sm11 is discussed in **lines 510-517**.

## Point by point response to reviewers’ comments (2^nd^ round)

### Response in blue

We would like to express our gratitude for the valuable feedback provided by the reviewers. With their insightful comments and our broader analyses, our results more strongly support and further extend the pre-clinical aims of the study, i.e., demonstrate the pro-oligodendrogenic and anti-inflammatory activities of leucovorin and dyclonine, two FDA-approved compounds, leading to brain repair in two models of oligodendrocyte pathologies, thus significantly widening its potential for future clinical impact.

### Reviewers’ comments

#### Reviewer #2 (Remarks to the Author)

Thank you to the authors for their revision and additional experiments. While the authors conducted numerous additional experiments and useful discussion, which help strengthen their claims, there are still points that are central to their claims that need to be strengthened.

1) The authors main aim is to identify compounds that promote repair that can translate to pathologies such as early preterm-birth brain injury (PBI) and adult multiple sclerosis (MS). They have done extensive work to show the enhancement of proliferation and oligodendroglial fate, as well as differentiation. However, as the ultimate aim is to promote myelination or remyelination, this still needs to be strengthened.

a. The fate mapping experiments (Figure 8) add to the story, but still do not address the myelination component. For instance, there could be many mature oligodendrocytes, which does not translate into myelin formation. The claim that these are (re)myelinating oligodendrocytes is not supported with data.

We would like to express our disagreement with the reviewer’s statement. Following previous suggestions from the reviewer, to strengthen this aspect of the study, we performed two relevant and demanding experiments: (1) a new experiment using 30 additional adult mice to genetically label and quantify the newly formed OLs in the lesion area, using the pro-myelinating compound clemastine as positive control; (2) an experiment using 20 adult mice to perform EM analysis (see next point).

In our immunofluorescent analysis, we used GSTp, that together with ASPA (Pan et al., 2020; https://doi.org/10.1038/s41593-019-0582-1), is a marker labeling the cytoplasm of mature/myelinating oligodendrocytes, thus allowing quantification of OL numbers, as previously demonstrated by several studies (Zhou et al., 2021; https://doi.org/10.7554/eLife.60467; Hassel et al., 2023; https://doi.org/10.1002/glia.24373; Tansey & Cammer, 1991; https://doi.org/10.1111/j.1471-4159.1991.tb02104). Moreover, we provided direct evidence that GSTp^+^ cells are indeed myelinating OLs. First, in the context of the postnatal brain, we show that GSTp^+^ cells have the typical morphology of myelinating OLs visualized by the GFP reporter-labeling in myelin segments/internodes (Fig. 6 extended data 1e). Second, in the context of the adult de/remyelination model, where we genetically labeled OLs generated from adult OPCs (by administrating tamoxifen before lesion induction to *Pdgfra-CreERT/Rosa26^flox-stop-flox-YFP^* mice), we used GSTp^+^ to immunodetect myelinating OLs, showing that many GSTp^+^ cells presented the typical alignment of myelinating OLs along axons in the corpus callosum (Fig. 8d). Moreover, at the same time, we used Bcas1 to immunodetect pre-myelinating/immature OLs (Fard et al., 2017), finding no overlap between Bcas1^+^/YFP^+^ cells, presenting a large cytoplasm and long processes typical of pre-myelinating/immature OLs (see Fig 8d, high mag panels), and GSTp^+^/YFP^+^ cells, having a small rounded cytoplasm typical of myelinating OLs (Fig. 8d), strongly supporting that GSTp^+^/YFP^+^ cells are newly-formed (YFP^+^ cells) myelinating OLs.

Finally, we would like to highlight that to the best of our knowledge, we did not find papers providing evidence of the existence of mature non-myelinating OLs in the LPC or cuprizone de/remyelinating murine models. Indeed, the heterogeneity of mature OLs shown by single-cell transcriptomic analyses in adult mice (Marques et al., 2016) and in the brain of human MS patients and healthy controls (Jakel et al., 2019) has not been shown by follow-up studies to correspond to non-myelinating oligodendrocytes. For example, time-lapse analysis in the cuprizone model, using transgenic mice reporting with fluorescence OL myelin segments, identified regenerated oligodendrocytes in the lesion area by the mature form reached 12– 14 days after the first appearance, all of them being myelinating oligodendrocytes (Orthmann-Murphy et al., 2020, DOI: 10.7554/eLife.56621). Also, in a recent study performing combination between spatial transcriptomics and electronic microscopy analysis in the LPC de/remyelinating model (Androvic et al., 2023, DOI: 10.1038/s41467-023-39447-9), the authors identified four mature OL subtypes/clusters (Oligo 1-4), with Oligo 3 and 4 clusters representing injury-responding states of mature oligodendrocytes, with Oligo 3 OLs expressing marker genes of previously defined disease-associated OLs in aging, AD and MS models, and Oligo 4 OLs expressing a battery of interferon-stimulated genes, and thus called interferon responsive OLs. It is worth noting that the study did not provide any evidence or suggestion that one of these mature OL subtypes corresponded to mature non-myelinating OLs.

To make this point clearer in lines 367-373, we have now expressed it as follows: [‘and at 10 dpl identified by immunofluorescence newly formed OLs (YFP^+^ cells) being either Bcas1, a marker of immature/pre-myelinating OLs (Fard et al., 2017), or GSTπ, a marker restricted to myelinating OLs (Zhou et al., 2021). Interestingly, we did not find overlap between YFP^+^/Bcas1^+^ cells and YFP^+^/GSTπ^+^ cells, suggesting that Bcas1 and GSTπ indeed identify immature/pre-myelinating and mature/myelinating OLs, respectively.’]

Tansey FA & Cammer W. A pi form of glutathione-S-transferase is a myelin- and oligodendrocyte-associated enzyme in mouse brain. JNeurochem 1991. DOI: 10.1111/j.1471-4159.1991.tb02104

Pan, …, Jonah R. Chan & Mazen A. Kheirbek. Preservation of a remote fear memory requires new myelin formation. Nat Neurosci. 10 Feb 2020. DOI: 10.1038/s41593-019-0582-1

b. Did the authors examine a quantitative or semi-quantitative measure of remyelination in their EM study? G-ratios likely are not a very useful metric in the corpus callosum.

As mentioned above, we used 20 additional adult animals to perform the EM analysis, showing that myelinating OLs in the lesion area, identified by their nucleus of round- or oval-shape having densely packed chromatin and processes wrapping around myelinated axons, were more abundant (**semi-quantitative measure**) in animals treated with leucovorin, dyclonine, and clemastine compared to vehicle-treated animals, as illustrated in Fig. 8e, paralleling the results quantified using immunodetection of GSTp^+^/YFP^+^ OLs (Fig. 8b, d). We now added a supplemental figure (**Figure 8 extended data 1**) with higher magnification micrographs illustrating newly formed (re)myelinating OLs (mOLs) in the lesion area, identified by their typical ultrastructural traits (i.e., oval-shape nucleus having densely packed chromatin with several heterochromatin spots and large cytoplasm in continuity with axons presenting compact myelin ultrastructure, red arrows) in animals treated with our compounds.

We also used EM to quantify myelinated axons in the lesion area (identified by cellular and extracellular disrupted structures; see Fig. 8e) finding that almost all axons were (re)myelinated, as illustrated in panels of Fig. 8f. Despite our early timing of analysis for the remyelination of the lesion (10 days post-lesion), we could detect a tendency of increased myelin thickness (decrease in g-ratios; Fig. 8g; **semi-quantitative measure**) in axons from all compound-treated animals compared to vehicle-treated. Moreover, we found a statistical increase in myelin thickness (lower g-ratio) of axons with a diameter larger than 1mm in leucovorin-treated animals, compared to the vehicle (Fig. 8h; **quantitative measure**). These results could be interpreted as compounds accelerating the process of lesion remyelination. Nevertheless, we only state: [‘suggesting an increased remyelination (more wrapping) induced by the compound treatment’ […] ‘Altogether, these results show that, similar to clemastine, leucovorin and dyclonine promote OL differentiation and remyelination *in vivo* in the context of adult brain demyelination, with leucovorin showing the strongest effect’.]

To be even more careful in our conclusions, we propose to rephrase this in lines 387-388 as follows: [‘promote OL differentiation and **thus** remyelination. **The possibility of leucovorin and dyclonine directly inducing myelin formation would require further investigation**.’]

c. Figure 6f, h, j – was the difference between Hx vs. Hx + Dycl. statistically significant? It is not shown, but the text is inconsistent in the results (mentioning a rescue and then more mitigated effects).

Both Fig. 6h and 6j show that the statistical reduction in CC1^+^ cells and GSTp^+^ cells between Hx vs. Nx is ‘rescued’ (non-significant difference with normoxic group, n.s.) in hypoxic animals treated with dyclonine and leucovorin. Only in the case of leucovorin, the data reach a statistical difference between Hx *vs.* Hx + Leuc, both in Fig. 6j, 6h, and 6l, as also shown in the heatmap representations depicting the automatic quantifications of the density of GSTp^+^ cells (myelinating OLs) in different brain regions (Fig. 6k).

According to the reviewer’s comment we propose to change the text in main lines 293-295 as follows:

[‘their differentiation into CC1^+^/Olig2^+^ OLs was impaired by hypoxia but **this difference with the normoxic group was** rescued by both dyclonine and leucovorin treatments, **with leucovorin also reaching statistical difference with the hypoxic group**’]. In lines 298-299 [‘Quantification of Olig2^+^/GSTπ^+^ cell density revealed a marked effect of hypoxia on OL maturation that was fully rescued by leucovorin treatment.’]

We have also changed the figure legend of Fig. 6 accordingly: [‘(h) showing that dyclonine and leucovorin rescue the **reduced density of differentiating OLs** (CC1^+^/Olig2^+^ cells) **induced by** neonatal chronic hypoxia within the cortex at P19 to **the density found in the normoxic group, with leucovorin reaching statistical difference with the hypoxic group**’.]

It is worth noting that in the section named ‘limitations of the study’, we already discuss this as follows: [‘the increase in OPC proliferation and numbers mediated by dyclonine in the hypoxia model do not rescue the density of myelinating OLs, contrasting with its effects in adult brain de/remyelination, calling for a better understanding of the environmental differences of each pathological model, such as microglial involvement.]

d. The authors conclude that the compounds have the capacity to cross the BBB, but no data are shown to directly measure compound concentration in the brain to support this conclusion?

It should be considered that we selected leucovorin and dyclonine given that they were FDA-approved compounds with previous publications demonstrating the capacity to cross the BBB. Indeed, for leucovorin/Sm11, the crossing of the BBB is supported by several publications both in animal models and in humans, as mentioned in the discussion section lines 499-505: [‘Both leucovorin and folic acid can be transformed by different enzymes into 5-methyl-tetrahydro-folate, which efficiently crosses the BBB using the folate receptor alpha (FRα) transporter, and is thought to be the main active folate in the CNS (Scaglione and Panzavolta, 2014). Leucovorin has the advantage over folic acid of using other transporters present in the choroid plexus to get into the CNS, i.e., the proton-coupled folate transporter (PCFT) and the reduced folate carrier (RFC) (Mafi et al., 2020).’ (…) leucovorin has been used to treat epileptic patients having mutations in the folate receptor alpha gene, FOLR1 (Mafi et al., 2020)’]. We also give references supporting it in Table S8 column F, and in the file called *references BBB* (S8-6, S8-1, S8-2). For dyclonine/Sm5, we mention in the discussion lines 484-487 that [‘In a study of drug repositioning in the context of Friedreich Ataxia, dyclonine was found to confer protection against diamide-induced oxidative stress through binding to the transcription factor NRF2’].

To make this point clearer, we have now added the original reference of this study (Sahdeo et al., 2014; doi: 10.1093/hmg/ddu408), already cited in the methods section, in which mice and human patients are treated with oral administration of dyclonine showing improvement from symptoms associated with Friederich Ataxia. We also provide the original reference of crossing the BBB in S8-21 in the file *references BBB*.

Finally, it is beyond the scope of this large screening study to generate data providing compound concentration in the mouse brain, which should be done in follow-up studies.

2) Thank you for adding Methods Table 2 to summarize the expression datasets included in this study and the statistics for selecting DEGs. However, the single-cell data processing documented in lines 597-600 is still confusing. How did the authors apply both ‘min genes=500 and max genes<200’ filtering conditions simultaneously? Why are the authors interested in cells with high mitochondrial percentage (>10%)?

a. The following link may help with general guidance: https://satijalab.org/seurat/articles/pbmc3k_tutorial.html

b. I assume the authors used the first 20 PC dimensions for UMAP or tSNE clustering with a clustering reduction of 0.9? For the 1,598 DEGs identified, which statistical method was used here?

(using the following criteria: min.cells = 10; min.genes = 500; number of UMI >5100; max genes <200; mitochondrial percentage >0.10, analysis in 20 dimensions with a resolution of 0.9)

We thank the reviewer for spotting this mistake. We have corrected this paragraph in lines 602-612 as follows: [‘using Marques and colleagues’ (2016) supplemental table ‘aaf6463 Table S1’, sheet ‘specific genes’ we pooled all genes except those of VLMC column obtaining 532 unique genes. To extract more information, raw data from 5072 cells were processed with the Seurat package v2 (Stuart et al., 2019) using the following criteria: min.cells = 10; min.genes = 500; mitochondrial percentage <0.10, 20 dimensions to find neighbors, a resolution of 0.9 to find clusters, selecting the top150 genes from each cluster from FindAllMarkers function, identifying 1598 DEGs (Methods Table 2).’]. We have modified accordingly the text in Methods Table 2 and provide the R script (‘Marques 2016 gene selection.R’).

#### Reviewer #3 (Remarks to the Author)

I appreciate the authors responses to my questions, and feel now they have addressed all the concerns.

We thank the reviewer for the useful comments that have allowed us to improve the quality of the study.

